# Retuning ventral hippocampal output improves social memory and reduces anxiety-related behavior in temporal lobe epilepsy

**DOI:** 10.1101/2025.09.12.675815

**Authors:** Anthoni Goodman, Emily Schellinger, Julia Kahn, Hajime Takano, Douglas Coulter, Srdjan Joksimovic

## Abstract

Temporal lobe epilepsy (TLE) is associated with cognitive, social, and emotional comorbidities that persist between seizures and remain poorly addressed by seizure-focused treatment. We examined whether these interictal deficits reflect altered ventral CA1/subiculum (vCA1/Sub) output and whether shifting principal-cell excitability can improve behavior despite persistent pathology. In pilocarpine-treated mice, vCA1/Sub principal neurons were depolarized and showed altered recruitment, alveus/oriens stimulation produced diminished hyperpolarizing responses, and parvalbumin- and somatostatin-immunopositive cell densities were reduced in stratum oriens. Epileptic mice did not demonstrate social discrimination above chance despite retained social approach and social-odor preference and showed increased open-field center avoidance. Multiplexed chemogenetic inhibition of vCA1/Sub principal cells improved social discrimination and center exploration, whereas stronger inhibition did not reproduce the social-memory benefit. Within individual mice, discrimination was higher in predicted retuned than off-optimal states, and performance across inferred excitability states followed an inverted-U-like relationship. Anxiety-related and social improvements covaried across epileptic mice, consistent with shared dependence on vCA1/Sub circuit state. In a separate EEG cohort, moderate inhibition suppressed local hippocampal activity during the behavior-aligned interval without detectably reducing spontaneous seizures or early interictal spikes. These findings identify excessive vCA1/Sub output as a modifiable contributor to socio-affective dysfunction in TLE and show that retuning a disease-altered circuit can improve interictal behavior without acute seizure suppression.

**Highlights:**

Chronic epilepsy shifts vCA1/Sub toward a depolarized, disinhibited output state
Moderate vCA1/Sub inhibition improves social discrimination in epileptic mice
Stronger inhibition does not reproduce the behavioral benefit
vCA1/Sub inhibition improves behavior without acute seizure suppression

## Introduction

Temporal lobe epilepsy (TLE), the most common adult epilepsy, disrupts cognitive, emotional, and social function, often beyond the burden of seizures themselves ^1–3^. Current treatments are designed primarily to suppress seizures but only partially address, and may worsen, interictal cognitive and affective comorbidities, underscoring the need to identify circuit mechanisms that maintain dysfunction between seizures ^3^.

Growing evidence points to persistent hippocampal hyperexcitability, rather than seizure frequency alone, as a driver of behavioral impairment in chronic epilepsy ^4,5^. The ventral CA1-subiculum transition (vCA1/Sub) is a major hippocampal output region, where pyramidal projections to limbic and cortical targets originate, and supports memory encoding and recall, emotional regulation, and social cognition ^6–8^, and show prominent structural and functional abnormalities in TLE ^9,10^. Whether selectively normalizing hyperexcitability in this disease-altered output region can improve persistent social and emotional deficits remains unclear. Because either excessive or insufficient recruitment of hippocampal output circuits can impair memory ^4^, successful social cognition may depend on maintaining vCA1/Sub activity within a permissive functional range rather than simply minimizing excitability.

Social dysfunction is a prevalent but understudied comorbidity of epilepsy that affects quality of life ^2,11^. People with TLE frequently struggle to interpret social cues, impairing relationships and increasing isolation ^12,13^. Effective social cognition requires hippocampal processing of social memories, which can be disrupted by aberrant hippocampal excitability ^7,8^. Rodent studies likewise demonstrate that ventral hippocampal projections to limbic and cortical targets are essential for normal social behavior, and that perturbing these pathways disrupts social interaction and social memory ^14,15^. Prior work in limbic and pilocarpine epilepsy models further shows persistent social-behavior deficits associated with abnormal network activity and interictal epileptiform features, supporting these models as tools for dissecting disease mechanisms underlying social dysfunction in TLE ^4,8^.

At the cellular level, chronic TLE is characterized by loss and dysfunction of parvalbumin (PV) and somatostatin (SST) interneurons, which regulate hippocampal excitability and synchrony ^16–18^. Loss of perisomatic PV inhibition and dendrite-targeting SST/O-LM control is expected to reduce local circuit inhibition, promote pathological hyperexcitability, and degrade hippocampal-dependent behaviors, including social memory ^19^ ^6,8^. However, it remains unclear whether restoring the functional output of surviving principal neurons, despite persistent interneuron pathology, is sufficient to improve interictal behavioral deficits.

Here, we tested the hypothesis that hyperexcitable vCA1/Sub principal-cell output contributes to social memory and anxiety-related deficits in chronic epilepsy. Using a pilocarpine model of TLE, we applied bidirectional, within-animal chemogenetic modulation of vCA1/Sub principal-cell excitability, supported by interneuron quantification, ex vivo and in vivo electrophysiology, and behavioral testing. We asked whether shifting principal-cell excitability back toward a physiological range could improve social and emotional behavior in a circuit already altered by epileptic injury, and whether any such improvement was separable from acute changes in seizures. Finally, we integrated behavioral outcomes across manipulations to test whether successful social discrimination corresponded to an intermediate, permissive range of inferred vCA1/Sub excitability.

## Results

### Male Pilo mice retain social approach but show reduced investigation and impaired social recognition

If vCA1/Sub dysfunction contributes to social deficits in chronic epilepsy, Pilo mice should show impaired discrimination between familiar and novel conspecifics despite retaining basic social approach and the ability to detect socially relevant cues. We tested male control and Pilo mice using a validated social-memory task ^4,7^ ≥7 weeks after pilocarpine-induced status epilepticus, focusing on persistent behavioral dysfunction during the chronic phase rather than acute post-SE effects (**Fig. 1A**) ^20^. Mice first underwent a social-approach phase in which they explored a stimulus mouse and a novel object, followed by familiarization with the stimulus mouse and a social-discrimination test containing the familiar and a novel conspecific ^21^.

**Figure 1.**
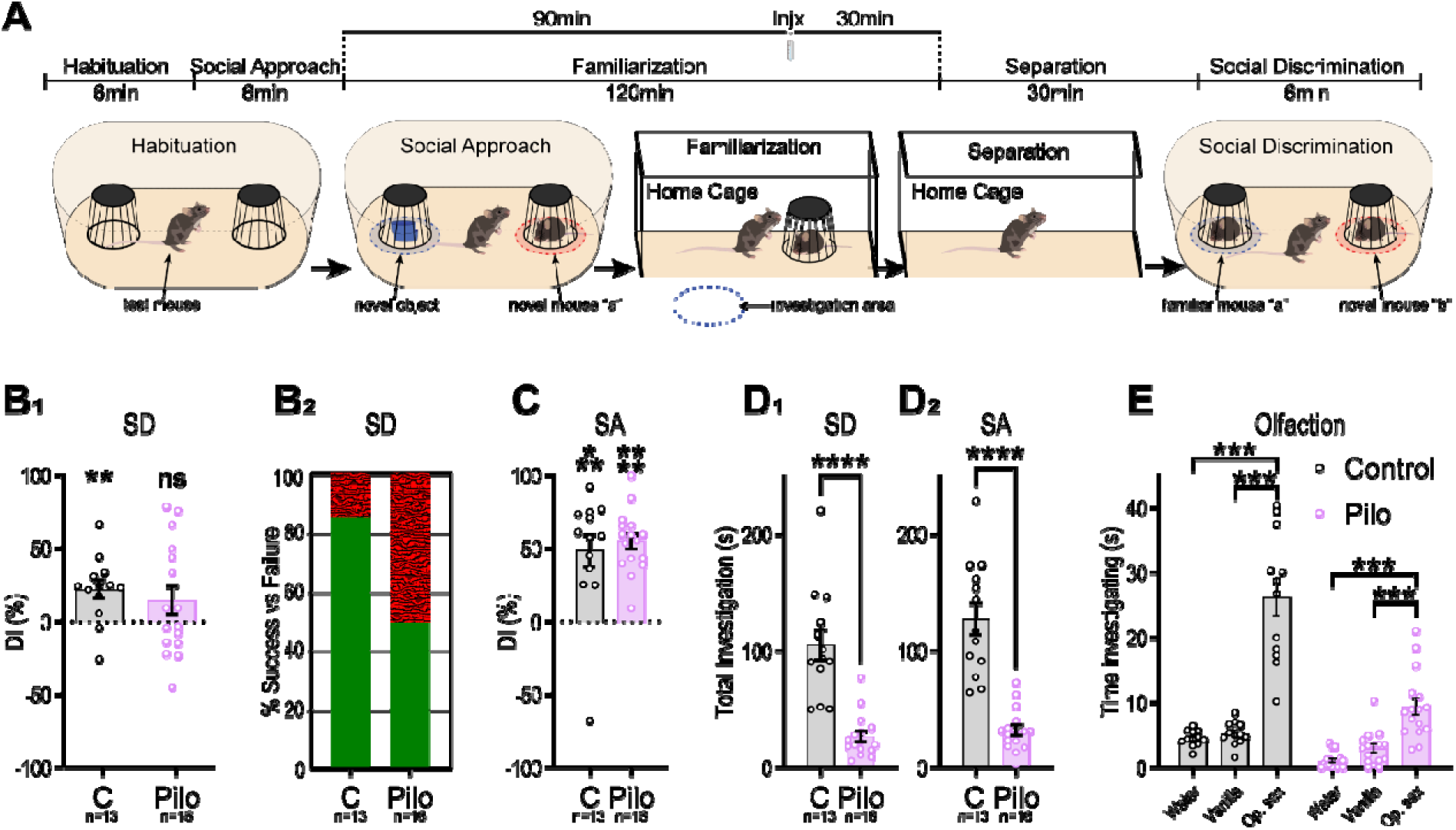
Male Pilo mice retain social approach and social-odor preference but show reduced investigation and impaired social recognition. **A**. Social-memory paradigm. Following habituation to the arena and empty stimulus cups, mice underwent a social-approach trial (SA) containing a novel object and a novel stimulus mouse. The test and stimulus mice were then familiarized for 2 h, followed by a 30-min separation. During social discrimination (SD), the test mouse explored the familiar stimulus mouse and a novel conspecific. Treatments used in later experiments were administered 1 h before social discrimination; the baseline comparison shown here reflects the corresponding control condition. **B1**. Social-discrimination index (DI), calculated as 100×(novel investigation−familiar investigation)/(novel investigation+familiar investigation). A value of 0 represents chance-level investigation. Control mice showed novel-directed discrimination, whereas Pilo mice did not differ from chance. **B2**. Descriptive proportion of mice showing novel-directed discrimination, defined as DI>0. This binary representation summarizes individual task competence and complements the continuous **DI**. **C**. Social-approach preference for the stimulus mouse relative to the novel object. Both groups preferentially investigated the social stimulus. **D1-2.** Absolute investigation was reduced in Pilo mice during the social-approach and social-discrimination phases. E. Investigation of water, vanilla, and an opposite-sex social odor. Although overall investigation was lower in Pilo mice, both groups preferentially investigated the social odor. *Points represent individual animals; bars show mean±SEM. One-sample t tests versus 0 were used for discrimination and social-approach indices. Between-group investigation times were compared using Welch’s t tests. Odor investigation was analyzed using mixed-effects models with corrected post hoc comparisons. p<0.05 *, p<0.01 **, p<0.001 ***, p<0.0001 \*\*\*\**.

During social discrimination, control mice had DI values significantly above DI=0%, indicating greater than 50% allocation of conspecific investigation to the novel mouse, demonstrating novel-directed discrimination above chance (single-sample *t* test versus DI=0%: *t_(12)_*=3.45, *p*<0.01; **Fig. 1B1**). In contrast, Pilo mice did not demonstrate novel-directed discrimination above chance at the group level (*t*_(15)_=1.43, *p*=0.17). Consistent with these continuous outcomes, a numerically larger proportion of control than Pilo mice crossed the chance boundary by directing more investigation toward the novel conspecific (**Fig. 1B2**). Thus, control mice demonstrated group-level social discrimination, whereas chronic Pilo mice did not.

Despite this difference in task performance, Pilo mice retained relative social preference. During social approach, both control and Pilo mice devoted significantly more investigation to the stimulus mouse than to the novel object (control: *t*_(12)_=4.296, *p*=0.001; Pilo: *t*_(15)_=10.23, *p*<0.0001; **Fig. 1C**). However, Pilo mice showed substantially lower absolute investigatory engagement.

They spent less time investigating the stimulus mouse than controls (control: 97±41 s; Pilo: 30±23 s; *t*_(26.51)_=6.36, *p*<0.0001) and less time interacting with the available stimulus cups overall (*t*_(14.62)_=6.58, *p*<0.0001; **Fig. 1D1–2**). Chronically epileptic mice therefore remained socially oriented but engaged less with the stimuli and failed to demonstrate social recognition at the group level.

Because social recognition in mice depends heavily on olfactory information ^22^, and pilocarpine-induced SE can damage olfactory structures ^23^, we next asked whether Pilo mice could detect and preferentially investigate social odors. Both control and Pilo mice investigated an opposite-sex social odor significantly longer than water or vanilla (all relevant pairwise comparisons, *p*<0.001; **Fig. 1E**; full statistics in **Table S2**). Although Pilo mice again showed lower overall investigation, they retained a clear preference for socially relevant olfactory cues. Their inconsistent social discrimination is therefore not readily explained by a gross inability to detect or preferentially investigate social odors.

### vCA1/Sub principal neurons are depolarized and show subtype-specific excitability changes in Pilo mice

Persistent hippocampal hyperexcitability is a defining feature of chronic epilepsy ^24–26^ and has been linked to impaired hippocampal-dependent recall, such as contextual fear conditioning (CFC) ^5^, spatial memory, and even social memory ^4^. We therefore asked whether principal neurons within the vCA1/Sub output region exhibit altered intrinsic excitability in Pilo mice and whether their activity can be shifted bidirectionally using chemogenetic tools. Inhibitory hM4D(Gi) receptors were expressed in vCA1/Sub principal neurons of Pilo mice, whereas excitatory hM3D(Gq) receptors were expressed in controls (**Fig. S2**). Whole-cell recordings were obtained from mCherry-positive principal neurons within the distal vCA1/proximal subiculum transition zone in acute horizontal slices (**Fig. 2A**).

**Figure 2.**
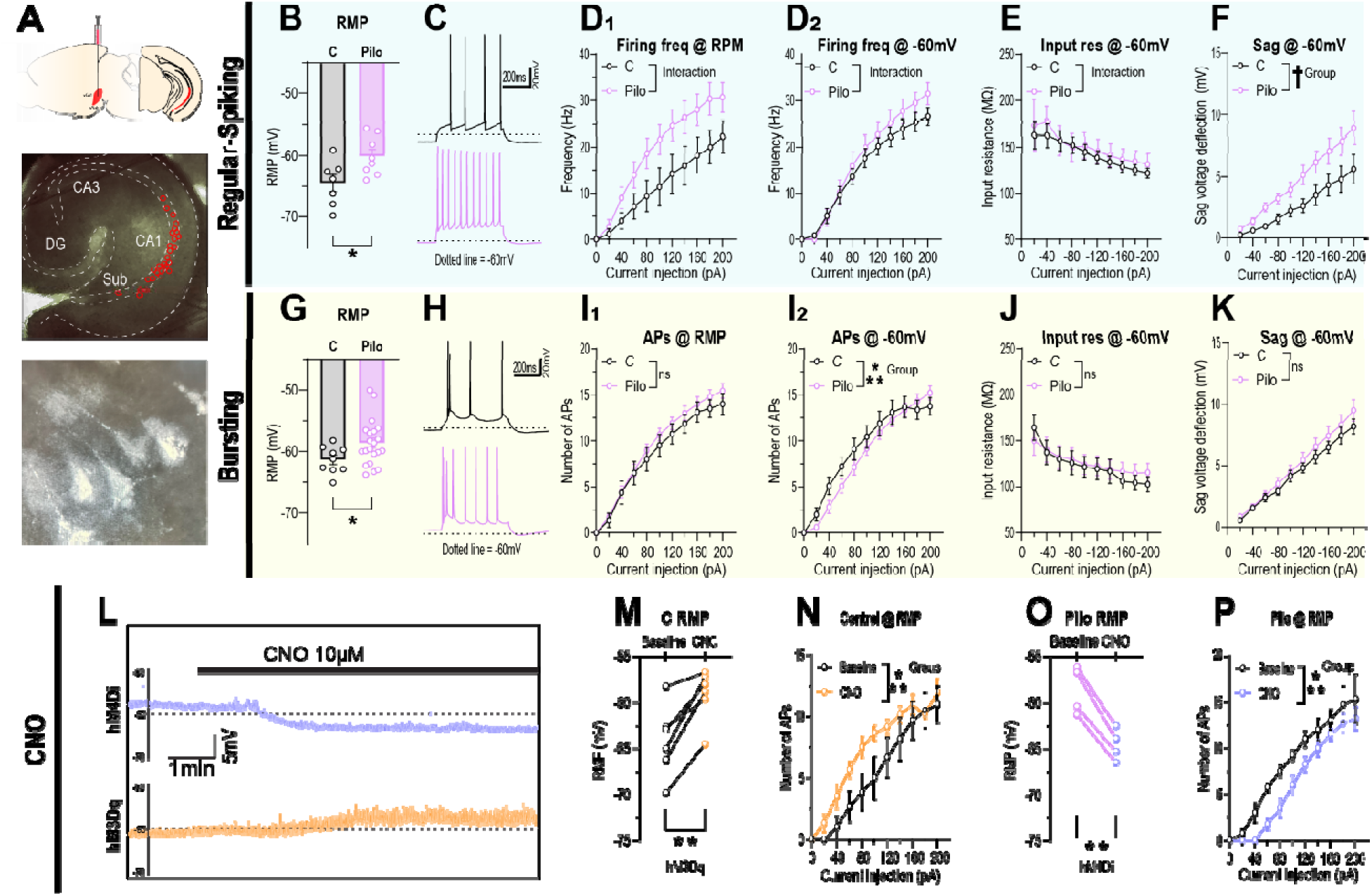
vCA1/Sub principal neurons are depolarized in Pilo mice and can be bidirectionally retuned chemogenetically. **A.** Top, schematic of the viral injection and recording region encompassing distal ventral CA1 and adjacent proximal ventral subiculum. Middle, representative horizontal hippocampal section showing the distribution of recorded mCherry-positive cells within the distal vCA1/proximal vSub transition zone. Bottom, representative mCherry-positive principal neuron selected for whole-cell recording. **Regular-spiking neurons: B.** Resting membrane potential was more depolarized in Pilo than control neurons. **C.** Representative responses from control and Pilo neurons during depolarizing current injection. **D1–2.** Firing-frequency input–output relationships measured from each cell’s native resting membrane potential (**D1**) and after holding cells at −60 mV (**D2**). Pilo neurons showed greater current-evoked firing under both conditions, reflected by significant group × current interactions. **E.** Input resistance measured during hyperpolarizing current steps from −60 mV. **F.** Voltage-sag deflection during hyperpolarizing current steps. **Bursting neurons: G.** Resting membrane potential was more depolarized in Pilo than control neurons. **H.** Representative current-evoked bursting responses. **I1–2.** Number of action potentials generated during depolarizing current steps from the native resting membrane potential (**I1**) and after holding cells at −60 mV (**I2**). Spike output did not differ at the native resting potential, whereas Pilo neurons generated fewer spikes at lower current injections when held at −60 mV, producing a significant group × current interaction. **J.** Input resistance. **K.** Voltage-sag deflection **Chemogenetic validation: L.** Representative continuous recordings during bath application of CNO (10 µM) from a Pilo neuron expressing hM4D(Gi) and a control neuron expressing hM3D(Gq). **M.** Paired resting membrane potentials before and after CNO in hM3D(Gq)-expressing control neurons. **N.** Current-evoked firing before and after CNO in hM3D(Gq)-expressing control neurons. **O.** Paired resting membrane potentials before and after CNO in hM4D(Gi)-expressing Pilo neurons. **P.** Current-evoked firing before and after CNO in hM4D(Gi)-expressing Pilo neurons. Points represent individual recorded cells; paired points or lines represent repeated measurements from the same cell. Input–output relationships were analyzed using mixed-effects models including current, group or treatment, and their interaction. Resting membrane-potential changes following CNO were analyzed using paired *t* tests. Error bars show mean±SEM. Exact model estimates, sample sizes, and complete statistical results are provided in Table S3. *p*<0.05 *, *p*<0.01 **, *p*<0.001 ***. **Sample sizes:** For regular-spiking neurons, control and Pilo samples were, respectively: RMP, 7 cells from 4 mice and 8 cells from 3 mice (B); firing at native RMP, 7/4 and 6/3 (D1); and firing, input resistance, and sag at −60 mV, 7/4 and 7/3 (D2, E, F). For bursting neurons, samples were: RMP, 10 cells from 5 mice and 22 cells from 7 mice (G); firing at native RMP, 8/3 and 22/7 (I1); and firing, input resistance, and sag at −60 mV, 9/5 and 20/7 (I2, J, K). For chemogenetic validation, the control hM3Dq cohort included 15 baseline cells and 6 CNO-exposed cells from 5 mice (M–N; M contains 6 paired cells), whereas the Pilo hM4Di cohort included 28 baseline cells and 5 CNO-exposed cells from 7 mice (O–P; O contains 4 paired cells). Cell numbers varied slightly at isolated current steps because of missing or excluded sweeps.

Recorded neurons were classified physiologically as regular-spiking or bursting cells ^27,28^ and analyzed separately because these populations differ in their intrinsic firing properties and projection patterns ^29,30^. Regular-spiking neurons from Pilo mice had resting membrane potentials approximately 4 mV more depolarized than those from controls (Pilo: −60.25±3.10 mV; control: −64.46±3.71 mV; *p*=0.033; **Fig. 2B**). Representative traces illustrate the firing patterns evoked by depolarizing current injection in control and Pilo regular-spiking neurons (**Fig. 2C**). Across increasing current steps, Pilo regular-spiking neurons generated more action potentials than control neurons when tested from their native resting membrane potentials, producing a significant group × current interaction (*p*=0.002; **Fig. 2D1**). This difference persisted when cells were held at −60 mV before current injection (*p*=0.002; **Fig. 2D2**), indicating that the increased input–output gain was not explained solely by the depolarized resting membrane potential.

Input resistance decreased across progressively larger hyperpolarizing current steps in both groups. Although the relationship between input resistance and current differed modestly by group, there was no overall group difference in input resistance at −60 mV (**Fig. 2E**; complete model statistics in **Fig. S3** and **Table S3**). Voltage-sag deflection also increased with stronger hyperpolarizing currents in both groups, without a clear overall difference between control and Pilo regular-spiking neurons (**Fig. 2F**).

Bursting neurons showed a similar disease-associated shift in resting membrane potential. Pilo bursting neurons were approximately 3 mV more depolarized than control bursting neurons (Pilo: −58.50±3.59 mV; control: −61.72±2.38 mV; *p*=0.018; **Fig. 2G**). Representative current-evoked bursting responses are shown in **Fig. 2H**. Unlike regular-spiking neurons, bursting neurons did not differ in total spike output when current steps were delivered from their native resting membrane potentials (**Fig. 2I1**). When held at −60 mV, however, Pilo bursting neurons generated fewer action potentials at lower current injections, producing a significant group × current interaction (*p*=0.001; **Fig. 2I2**). Input resistance did not differ between control and Pilo bursting neurons (**Fig. 2J**), and voltage-sag responses were also similar between groups (**Fig. 2K**).

Additional analysis of action-potential properties showed more depolarized spike thresholds and reduced spike amplitudes in both regular-spiking and bursting Pilo neurons, whereas spike half-widths did not differ (**Fig. S3**). Together, these findings identify a shared depolarization of vCA1/Sub principal neurons in chronic Pilo mice, accompanied by subtype-specific alterations in their recruitment. Increased current-evoked firing was most evident in the regular-spiking population, whereas bursting neurons showed altered recruitment only after their baseline membrane potentials were standardized.

Because resting membrane potential was consistently shifted in both neuronal subtypes, we next tested whether chemogenetic receptor activation could move vCA1/Sub principal neurons in opposite directions. Representative continuous recordings show that bath application of CNO (10 µM) depolarized an hM3D(Gq)-expressing control neuron and hyperpolarized an hM4D(Gi)-expressing Pilo neuron (**Fig. 2L**). In hM3D(Gq)-expressing control neurons, CNO shifted resting membrane potential by approximately +5 mV relative to baseline (mean change: +4.98 mV; paired *t* test, *t*_(5)_=4.988, *p*=0.004; **Fig. 2M**). This depolarization was accompanied by increased action-potential generation across depolarizing current steps (*p*<0.001; **Fig. 2N**).

Conversely, CNO hyperpolarized hM4D(Gi)-expressing Pilo neurons by approximately 6 mV (mean change: −5.93 mV; *t*_(3)_=11.09, *p*=0.002; **Fig. 2O**) and reduced current-evoked firing, with the strongest suppression at lower current steps and convergence at stronger depolarizing inputs (treatment and treatment × current effects, *p*<0.001; **Fig. 2P**). Thus, chemogenetic activation shifted vCA1/Sub principal-cell membrane potential and firing in opposite directions. In Pilo neurons, hM4D(Gi) activation moved resting membrane potential toward the range observed in control cells, establishing the physiological basis for subsequent experiments testing whether circuit retuning improves behavior.

### PV- and SST-immunopositive cell density is reduced in stratum oriens of distal vCA1 in Pilo mice

The altered membrane properties and recruitment of vCA1/Sub principal neurons could arise within a circuit that has undergone substantial changes in local inhibitory organization. PV- and SST-expressing interneurons are particularly vulnerable in chronic epilepsy and provide complementary control over pyramidal-cell output ^16–19^. We therefore quantified PV- and SST-immunopositive somata in distal vCA1 immediately adjacent to the subiculum, restricting the analysis to the CA1-laminated region in which strata oriens, pyramidale, and radiatum could be defined consistently (**Fig. 3A**).

**Figure 3.**
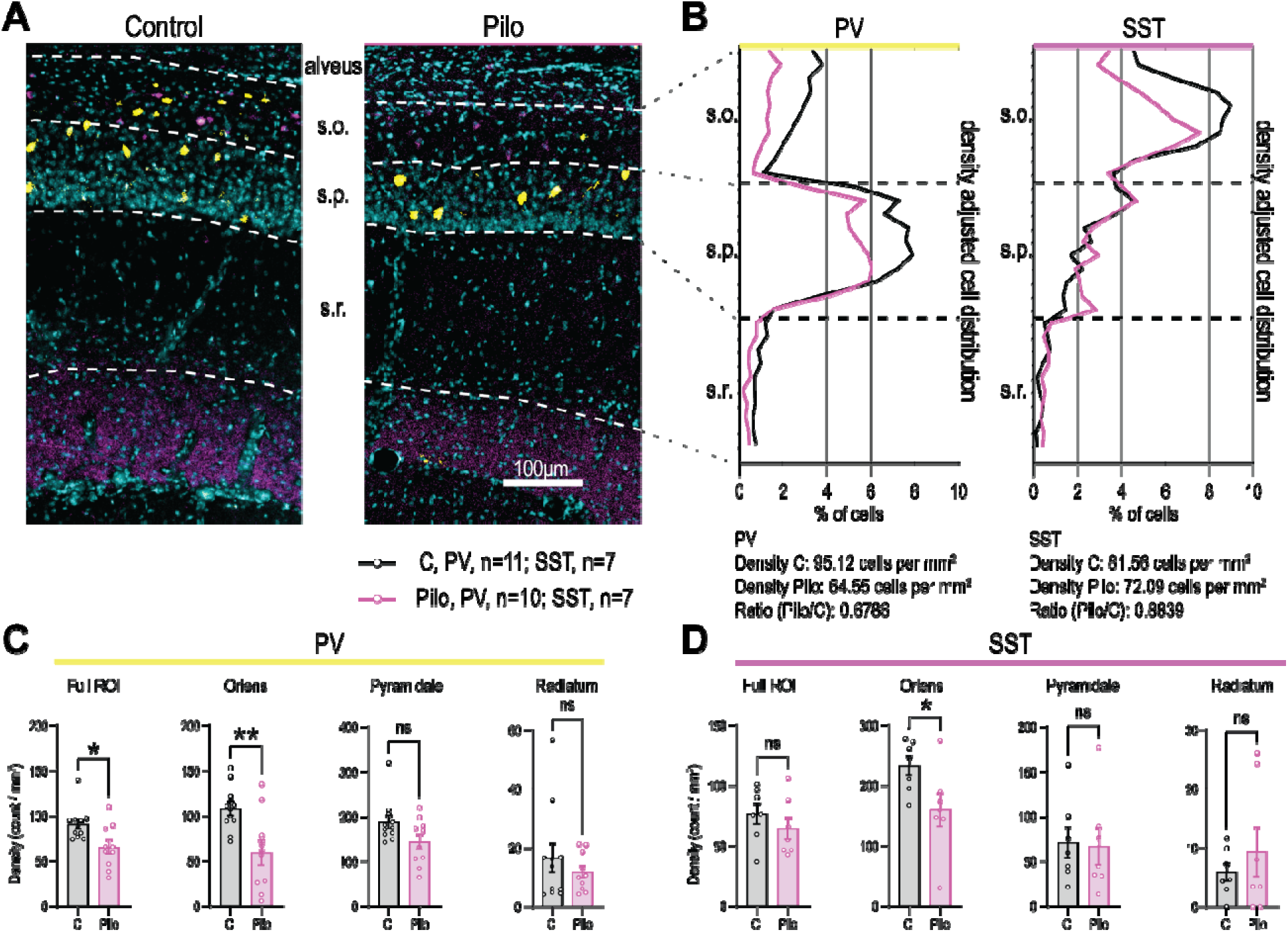
PV- and SST-immunopositive cell density is reduced in stratum oriens of distal vCA1 in Pilo mice. PV- and SST-immunopositive somata were quantified across strata oriens, pyramidale, and radiatum of distal vCA1. **A.** Representative images of distal vCA1 immediately adjacent to the subiculum in control and chronic Pilo mice. PV-immunopositive somata are shown in yellow, SST-immunopositive somata in magenta, and nuclear labeling in cyan. Dashed lines indicate the boundaries of strata oriens, pyramidale, and radiatum (s.o., s.p., and s.r., respectively). Scale bar, 100 µm. **B.** Radial distributions of PV- and SST-immunopositive somata across strata oriens, pyramidale, and radiatum. Cell-centroid locations were grouped into 10% spatial bins within each stratum. Control and Pilo distributions are shown separately for PV and SST. For visualization, Pilo distributions were scaled by the ratio of Pilo-to-control total cell density so that the profiles reflect both spatial distribution and overall group abundance. This normalization was used for visualization only; inferential comparisons were performed using mouse-level cell densities. **C.** PV-immunopositive cell density across the full distal vCA1 region and within each stratum. PV-positive density was reduced across the full region and more strongly within stratum oriens. Control, *n*=11 mice; Pilo, *n*=10 mice. **D.** SST-immunopositive cell density across the full distal vCA1 region and within each stratum. Full-region density did not differ between groups, but SST-positive density was reduced in stratum oriens. Control, *n*=7 mice; Pilo, *n*=7 mice. Each point represents the mean density for one mouse averaged across at least four tissue sections, with five sections analyzed when available. Bars show mean±SEM. Comparisons were performed using independent-samples *t* tests or Welch-corrected *t* tests where indicated. Complete statistics and effect sizes are provided in Table S4. *p*<0.05 *, *p*<0.01 **.

For each mouse, cell densities were averaged across ≥4 tissue sections, and the radial distributions of PV- and SST-immunopositive somata were mapped across the three strata using 10% spatial bins (**Fig. 3B**). These distributions showed that both cell populations were concentrated nonuniformly across the distal vCA1 laminae and provided the basis for layer-specific density comparisons.

PV-immunopositive cell density was approximately 28% lower across the full distal vCA1 region in Pilo mice than in controls (*t*_(19)_=2.685, *p*=0.0146; **Fig. 3C**). This difference was driven primarily by stratum oriens, where PV-positive density was reduced by approximately 45% (*t_(19)_* = 3.170, *p* = 0.005). PV-positive density in stratum pyramidale showed a nonsignificant reduction (*t_(19)_* = 2.004, *p* = 0.059), whereas stratum radiatum did not differ between groups (*t_(13.39)_* = 0.888, *p* = 0.39).

SST-immunopositive cell density did not differ across the full distal vCA1 region (*t*_(12)_=0.972, *p*=0.350; **Fig. 3D**), but showed a layer-specific reduction in stratum oriens. SST-positive density in this layer was approximately 32% lower in Pilo mice than controls (*t_(12)_* = 2.304, *p* = 0.039), whereas no group differences were detected in stratum pyramidale (*t_(12)_*= 0.151, *p* = 0.8824) or stratum radiatum (*t_(7.572)_* = 0.775, *p* = 0.462). The complete effect sizes were substantial for the PV and SST stratum-oriens comparisons, with η*²* values of 0.346 and 0.307, respectively.

Thus, two molecularly distinct inhibitory-cell populations showed convergent reductions in immunopositive somata within stratum oriens consistent with previous work ^31–33^. These measurements cannot distinguish interneuron loss from reduced expression of PV or SST within surviving cells, but they identify a layer-specific alteration in the inhibitory architecture surrounding the disease-shifted principal-cell population. We therefore next tested whether this anatomical pattern was accompanied by reduced functional recruitment of alveus/oriens-mediated inhibition.

### Alveus/oriens-evoked hyperpolarizing synaptic responses are reduced in Pilo vCA1/Sub principal neurons

The reduced density of PV- and SST-immunopositive somata in stratum oriens suggested that local inhibitory recruitment may be altered in chronic Pilo mice ^34^. We therefore asked whether activation of the alveus/oriens circuit produces diminished inhibitory control of vCA1/Sub principal neurons. In CA1, recurrent pyramidal-cell axons course through the alveus and recruit interneurons in stratum oriens, including O-LM and bistratified cells, which in turn provide dendritic inhibition to principal neurons ^35,36^ (**Fig. 4A**). Graded stimulation near the alveus/oriens border therefore recruits a mixed local circuit containing both excitatory axons and inhibitory interneurons, allowing the balance of evoked depolarizing and hyperpolarizing synaptic responses to be assessed.

**Figure 4.**
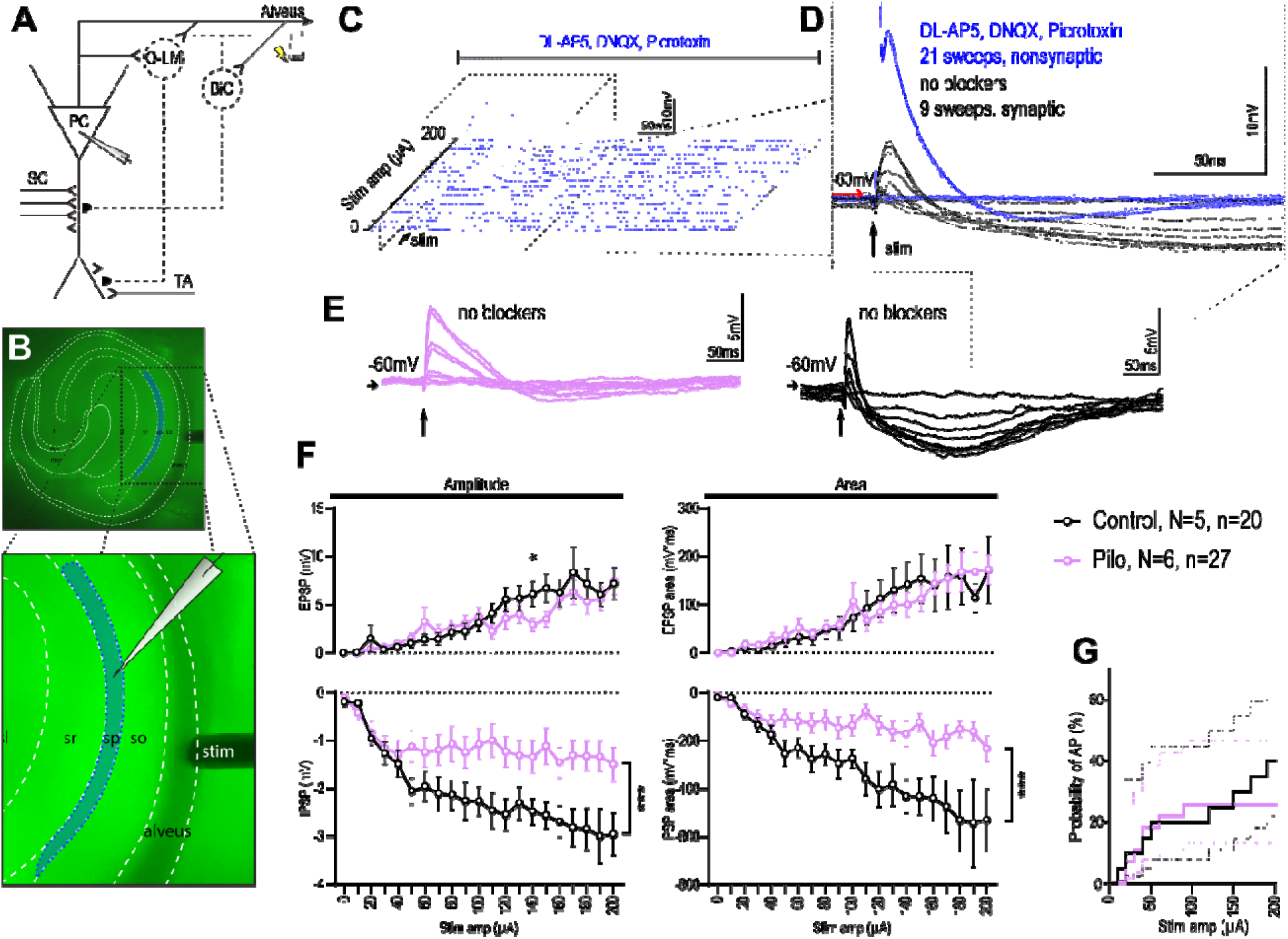
Alveus/oriens-evoked hyperpolarizing synaptic responses are reduced in Pilo vCA1/Sub principal neurons. **A**. Circuit schematic and experimental rationale. Recurrent principal-cell axons traveling through the alveus recruit interneurons in stratum oriens, including O-LM and bistratified cells, which provide inhibitory input to principal-cell dendrites. Dashed cells and connections illustrate the hypothesized reduction in inhibitory recruitment in chronic Pilo mice. The stimulation electrode was positioned near the alveus/oriens border, and principal neurons were recorded in the distal vCA1/proximal subiculum transition region. **B.** Representative horizontal hippocampal slice showing the recording region (blue) and stimulating-electrode position. Inset, principal neuron recorded in current clamp and concentric bipolar electrode positioned near the alveus/oriens border. Cells were biased to −60±1 mV before stimulation. **C.** Representative responses across the 0–200 µA stimulation series during blockade of fast glutamatergic and GABA_A_ synaptic transmission with DNQX, D-AP5, and picrotoxin. The protocol contained 21 stimulus intensities in 10-µA increments, each repeated twice. Graded subthreshold responses were absent during synaptic blockade, whereas an abrupt action potential and associated biphasic waveform appeared at higher stimulation intensities. **D**. Overlay comparing responses recorded during synaptic blockade (blue, all 21 sweeps) with graded responses obtained when synaptic transmission was intact (black, 9 sweeps). The persistent AP-associated waveform under receptor blockade was considered nonsynaptic. All sweeps containing an action potential were therefore excluded from analyses of synaptic depolarization and hyperpolarization. **E.** Representative non-AP-containing responses from Pilo (magenta) and control (black) neurons across selected nonzero stimulus intensities. Control neurons showed progressively larger hyperpolarizing responses following circuit recruitment, whereas the hyperpolarizing component was attenuated in the representative Pilo neuron. **F.** Amplitude and voltage–time integral of the depolarizing and hyperpolarizing components across stimulus intensity. Depolarization amplitude showed a modest group × stimulus interaction, whereas the depolarization integral increased with stimulus intensity without a significant group interaction. Both hyperpolarization amplitude and integral showed significant group × stimulus interactions, reflecting reduced hyperpolarizing responses in Pilo neurons, particularly at higher stimulation intensities. **G.** Kaplan–Meier representation of the stimulus intensity required to generate an antidromic action potential. AP-generation probability did not differ between control and Pilo neurons. Points and lines in F represent model-estimated or observed responses across stimulus intensity; error bars and dotted lines show mean±SEM. Repeated sweeps were averaged within cell and analyzed using mixed-effects models accounting for cells nested within animals. Complete model specifications, transformations, sample sizes, and effect sizes are provided in the Supplementary Methods and Statistics. *p*<0.05 *, *p*<0.01 **, *p*<0.001 ***.

Whole-cell current-clamp recordings were obtained from principal neurons in the distal vCA1/proximal subiculum transition region while a concentric bipolar electrode stimulated the alveus/oriens border (**Fig. 4B**). Cells were biased to −60±1 mV before stimulation to minimize the influence of the group difference in resting membrane potential identified in Figure 2. Stimulation intensity increased from 0 to 200 µA in 10-µA increments, with each intensity repeated twice. Evoked responses were variable across sweeps and could contain a depolarizing component, a hyperpolarizing component, or a depolarization followed by hyperpolarization, consistent with recruitment of recurrent excitatory and inhibitory circuitry ^37^.

At higher stimulation intensities, alveus stimulation occasionally generated an abrupt antidromic action potential in the recorded neuron. This action potential was followed by smaller biphasic voltage deflections that could otherwise be mistaken for postsynaptic responses. To determine whether these AP-associated deflections were synaptic, the stimulation series was repeated during combined blockade of AMPA/kainate, NMDA, and GABA_A_ receptors with DNQX, D-AP5, and picrotoxin (**Fig. 4C**). Under synaptic blockade, graded subthreshold responses were eliminated, whereas the abrupt action potential and its associated biphasic waveform persisted (**Fig. 4C–D**). We therefore excluded all AP-containing sweeps from subsequent synaptic analyses. Depolarizing and hyperpolarizing components measured in the remaining sweeps represented graded, blocker-sensitive synaptic responses rather than the nonsynaptic AP-associated waveform (**Fig. 4E**).

Because direct antidromic activation could differ between groups and alter the number of sweeps available for synaptic analysis, we first compared the stimulation intensity at which an action potential was generated. AP probability across the stimulation series did not differ between control and Pilo neurons (marginal Cox model: hazard ratio=0.642, 95% CI 0.224– 1.837, *p*=0.353; **Fig. 4G**). Thus, the groups did not differ detectably in their susceptibility to overt antidromic spike generation under this protocol.

We next quantified the amplitude and voltage–time integral of the hyperpolarizing synaptic component (**Fig. 4F**). Hyperpolarization amplitude increased more strongly with stimulation intensity in control than Pilo neurons, producing a robust group × stimulus interaction (*p*<0.001; _pseudo_-*R*²=0.726, *f*²=2.65). The difference became progressively larger as stronger stimulation recruited more of the local circuit. The hyperpolarization integral, which incorporates both response magnitude and duration, showed a concordant group × stimulus interaction (*p*=0.003; _pseudo_-*R*²=0.228, *f*²=0.296). Amplitude and integral were moderately correlated (*r*=0.500), indicating that they captured related but nonidentical features of the hyperpolarizing response.

Depolarizing synaptic responses were comparatively preserved. Depolarization amplitude showed a smaller group × stimulus interaction (*p*=0.004; _pseudo_-*R*²=0.230, *f*²=0.299), consistent with modestly smaller responses in Pilo neurons at portions of the stimulation range. In contrast, the depolarization integral increased strongly with stimulus intensity (*p*<0.001) without evidence of a group-dependent difference in its stimulus-response relationship. Depolarization amplitude and integral were moderately correlated (*r*=0.612).

Together, these results show a disproportionate attenuation of the hyperpolarizing component recruited by alveus/oriens stimulation in Pilo neurons. Because graded depolarizing responses remained relatively preserved, AP-generation probability was similar between groups, and the AP-associated nonsynaptic waveform was excluded pharmacologically, the findings are consistent with reduced recruitment of local inhibitory control over vCA1/Sub principal neurons in chronic epilepsy.

Reducing seizure burden can improve behavioral performance in chronic epilepsy models, and suppressing interictal epileptiform activity or its network consequences may preserve behavior even when overt seizures persist. Therefore, the behavioral effects of inhibiting vCA1/Sub principal neurons could result either from retuning circuit processes that directly support behavior or indirectly from acute suppression of epileptiform activity. This possibility is especially relevant because the ventral hippocampal formation is frequently recruited near spontaneous seizure onset in pilocarpine models and contributes to bilateral propagation and generalization.

To distinguish these possibilities, a separate cohort of five chronic Pilo mice expressing hM4D(Gi) bilaterally in vCA1/Sub underwent continuous video-EEG monitoring for approximately 2 months. A hippocampal depth electrode was positioned immediately dorsal to and adjacent to the mCherry-positive expression field, allowing electrophysiological changes near the manipulated tissue to be measured directly (**Fig. 5A**). Histological verification of electrode position and viral expression was required for inclusion.

**Figure 5.**
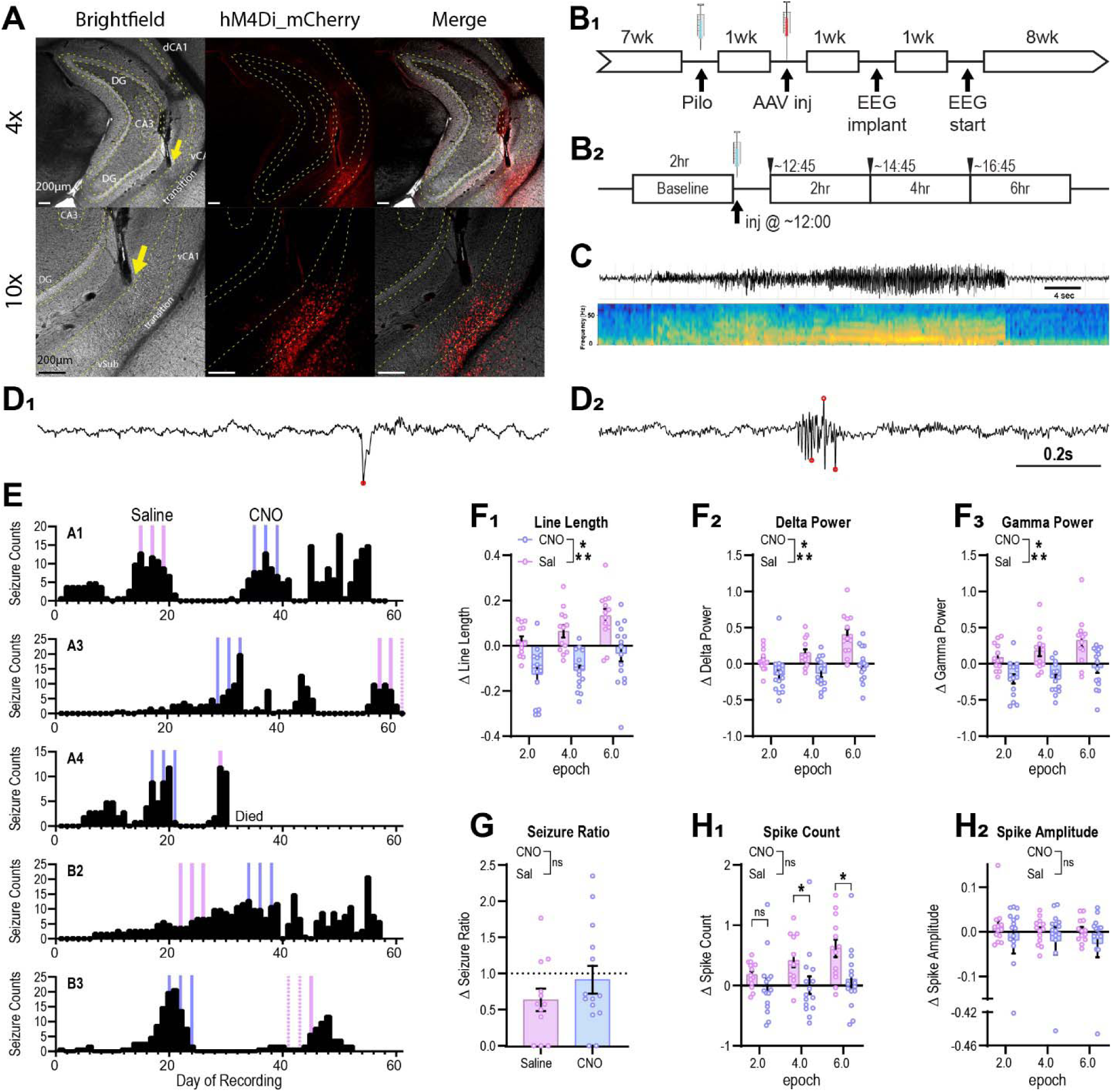
Chemogenetic inhibition of vCA1/Sub principal cells suppresses local hippocampal EEG activity without detectable acute seizure reduction. Activation of inhibitory DREADDs in vCA1/Sub pyramidal cells has immediate effects on local circuit activity, but did not detectably reduce seizures or early interictal spikes in pilocarpine mice. **A.** Histological verification of viral expression and depth-electrode placement in a representative chronic Pilo mouse. Brightfield, hM4D(Gi)-mCherry fluorescence, and merged images are shown at 4× and 10× magnification. The yellow arrow indicates the depth-electrode tract immediately dorsal to the mCherry-positive vCA1/Sub expression field. Scale bars, 200 µm. **B.** Experimental timelines (**B_1_**) Pilo mice received bilateral vCA1/Sub viral injections, followed by EEG implantation and approximately 8 weeks of continuous video-EEG recording. (**B_2_**) Each injection session included a 2 h pre-injection baseline followed by three post-injection analysis epochs: 45 min–2 h 45 min, 2 h 45 min–4 h 45 min, and 4 h 45 min–6 h 45 min. The first epoch encompassed the timing used for behavioral testing in separate behavioral cohorts. **C.** Representative confirmed electrographic seizure and corresponding time–frequency spectrogram. Seizures were identified electrographically and verified using synchronized video. **D.** Representative interictal events detected outside ictal and peri-ictal intervals. (**D_1_).** Isolated interictal spike. (**D_2_).** Clustered interictal spikes. Red markers indicate peaks meeting the prespecified detection criteria. **E.** Daily confirmed seizure counts for each of five mice across approximately 60 days of recording. Four mice showed the clustered seizure pattern characteristic of the pilocarpine model. Vertical lines indicate saline treatment days (magenta) and CNO treatment days (1.5 mg/kg; blue). Dotted treatment markers indicate sessions excluded from the primary seizure-burden analysis according to the quality-control criteria described in the Supplementary Methods. Recording for mouse A4 ended at the point indicated. **F.** Fractional changes from the session-specific pre-injection baseline in hippocampal line length (**F_1_**), raw delta power, 1–4 Hz (**F_2_**), and raw gamma power, 30–70 Hz (**F_3_**). CNO reduced all three target-engagement measures relative to saline across the post-injection period and during the first, behavior-aligned epoch. **G.** Descriptive observed-to-expected seizure ratios during the 4 h interval beginning 45 min after injection. Expected counts were estimated from the corresponding circadian interval on matched non-injection cluster days. The dotted line indicates observed seizure burden equal to the untreated expectation. Inferential testing was performed on seizure counts using a Poisson model with expected counts included as an offset; no CNO-associated reduction was detected. **H.** Interictal spike measures after exclusion of ictal intervals and the 2 min immediately before and after each seizure. **H_1_.** Composite spike frequency did not differ between CNO and saline during the first, behavior-aligned epoch, but was lower after CNO during the second and third epochs in planned temporal contrasts. The overall treatment-by-epoch interaction was not significant. **H_2_.** Composite spike amplitude did not differ during the behavior-aligned epoch. Five mice contributed repeated treatment sessions. Session-level observations are shown, and mixed-effects models accounted for repeated observations across mice, seizure clusters, injection sessions, and post-injection epochs. Bars and error bars show mean±SEM. Complete session counts, model estimates, confidence intervals, multiplicity corrections, and sensitivity analyses are provided in Table S6. *p*<0.05 *, *p*<0.01 **, *p*<0.001 ***

Because spontaneous seizures occurred in clusters, saline and CNO treatment series were initiated after the onset of an active cluster and evaluated during separate clusters (**Fig. 5B1,E**). For each injection session, the 2 h immediately before treatment served as baseline. Post-injection activity was quantified in three consecutive windows beginning 45 min after injection: 45 min–2 h 45 min, 2 h 45 min–4 h 45 min, and 4 h 45 min–6 h 45 min (**Fig. 5B2**). The first window encompassed the interval in which behavioral testing was performed in the separate behavioral cohorts and is therefore referred to as the behavior-aligned epoch.

We first evaluated local electrophysiological target engagement using hippocampal line length, raw delta power, and raw gamma power. Ictal intervals and the surrounding peri-ictal periods were excluded so that these measures reflected non-ictal activity. Values from each post-injection window were normalized to the immediately preceding session-specific baseline and analyzed using mixed-effects models accounting for repeated observations across mice, seizure clusters, injection sessions, and post-injection epochs.

Across the full post-injection period, CNO reduced line length relative to saline (β=−0.159, Holm-adjusted *p*=8.10×10⁻□; **Fig. 5F1**), as well as delta power (β=−0.290, adjusted *p*=1.19×10⁻□; **Fig. 5F2**) and gamma power (β=−0.340, adjusted *p*=1.19×10⁻□; **Fig. 5F3**). These reductions were already evident during the behavior-aligned epoch: line length (β=−0.146, adjusted *p*<0.001), delta power (β=−0.195, adjusted *p*=0.019), and gamma power (β=−0.286, adjusted *p*=0.002) were each lower after CNO than saline. Thus, hM4D(Gi) activation produced measurable suppression of local hippocampal EEG activity during the interval relevant to behavioral testing.

We next asked whether this local suppression was accompanied by an acute reduction in spontaneous seizures. Because overt seizures were considerably less frequent than the continuous EEG features, seizure burden was quantified over one 4 h interval beginning 45 min after injection rather than divided among the three 2 h epochs. Observed seizure counts were modeled relative to mouse-specific expected counts from the corresponding circadian window on matched non-injection cluster days. CNO did not produce a detectable reduction in seizure burden relative to saline after accounting for expected cluster activity (rate ratio=1.36, *p*=0.316; **Fig. 5G**).

Finally, we examined interictal spike activity after excluding ictal and peri-ictal intervals. Composite spike frequency and amplitude were quantified during the same sequential post-injection epochs. The overall treatment-by-epoch interaction for spike frequency was not significant (likelihood-ratio χ²(2)=4.38, *p*=0.112). During the behavior-aligned epoch, spike frequency did not differ between CNO and saline conditions (β=−0.193, adjusted *p*=0.282; **Fig. 5H1**), nor did spike amplitude (β=−0.033, adjusted *p*=0.562; **Fig. 5H2**). Planned temporal contrasts detected lower spike frequency after CNO during the second (β=−0.411, adjusted *p*=0.043) and third epochs (β=−0.512, adjusted *p*=0.013), indicating a delayed effect on interictal spike burden rather than an immediate change coinciding with behavioral testing.

Together, these results demonstrate clear local electrophysiological engagement of hM4D(Gi)-expressing vCA1/Sub principal neurons during the behavior-aligned interval, without a detectable acute reduction in spontaneous seizures or early interictal spike activity. The behavioral effects of vCA1/Sub inhibition are therefore unlikely to be explained solely by acute seizure suppression or immediate elimination of interictal spikes, although the later reduction in spike frequency indicates that the manipulation exerts broader network effects over time. Behavioral testing in the separate cohorts occurred within this same post-injection window, an overlap we return to when interpreting the behavioral rescue below.

### vCA1/Sub inhibition reduces anxiety-related center avoidance and rapid-movement bursts in Pilo mice

Because the ventral hippocampus is a major regulator of affective state and exploratory behavior ^38^, and chronic TLE is commonly associated with increased anxiety-related behavior ^39^, we asked whether reducing vCA1/Sub principal-cell activity would alter open-field behavior in Pilo mice. KORD was coexpressed with hM3Dq in control mice and with hM4Di in Pilo mice, allowing the same inhibitory manipulation to be applied in both healthy and disease-altered circuits. All animals received the selective KORD agonist salvinorin B (Sb; 1.5 mg/kg, s.c.) 1 h before testing. KORD-negative control and Pilo mice therefore controlled for effects of Sb unrelated to chemogenetic receptor activation. Representative movement paths suggested that KORD-negative Pilo mice explored the arena using sparse trajectories punctuated by rapid traversals, whereas KORD-positive Pilo mice showed more continuous exploration of both peripheral and central regions (**Fig. 6A**).

**Figure 6.**
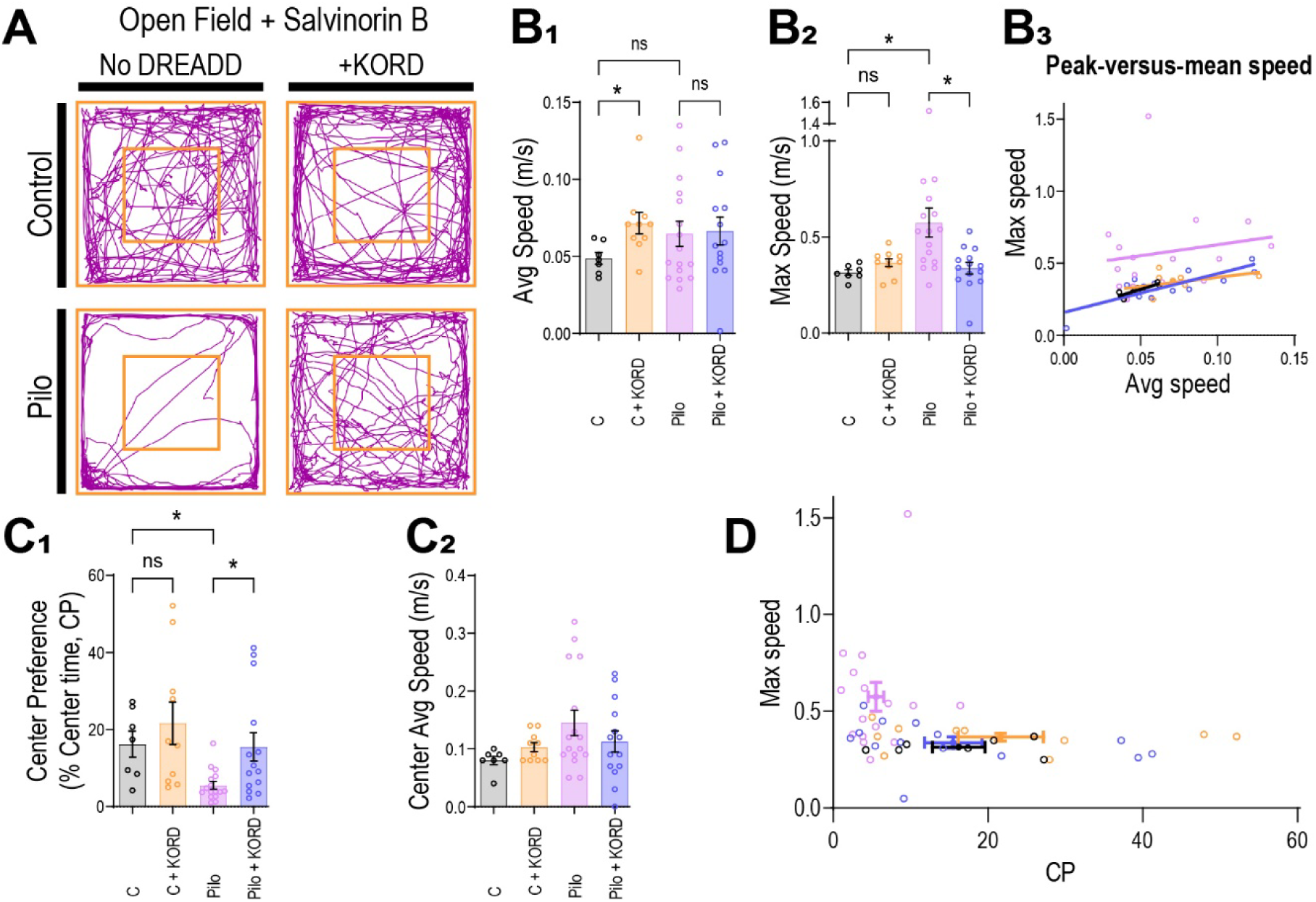
KORD-mediated inhibition of vCA1/Sub principal neurons reduces an anxiety-related open-field phenotype in Pilo mice. All mice received salvinorin B (Sb; 1.5 mg/kg, s.c.) 1 h before open-field testing. KORD-positive control mice coexpressed hM3Dq and KORD, whereas KORD-positive Pilo mice coexpressed hM4Di and KORD. Only KORD was activated during this experiment. KORD-negative control and Pilo groups controlled for ligand-related effects. **A.** Representative open-field trajectories from KORD-negative and KORD-positive control mice and KORD-negative and KORD-positive Pilo mice. The orange square indicates the center region used to calculate center preference. KORD-negative Pilo mice showed sparse exploration punctuated by rapid traversals, whereas KORD-positive Pilo mice showed more extensive exploration of the arena and center. **B_1_.** Average locomotor speed differed across groups. KORD activation increased mean speed in control mice but did not significantly alter mean speed in Pilo mice. **B_2_.** Maximum speed differed across groups. KORD-negative Pilo mice reached higher peak speeds than KORD-negative controls and KORD-positive Pilo mice. **B_3_.** Relationship between maximum and average speed. KORD-negative Pilo mice showed disproportionately high peak speeds relative to mean speed, consistent with bursts of rapid movement rather than uniformly elevated locomotion. Lines represent descriptive group-specific fits and were not used as independent inferential tests. **C_1_.** Center preference, calculated as the percentage of test time spent in the center region. KORD-negative Pilo mice showed reduced center exploration compared with controls, whereas KORD activation increased center preference in Pilo mice. KORD activation did not alter center preference in controls. **C_2_.** Average speed within the center region did not differ significantly across groups. **D.** Joint distribution of center preference and maximum speed. Open symbols represent individual mice. Crosshairs are centered on the group means, with horizontal and vertical bars indicating SEM for center preference and maximum speed, respectively. KORD-negative Pilo mice occupied a behavioral space characterized by reduced center preference and elevated maximum speed, whereas KORD-positive Pilo mice shifted toward the control distribution on both measures. This panel provides a descriptive joint representation of the two outcomes. Bars show mean±SEM, and individual points represent mice. Mean and maximum speed were analyzed using Welch’s ANOVA followed by prespecified corrected comparisons. Center preference was analyzed using a Kruskal–Wallis test followed by corrected pairwise comparisons. Complete statistics and sample sizes are provided in the supplementary tables. *p*<0.05 *; ns, not significant.

Mean locomotor speed differed across groups (Welch’s ANOVA: *F*_(3,23.16)_=3.437, *p*=0.0335; **Fig. 6B1**). KORD activation increased average speed by approximately 47% in control mice relative to KORD-negative controls (*p*=0.039). In contrast, KORD activation did not significantly alter mean speed in Pilo mice. Thus, the effect of vCA1/Sub inhibition on overall locomotor activity depended on the underlying circuit state.

Maximum speed showed a different pattern (Welch’s ANOVA: *F*_(3,23.73)_=4.498, *p=*0.012; **Fig. 6B2**). KORD-negative Pilo mice reached higher peak speeds than KORD-negative controls (*t*_(16.13)_=3.395, *p*=0.011) and KORD-positive Pilo mice (*t*_(19.59)_=2.939, *p*=0.024). KORD activation did not significantly affect maximum speed in controls. Plotting maximum against average speed showed that KORD-negative Pilo mice reached disproportionately high peak speeds relative to their mean locomotor activity, consistent with bursts of rapid movement rather than sustained hyperlocomotion (**Fig. 6B3**). The fitted lines in this panel are descriptive and were not treated as an additional inferential analysis.

Center preference, defined as the percentage of test time spent in the central region, also differed across groups (Kruskal–Wallis: *H*_(3)_=14.05, *p*=0.003; **Fig. 6C1**). KORD-negative Pilo mice spent less time in the center than KORD-negative controls, consistent with increased anxiety-related center avoidance (*p*=0.020). KORD-mediated inhibition increased center preference in Pilo mice (*p*=0.043), shifting center exploration into the range observed in controls. In contrast, KORD activation did not alter center preference in control mice (*p*>0.99).

Average speed within the center region did not differ across groups (**Fig. 6C2**). The increase in center occupancy following KORD activation in Pilo mice was therefore not accompanied by a generalized increase in movement speed within the center itself.

To visualize the combined locomotor and anxiety-related phenotype, we plotted maximum speed against center preference for each mouse and overlaid the group means and SEMs for both measures (**Fig. 6D**). KORD-negative Pilo mice occupied a distinct region characterized by low center preference and high maximum speed. KORD-positive Pilo mice shifted toward the control distribution on both dimensions, showing greater center exploration together with lower rapid-movement bursts. This panel is a descriptive joint representation of the two measured outcomes rather than a separate multivariate statistical test.

Together, these findings show that chronic Pilo mice exhibit an open-field phenotype characterized by center avoidance and bursts of rapid locomotion. KORD-mediated inhibition of vCA1/Sub principal neurons increased center exploration and reduced rapid-movement bursts in Pilo mice, shifting both measures toward the control pattern. In contrast, the same manipulation increased average locomotor speed in control mice without changing their center preference or maximum speed. The behavioral consequences of vCA1/Sub inhibition therefore depend on the preexisting circuit state.

### Moderate inhibition of vCA1/Sub principal cells improves social discrimination in Pilo mice and supports a permissive excitability range

We next tested whether directly shifting vCA1/Sub principal-cell excitability changes social-discrimination performance ^4^. Control+DREADD mice coexpressed excitatory hM3Dq and inhibitory KORD receptors, permitting movement in opposite directions from the presumed physiological state. Pilo+DREADD mice coexpressed inhibitory hM4Di and KORD receptors, allowing inhibition to be reproduced using two receptor–ligand systems. Separate control and Pilo cohorts without DREADD expression received the same ligands to assess drug-related and repeated-testing effects. Mice completed social-discrimination sessions under saline, CNO, salvinorin B (Sb), and a final saline reversal condition; Pilo+DREADD mice also received high-dose Sb (Sb3) to test stronger inhibition. Continuous discrimination index (DI) was the primary behavioral measure. We additionally summarized whether each mouse crossed the task’s chance boundary by showing the proportion with DI>0, indicating greater investigation of the novel than the familiar conspecific. This binary representation is the sign of the continuous DI rather than a separate behavioral construct. Missing DI values resulting from nonparticipation or day-of-testing seizures were addressed using multiple imputation as described in Methods; investigation-time outcomes were not imputed.

Ligand administration did not produce systematic changes in social discrimination in mice lacking DREADD expression. Continuous DI and the proportion of mice with DI>0 remained broadly stable across treatments in both control/no-DREADD and Pilo/no-DREADD cohorts (**Fig. 7A2,B2**). Total conspecific investigation likewise showed no detectable treatment-related effect in these groups. These findings argue against a substantial effect of CNO, Sb, dose, or repeated testing on social-discrimination performance in the absence of receptor expression.

**Figure 7.**
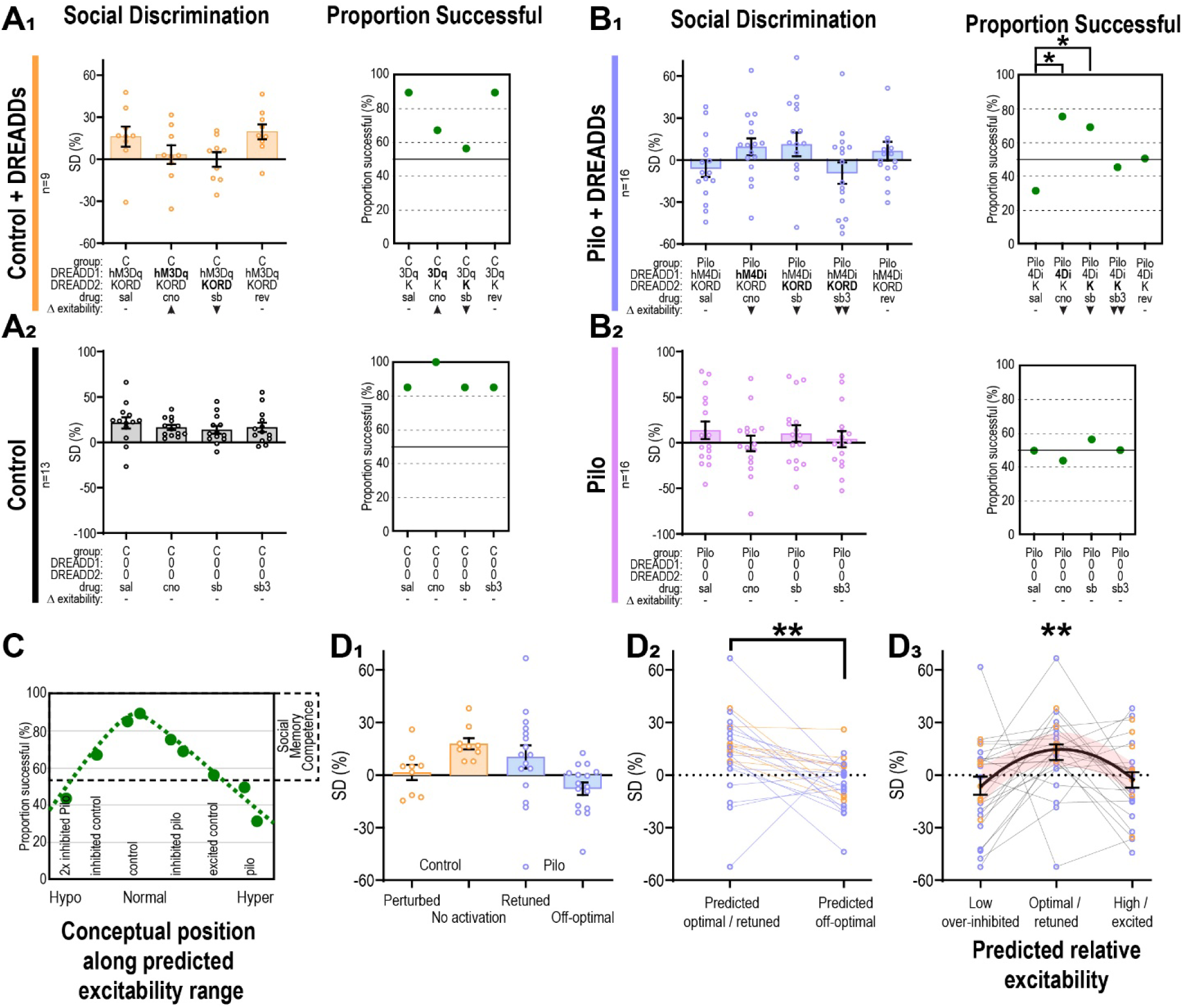
Moderate inhibition of vCA1/Sub principal cells improves social discrimination in Pilo mice and is consistent with a permissive excitability range. Social-discrimination index (DI) and the proportion of mice with DI>0 were evaluated following chemogenetic manipulations predicted to alter vCA1/Sub principal-cell excitability. DI>0 indicates that a mouse investigated the novel conspecific more than the familiar conspecific and is shown as a complementary individual-level representation of the continuous DI. **A–B. Treatment-level social-discrimination results.** Continuous DI values are shown on the left, and the proportion of mice with DI>0 is shown on the right. Columns are identified by experimental group, DREADD construct, treatment, and predicted direction of excitability change. Activated constructs are shown in bold. Symbols indicate no predicted change (–), increased excitability (▴), moderate inhibition (▾), or stronger inhibition (▾▾). Treatments were saline, CNO (1.5 mg/kg), Sb (1.5 mg/kg), and reversal saline; Pilo+DREADD mice additionally received Sb3 (3.0 mg/kg) to test stronger inhibition. **A1. Control+DREADD mice.** Non-epileptic mice coexpressed excitatory hM3Dq and inhibitory KORD receptors. CNO and Sb therefore represented perturbations in opposite directions from the presumed physiological state, whereas saline and reversal saline represented conditions without DREADD activation. Neither manipulation produced a statistically reliable treatment-level change in social-discrimination performance. **A2. Control/no-DREADD mice.** Ligand-treated non-epileptic mice without DREADD expression showed no systematic treatment-associated change in continuous DI or the proportion with DI>0. **B1. Pilo+DREADD mice.** Epileptic mice coexpressed inhibitory hM4Di and KORD receptors. Moderate inhibition with CNO or Sb shifted DI upward and increased the proportion of mice with DI>0 relative to saline. Prespecified logistic-model contrasts were significant for CNO versus saline (*p*=0.015) and Sb versus saline (*p*=0.036). Stronger inhibition with Sb3 did not reproduce this pattern. **B2. Pilo/no-DREADD mice.** Ligand-treated epileptic mice without DREADD expression showed no corresponding systematic improvement. **C. Conceptual tuning-range synthesis.** Green points show condition-level proportions of mice with DI>0. Conditions were positioned horizontally according to their predicted direction and relative magnitude of effect on vCA1/Sub excitability, based on disease state, receptor identity, ligand and dose, the electrophysiological findings of the present study, and prior work. Horizontal position does not represent a directly measured in vivo excitability value. The green inverted-U curve is conceptual and was not fitted to the condition-level proportions. The horizontal dashed line denotes 50% of the cohort with DI>0. **D1. Mouse-level predicted tuning states.** Repeated treatment sessions were collapsed into biologically defined state means. For Control+DREADD mice, the no-activation state was the mean of saline and reversal sessions, and the perturbed state was the mean of CNO and Sb sessions. For Pilo+DREADD mice, the retuned state was the mean of CNO and Sb sessions, and the off-optimal state was the mean of saline and Sb3 sessions. Points represent individual mice. **D2. Within-mouse comparison of predicted optimal and off-optimal states.** For each DREADD-expressing mouse, the predicted optimal/retuned state was compared with its predicted off-optimal state. DI was 17.43 percentage points higher in the predicted optimal/retuned state (bootstrap 95% CI, 6.80–27.55; paired *t* test, *p*=0.0038; *n*=25 mice). Lines connect state means from the same mouse. **D3. Inverted-U consistency analysis.** Mouse-level observations were assigned to predicted low/over-inhibited, optimal/retuned, or high/excited positions. For Control+DREADD mice, these positions were represented by Sb, mean(saline, reversal), and CNO, respectively. For Pilo+DREADD mice, they were represented by Sb3, mean(CNO, Sb), and saline, respectively. The black curve represents the fitted quadratic model with standard errors clustered by mouse, and shading shows the 95% confidence interval. The quadratic term was negative and significant (β₂=−17.43, 95% CI −28.26 to −6.61, *p*=0.0016), whereas the linear position term was not significant (*p*=0.679), consistent with an inverted-U-like relationship between predicted relative tuning position and social-discrimination performance. For continuous-data and state-summary panels, bars or black symbols indicate means and error bars indicate ±SEM. Proportion panels show the proportion of mice with DI>0. Orange denotes Control+DREADD mice, blue denotes Pilo+DREADD mice, black denotes Control/no-DREADD mice, and magenta denotes Pilo/no-

In Control+DREADD mice, hM3Dq/CNO and KORD/Sb shifted vCA1/Sub principal-cell activity in opposite directions (Fig. 7A1). Mean discrimination was positive under saline (16.07 ± 6.8%) and reversal (9.52 ± 6.2%) but lower under both hM3Dq/CNO (3.27 ± 6.3%) and KORD/Sb (−0.27 ± 5.8%), with similar descriptive reductions under manipulations predicted to shift activity in opposite directions from the no-activation state. Saline and reversal sessions yielded consistently high rates of novel-directed discrimination, with 8 of 9 mice showing DI>0 under both conditions, providing additional evidence against performance drift or lasting effects of repeated testing. Relative to this stable no-activation state, the proportion with DI>0 declined to 5 of 9 during KORD-mediated inhibition and 6 of 9 during hM3Dq-mediated excitation.

Although neither the continuous DI nor the treatment-level DI>0 contrasts reached statistical significance, the shared direction of these shifts was consistent with the tuning-range model: conditions predicted to move the healthy circuit away from its functional range were accompanied by lower discrimination, whereas KORD-mediated inhibition produced the opposite, beneficial effect in Pilo mice. A different pattern emerged in Pilo+DREADD mice (**Fig. 7B1**). Under saline, social-discrimination performance was variable and relatively few mice showed DI>0. Moderate inhibition with either hM4Di/CNO or KORD/Sb shifted the DI distribution upward and increased the proportion of mice showing novel-directed discrimination. Prespecified treatment contrasts indicated higher odds of DI>0 after CNO than saline (*p*=0.0146) and after Sb than saline (*p*= 0.0357). Stronger KORD activation with Sb3 did not reproduce this improvement, and performance during the final saline reversal condition returned toward the untreated pattern. Total investigation varied across sessions, but no individual treatment differed significantly from saline after correction, making a simple increase in social engagement an unlikely explanation for the discrimination effect.

The treatment-level results suggested that moderate inhibition could move the epileptic circuit into a more functional state, whereas stronger inhibition did not reproduce this benefit. However, the manipulations were not calibrated on a common in vivo excitability scale. We therefore did not use them to define numerical boundaries of circuit excitability. Instead, we integrated the conditions according to their predicted relative position based on disease state, receptor identity, ligand and dose, the ex vivo physiological effects demonstrated in Figure 2, and prior work. Condition-level proportions with DI>0 were distributed in a pattern consistent with a permissive functional range: successful discrimination was most frequent under conditions predicted to place vCA1/Sub output near an intermediate state and less frequent under conditions predicted to produce either excessive or insufficient activity (**Fig. 7C**). The horizontal positions and inverted-U curve in this panel are conceptual and do not represent directly measured in vivo excitability or a fitted dose–response function.

We next tested this model using continuous, mouse-level DI values. For Control+DREADD mice, saline and reversal sessions were averaged to represent the no-activation state, whereas CNO and Sb sessions were averaged to represent perturbation away from that state. For Pilo+DREADD mice, CNO and Sb were averaged to represent the predicted retuned state, whereas saline and Sb3 were averaged to represent the predicted off-optimal state (**Fig. 7D1**). This aggregation reduced the influence of any single spontaneous-behavior session while preserving the biologically defined comparison within each mouse.

Across the 25 DREADD-expressing mice, DI was 17.43 percentage points higher in the predicted optimal or retuned state than in the corresponding off-optimal state (bootstrap 95% CI, 6.80–27.55; paired *t* test, *p*=0.0038; **Fig. 7D2**). Thus, the same animals performed better when their treatment condition was predicted to place vCA1/Sub activity nearer the functional range.

To test whether performance followed the predicted curvature, mouse-level observations were assigned to predicted low/over-inhibited, intermediate optimal/retuned, or high/excited positions. A quadratic model with standard errors clustered by mouse showed a significant negative quadratic term (β₂=−17.43, 95% CI −28.26 to −6.61, *p*=0.0016), whereas the linear position term was not significant (*p*=0.679; **Fig. 7D3**). This pattern is consistent with an inverted-U-like relationship between predicted relative excitability and social-discrimination performance. Because excitability was not measured concurrently in each behavioral session, this analysis supports—but does not directly map—the proposed tuning range.

Because KORD-mediated inhibition also altered open-field behavior, we next examined whether affective and social responses were consistent across individual mice using complete-case observations from the overlapping cohorts. In Pilo+DREADD mice, greater center exploration during Sb treatment was associated with better social-discrimination performance and with a larger Sb-associated improvement in DI relative to the saline (unactivated) social-discrimination state. These results indicate that affective and social behavioral responses to vCA1/Sub inhibition covaried across individual Pilo mice (**Fig. S8**; complete statistics in **Table S8**).

Together, these experiments show that moderate inhibition of vCA1/Sub principal neurons improves social-discrimination performance in chronic Pilo mice, whereas stronger inhibition fails to provide the same benefit. The convergence of replicated inhibitory manipulations, ligand-treated no-DREADD controls, within-mouse state comparisons, and the quadratic state analysis supports a model in which successful social discrimination depends on maintaining vCA1/Sub output within a permissive functional range. These findings identify excessive vCA1/Sub principal-cell activity as a modifiable contributor to social-memory dysfunction in chronic epilepsy.

## Discussion

Chronic temporal lobe epilepsy (TLE) produces cognitive, emotional, and social dysfunction that often persists between seizures and remains poorly addressed by seizure-focused treatment ^1–3^. Here, we identify the ventral CA1/subiculum transition region (vCA1/Sub) as a disease-altered hippocampal output node that remains functionally modifiable in chronic epilepsy. Pilo mice showed persistent changes in vCA1/Sub principal-cell physiology together with reduced PV-and SST-immunopositive somata in stratum oriens and weaker alveus/oriens-evoked hyperpolarizing responses. Moderate inhibition of vCA1/Sub principal neurons improved social discrimination and increased open-field center exploration, whereas stronger inhibition did not reproduce the same behavioral benefit. In a separate EEG cohort, inhibition suppressed local hippocampal activity during the interval corresponding to behavioral testing without detectably reducing acute spontaneous seizures or early interictal spikes. Together, these findings support a model in which interictal behavioral dysfunction depends in part on the operating state of a chronically altered hippocampal output circuit, and can therefore improve even when the underlying pathology and epilepsy persist.

The behavioral findings provide an important starting point for this interpretation. Control mice demonstrated novel-directed social discrimination above chance, whereas chronic Pilo mice did not, despite preserved social approach and preferential investigation of social odors. Pilo mice also investigated the available stimuli substantially less overall. Thus, chronic epilepsy altered social-memory performance and behavioral engagement without eliminating social motivation or the ability to detect socially relevant olfactory cues. The direct continuous comparison of discrimination magnitude between control and Pilo groups was variable and not statistically significant, however, making the subsequent perturbation experiments particularly important. Rather than assigning mechanism from a single case–control difference, those experiments asked whether deliberately shifting the state of vCA1/Sub could change behavioral performance within a chronically epileptic circuit.

The physiological experiments indicate that this circuit alteration is not a uniform increase in excitability. Both regular-spiking and bursting vCA1/Sub principal neurons were consistently depolarized, with the clearest increase in current-evoked firing among regular-spiking neurons. These changes occurred within a region showing substantial remodeling of inhibitory circuitry. PV- and SST-immunopositive somata were both reduced in stratum oriens, and recruitment of the alveus/oriens circuit produced markedly smaller hyperpolarizing responses in Pilo principal neurons. The histological measurements cannot distinguish interneuron loss from reduced marker expression, and the stimulation experiment cannot assign the functional deficit to a single interneuron class. Nevertheless, the convergence of principal-cell depolarization, altered recruitment, layer-specific interneuron-marker reductions, and weakened recruited hyperpolarization is consistent with a persistent shift in the balance controlling vCA1/Sub output. This interpretation also accommodates prior studies showing that epilepsy can alter interneuron number, marker expression, synchronization, or functional recruitment to different degrees across models ^16–19,40–42^.

The key question was whether this altered circuit state remained behaviorally tunable. In Pilo mice, moderate inhibition through either hM4Di/CNO or KORD/Sb improved social-discrimination performance, whereas stronger KORD activation with Sb3 did not reproduce the benefit. Ligand-treated animals lacking DREADD expression showed no corresponding improvement, arguing against a simple drug effect. The replication across two independent inhibitory receptor–ligand systems is therefore consistent with an effect of reducing activity in the targeted principal-cell population. At the same time, the absence of additional benefit with stronger inhibition argues against a simple relationship in which progressively suppressing vCA1/Sub output is increasingly advantageous. In control mice, manipulations predicted to shift the circuit away from its normal state also tended to reduce discrimination, although those treatment-level contrasts were not significant. Taken together, the data are more consistent with a permissive operating range than with a model of uniformly beneficial inhibition.

This tuning-range interpretation is supported by more than the conceptual arrangement of treatment conditions in **Fig. 7C**. The manipulations were not calibrated on a common in vivo excitability scale, so their horizontal positions are necessarily inferred from disease state, receptor identity, ligand and dose, ex vivo physiology, and prior work. We therefore tested the same model using continuous within-mouse behavioral data. Across DREADD-expressing animals, social discrimination was higher when treatment conditions were predicted to place vCA1/Sub output in an optimal or retuned state than when the same mice were in predicted off-optimal states. When mouse-level observations were organized into predicted low, intermediate, and high positions, discrimination also followed significant negative quadratic curvature. These analyses do not establish numerical boundaries for an optimal excitability range, but they provide independent evidence that behavioral performance depends on relative circuit state rather than simply on the presence or magnitude of an inhibitory manipulation. This extends previous work showing that either excessive or insufficient hippocampal recruitment can disrupt behavior and that ventral hippocampal output is particularly important for social-memory processing ^4,7,8^.

The EEG experiments further distinguish circuit retuning from an acute antiseizure mechanism. In the separate chronic EEG cohort, hM4Di activation reduced hippocampal line length and delta- and gamma-band power during the first post-injection epoch, demonstrating clear electrophysiological engagement near the manipulated tissue during the interval corresponding to behavioral testing. Yet we detected no acute reduction in spontaneous seizure burden and no reduction in interictal spike frequency or amplitude during that same early period. Spike frequency decreased at later epochs in the primary analysis, but these later differences were not retained after adjustment for matched non-injection seizure-cluster activity. The most robust temporal correspondence was therefore between behavioral improvement and suppression of ongoing local hippocampal activity, not between behavioral improvement and acute elimination of seizures or interictal spikes. This distinction is important because cognitive dysfunction in epilepsy can reflect abnormal circuit recruitment, synchronization, and information processing between overt epileptic events rather than seizure burden alone ^41,43,44^.

The effects of vCA1/Sub inhibition were also evident outside the social-memory task. Chronic Pilo mice spent less time in the center of the open field and reached disproportionately high maximum speeds relative to their average locomotion, producing a pattern of center avoidance accompanied by brief rapid-movement bursts. KORD-mediated inhibition increased center preference and reduced maximum speed in Pilo mice, shifting both measures toward the control pattern. The same manipulation produced a different effect in healthy controls, increasing average speed without altering center preference or maximum speed. Thus, as in the social-memory experiment, the behavioral consequence of reducing vCA1/Sub output depended on the state of the circuit being manipulated. Among Pilo mice, greater center exploration during KORD activation also covaried with better social-discrimination performance and with greater Sb-associated improvement in discrimination. These exploratory cross-assay relationships do not establish that anxiety directly controls social memory, but they suggest that social and affective outcomes can change together when pathological ventral hippocampal output is shifted toward a more adaptive state.

This work extends our earlier circuit-based studies in complementary directions. Prior work in acquired epilepsy targeted dentate-gyrus cognitive dysfunction, whereas our previous vCA1/Sub study examined social-memory dysfunction in a 22q11.2 model ^4,43^. Here, we test the vCA1/Sub output node in acquired TLE and link behavioral retuning to convergent evidence of principal-cell depolarization, altered inhibitory architecture, reduced alveus/oriens-evoked hyperpolarizing responses, and in vivo electrophysiological target engagement. The chronic EEG cohort further addresses whether behavioral improvement coincides with local circuit modulation rather than acute seizure suppression. A central implication of these findings is that functional improvement does not require reversal of the underlying pathology. Chemogenetic inhibition did not restore PV-or SST-immunopositive populations, reconstruct local inhibitory circuitry, or eliminate chronic epilepsy. Instead, it acted downstream of those abnormalities by adjusting the output of surviving principal neurons. The therapeutic principle suggested by these experiments is therefore not chemogenetics itself, but *state-dependent compensation within a disease-altered circuit*. This distinction may be particularly important for cognitive and affective comorbidities, because global enhancement of inhibition may suppress both pathological and physiologically necessary activity. The failure of stronger KORD activation to improve social discrimination reinforces this point: the relevant objective may be restoration of useful circuit dynamics rather than maximal suppression.

This framework has broader relevance to TLE because the rodent ventral hippocampal formation corresponds approximately to the human anterior hippocampal system, which is strongly involved in memory, emotion, and psychiatric manifestations of temporal-lobe disease. Persistent interictal abnormalities in mesial temporal networks have been associated with cognitive and mood dysfunction in patients, suggesting that the behavioral burden of epilepsy may depend partly on how these networks operate between seizures ^44,45^. Existing therapeutic approaches provide proof of principle that pathological network states can be manipulated. Deep brain stimulation and responsive neurostimulation alter limbic and hippocampal activity, while interneuron transplantation, cell therapy, activity-dependent gene therapy, pharmacogenetic suppression, and closed-loop interventions offer progressively more regional, cellular, or temporal specificity ^46–55^. These strategies differ fundamentally in mechanism, but they share a potential objective with the present findings: modifying pathological network output while preserving the computations required for normal behavior.

Several limitations define the boundaries of this interpretation. Only male mice were studied because female controls did not demonstrate stable baseline discrimination in this assay, leaving sex- and hormone-dependent effects unresolved. The direct baseline comparison of continuous social-discrimination magnitude between control and Pilo groups was not statistically significant, so the strongest causal evidence comes from repeated circuit perturbation and within-mouse state analyses. The proposed tuning positions were inferred rather than measured concurrently during behavior, and the study therefore identifies a relative functional range rather than an absolute physiological set point. EEG and behavior were assessed in separate cohorts, and the EEG experiments tested hM4Di/CNO but not KORD/Sb. PV-and SST-immunopositive density cannot distinguish neuronal loss from altered marker expression, while the alveus/oriens experiment cannot identify the specific interneuron populations responsible for reduced hyperpolarization. Finally, open-field center preference is an anxiety-related measure rather than a comprehensive assay of affective state, and the cross-assay analyses were exploratory. Future studies will need to measure circuit state directly during behavior, identify the downstream projections responsible for the social and affective effects, and determine whether adaptive or closed-loop interventions can maintain vCA1/Sub output within the proposed functional range.

In conclusion, chronic epilepsy was associated with a persistently altered and disinhibited vCA1/Sub output state that remained functionally modifiable despite continued local pathology. Moderate principal-cell inhibition improved social discrimination and reduced anxiety-related center avoidance during local electrophysiological suppression, without detectable acute seizure reduction. Across manipulations, behavioral performance was more consistent with a permissive range of vCA1/Sub output than with a simple benefit of increasing inhibition. These findings support disease-state-specific circuit retuning as a strategy for addressing interictal cognitive and affective comorbidities that remain inadequately treated by seizure suppression alone.

## Materials and Methods

### Animals

Animals were single housed in a biosafety level 2 facility in ventilated cages with adequate nesting material on a 12-h light/dark cycle. Animals were provided *ad libitum* food and water and the temperature maintained at 20 ± 2 C. Additionally a white noise machine was always used in the housing and behavior rooms to reduce anxiety.

Wild-type C57BL/6(E) mice were purchased from Charles River Laboratories and used for wild-type and Pilocarpine-treatment groups as well as stimulus mice groups. Female mice have been reported to have significant shifts in their social interaction over the course of their estrus cycle ^56^ serving as a potential confound. Data collected from non-epileptic female mice presented high variability and discrimination scores which did not differ from chance (**Fig. S1**; *t_(28)_*=1.182, p=0.23) and were thus excluded from the current study. Mice were counterbalanced to experimental groups such that behavioral cohorts contained similar numbers of Pilo vs control mice with similar age. All procedures were approved by the Institutional Animal Care and Use Committee of the Children’s Hospital of Philadelphia.

### Viral Induction

Control mice were 7-9 weeks old and Pilocarpine-treated mice 7-12 days post SE induction for viral injection. Mice were anesthetized by deep isoflurane inhalation and received bilateral 200 nl viral injections (Hamilton Neuros, 7001) to ventral CA1 (vCA1). vCA1 was targeted at AP: −3.2 mm; ML: ±3.3 mm; DV: −4.5 mm to deliver viral bolus into ventral CA1 and ventral subiculum (vSub). vCA1 injections included DREADD constructs: Both AAV5-CaMKIIα-hM4D(Gi)-mCherry (Addgene viral prep # 50477-AAV5; http://n2t.net/addgene: 50477; RRID: Addgene_50477) (injected at approximately 8.80 x 10^6^ gc/injection), and AAV5-CaMKIIα-hM3D(Gq)-mCherry (Addgene viral construct # 50476-AAV5; http://n2t.net/addgene: 50476; RRID: Addgene_50476) (injected at approximately 6.20 x 10^6^ gc/injection) were gifts from Bryan Roth. The plasmid for AAV5-CaMKIIα-HA-KORD-IRES-mCitrine was purchased from Addgene (plasmid # 65418; http://n2t.net/addgene: 65418; RRID: Addgene_65418) (injected at approximately 5.20 x 10^6^ gc/injection) ^57^ and packaged into AAV5 by the CHOP Research Vector Core. AAV5-CaMKIIα-EGFP (injected at approximately 1.96 x 10^7^ gc/injection) was purchased from the Penn Vector Core.

To verify stereotaxic coordinates and viral expression, all mice were transcardially perfused with 1x Phosphate Buffered Saline (PBS) followed by 4% paraformaldehyde (PFA) fixation. Brains were removed and post-fixed in PFA overnight at 4 C. Brains were sliced and mounted for visual confirmation of viral expression. Animals with viral expression in dentate gyrus or entorhinal cortex were excluded from further analyses.

### Drugs

Clozapine *N*-oxide (CNO; Tocris) was dissolved in 4.0% dimethyl sulfoxide (DMSO)/saline to create a 10 mM stock, which was diluted in sterile saline to final doses (final DMSO < 0.5%) ^58,59^. Salvinorin B (Sb; Cayman Chemical) was dissolved in DMSO (25 mg/mL) and diluted to 5 mg/mL in sunflower oil (Sigma); this was diluted further in sunflower oil to working concentrations ^57^. CNO was administered at 5 or 10 mL/kg (0.75 or 1.5 mg/kg, respectively) i.p. and Sb was administered at 10 or 20 mL/kg (1.5 or 3.0 mg/kg, respectively) s.c.

### Status Epilepticus induction

Mice, age 7-8 weeks old underwent induction of status epilepticus (SE). Mice were pre-treated with scopolamine (1 mg/kg) to prevent peripheral damage, followed 30 minutes later by 315 mg/kg Pilocarpine (Pilo) subcutaneous injection (to avoid damage to intraperitoneal organs). Mice were monitored for a minimum of three sustained seizure events of stage 5 of the Racine scale ^60^ within an hour of Pilo treatment before administration of diazepam (5 mg/kg) to quell seizure activity. Mice that did not reach this criterion within the 1-h window were excluded because this induction profile did not reliably produce the chronic spontaneous recurrent seizures required for the epilepsy cohort. Mice were at least 7 weeks post-status epilepticus before testing began to ensure they were chronically epileptic ^20^.

### Behavior

All behaviors were semi-randomly assigned, such that cohorts of mice could be counterbalanced between tasks and treatments. All behaviors were conducted between hours (8am-1pm) during the light cycle. Social Memory (SM) and Olfaction assays were all conducted in a low-light environment following ≥ 30 (SM) or 15 (Olfaction) minute acclimation to reduced light via moving the home-cage to a curtained rack. All behavioral arenas were extensively cleaned with 70% EtOH and allowed to dry to completion prior to use.

Mice were monitored for at least one hour prior to any behavior by the experimenter to look for signs of recent (sluggishness, panting, any lack of responsiveness) or oncoming seizures (clonus, repetitive behaviors, loss of balance, spontaneous bursts of activity). Mice which underwent any behavioral seizure immediately before or during behavior were removed from the assay for that day. The purpose of this omission from behavior is to test mice which are in an interictal state and not those immediately recovering from (postictal) or actively undergoing behaviorally visible seizures. Due to the restrictions of monitoring the mice for more than 1 hour before behavior initiation we define interictal as ≥ 1 hour from behavioral seizure.

### Social Memory

Social Memory was conducted in a stadium-shaped arena (59 cm x 26 cm) with two removable chambers (inverted pencil cups of vertical bars [to allow minimal physical contact and full visual and olfactory investigation] which had plastic funnels adhered atop them to minimize climbing) into which an object or conspecific mouse may be placed. Test mice were habituated to the arena by 6-minutes of free exploration followed by return to their home cage for a 3-minute intermission. A stimulus mouse (stim) (novel 4–7-week-old same sex conspecific) was randomly assigned to one of the chambers and a novel object (bottle cap(s), Legos, or similar) was placed into the other. The test mouse was allowed to freely explore the now occupied arena for 6-minutes in a task called social approach (SA) before being returned to their home cage with the paired stim mouse. These mice freely interacted for 3-minutes (less if either attacked) after which the stimulus mouse was covered with a mesh pencil cup to allow for olfactory and visual interaction alone. This covered period lasted 2 hours to facilitate familiarization. One hour prior to the final task, test mice received an injection (Salvinorin B, CNO, or saline). Following the 2-hour familiarization, the stim mice were again allowed to freely interact with the test mice for 3 minutes before being returned to their own home cage. Thirty-minutes later the test mice were again placed into the arena with the familiarized stim mouse and a novel stim mouse each randomly assigned to a different chamber. This task, social discrimination (SD) lasted 6-minutes. Mice are social animals and will spend more time investigating a novel mouse ^21^. Social-discrimination index (DI) is calculated as 100×(novel investigation−familiar investigation)/(novel investigation+familiar investigation). Therefore, the amount of time spent investigating the novel mouse was used to create a discrimination index to determine if the test mouse could accurately recall the familiarized stim mouse, a necessary component of recognizing the other stim mouse as novel.

Social memory videos were scored via blinded video titles which underwent automated body part detection in DeepLabCut ^61^ followed by SimBA ^62^ for pose estimation and region of interest interaction analysis. Because chronic Pilo mice exhibit a readily identifiable phenotype, group identity (Control vs Pilo) could not be fully masked during live testing. However, behavioral experimenters were blinded to DREADD expression status (hM3Dq/hM4Di/KORD vs no-DREADD) and to drug condition during scoring via anonymized video IDs and a pre-specified analysis pipeline (DeepLabCut→SimBA). Specifically, each 6-minute video was given a randomly generated file number and analyzed by DeepLabCut to detect the nose, each ear, base of the skull, three spinal points, the left and right mid-points, and the tail’s base and tip.

Scoring information was fed into SimBA where the average speed and total distance traveled were measured based on the base of the skull point. The nose was used to determine when a mouse was actively interacting with a chamber (nose point within 1.5 cm of the outer edge of the chamber). This distance was chosen because the mouse had to be oriented towards the chamber for their nose to register inside of this diameter, better indicating interaction or investigation than proximity alone. Multiple samples of videos were checked from each behavioral cohort to verify correct detection and scoring.

### Olfaction

Since mice rely on olfactory cues to identify one another ^22^, we verified that experimental mice could identify social odors using an olfaction assay adapted from ^63^. This assay compares investigation (sniffing, whisking, licking, or grasping) of various odors separately. A fresh cage with clean bedding was used in which a cotton swab was secured to hang from the lid such that the cotton tip was approximately 5 cm from the cage floor. The tip was dipped into water, a 1:100 dilution of vanilla extract (McCormick), or dirtied on the soiled floor of an opposite sex conspecific (always postpubescent). Mice were placed into the olfaction chamber with a scented swab for 2-minutes before being returned to their home-cage for 3-minute intervals between trials. The duration of the mouse’s investigation of the cotton swab was annotated frame-by-frame using BORIS (v.7.10.2) ^64^ by experimenters blinded to animal and odor identification. The assay was conducted in low light during daytime hours following 15^+^ minutes of low-light adaptation.

### Open Field

To evaluate locomotor and anxiety-related behavior, mice were placed for 5 min into a square open-field arena (40 × 40 cm floor area, 39 cm wall height) under standard room illumination. For spatial analysis, the arena was divided into 16 equal sectors, and the eight middle sectors, comprising 50% of the total floor area, were designated as the center region. Center preference (CP) was calculated as the percentage of total test time spent within this region. Time spent in the center, average locomotor speed, maximum speed, center-region average speed, and total distance traveled were quantified using AnyMaze software.

### Biochemistry

#### Immunohistochemistry

The right hemisphere was sliced coronally at 50µm and used for immunohistochemistry (IHC) for the detection of Parvalbumin (PV) containing interneurons using 1:1000 (mab1572) 1 ab in mouse and 1:500 SST (PA5-82678) 1 ab in rabbit. Secondary antibodies used were goat-anti-rabbit conjugated with Alexa Fluor™ Plus 488 (1:250) and goat-anti-mouse conjugated with Alexa Fluor™ Plus 555 (1:250). Slices were rinsed with 1X PBS and incubated in sterile filtered blocking buffer (BB) of base 1X PBS containing 0.3% Triton X-100, 1% normal goat serum, and 1% bovine serum albumin for 1 hour at room temperature. Slices were transferred into tubes containing Primary antibodies diluted into BB and incubated at 4 C for 18 hours. Slices were washed in 1X PBS + 0.3% Triton X-100, 6x 10 minutes before being transferred into the secondary antibody containing solution where they incubated for 1.5 hours at room temperature. Slices were then incubated in Hoechst 1µg/mL in PBS prior to being washed (as above) before being mounted and coverslipped.

#### Imaging

PV/SST-labeled sections were imaged using a Leica DMI8-CS confocal microscope with a HC PL APO CS2 10×/0.40 objective under standardized acquisition settings. Detailed acquisition parameters are provided in the Supplementary Methods.

#### Cell counting

For PV/SST cell counting, average projections were used to draw the primary ROI which was then split into subregions using custom macros in FIJI. LabKit ^65^ was used to generate several classifiers on a subset of images with varying pixel intensities. Custom FIJI macros were used to manually compare classification outputs for each image to determine optimal object (cell) segmentation. Objects within the full ROI were counted in FIJI using 3D Objects Counter and centroid locations were used to calculate distribution within subregions using custom Python scripting. Complete scripts may be found here: https://github.com/AnthoniGM/Cell_Counting_Distribution

### Electrophysiology

#### Brain slice preparation

For ex vivo electrophysiology, a subset of mice that had completed behavioral testing was subsequently used when available. Mice were anesthetized briefly with isoflurane and transcardially perfused with ∼10 mL ice-cold, oxygenated, high sucrose cutting artificial cerebrospinal solution (aCSF) containing (in mM): NaCl 85, KCl 2.5, MgCl_2_ 4, CaCl_2_ 0.5, NaH_2_PO_4_ 1.25, NaHCO_3_ 25, D-glucose 25, sucrose 75. The mouse was rapidly decapitated, the brain removed and sliced horizontally at 300 µm using a vibrating micro slicer (Leica VT1200S, speed 1.0 mm/min, amplitude 1.2 mm) in ice cold carbogen (95% O_2_, 5% CO_2_) bubbled cutting aCSF. Slices were incubated in 50:50 cutting aCSF and regular aCSF containing (in mM): NaCl 130, KCl 3, MgCl_2_ 1, CaCl_2_ 2, NaH_2_PO_4_ 1.25, NaHCO_3_ 26, D-glucose 10, for 60 min at 35 C, then lowered to room temperature at which point they were ready for use.

#### Patch-clamp electrophysiology recordings

Glass micropipettes (Sutter Instruments O.D. 1.5 mm) were pulled using a Sutter Instruments Model P-1000 and fabricated to maintain an initial resistance of 3-5 MΩ. Whole cell, non-stimulated current-clamp recordings were conducted in regular aCSF at room temperature using an internal solution of (in mM): K-gluconate 120, EGTA 0.6, HEPES 20.0, MgCl_2_ 5.0, ATP-Na_2_ 2.0, and GTP-Tris 0.3 of pH 7.32 and 284 mOsmol. Electrophysiological recordings were made randomly from mCherry-expressing principal hippocampal neurons throughout the pyramidal and polymorphic layers between ventral CA1 and subiculum areas. mCherry-expressing CA1/Sub neurons were identified using a microscope equipped with epifluorescence and IR-DIC optics. Intrinsic excitability of these neurons was characterized by using a multi-step protocol which consisted of injecting a family of depolarizing (0-200 pA) current pulses of 500 ms duration in 20 pA increments followed by a series of hyperpolarizing currents of the same duration stepping from 0 to −200 pA in 20 pA increments. Regular-spiking and depolarization-induced burst firing properties of vCA1/Sub neurons were characterized by examining the membrane responses to depolarizing current injections at the membrane potential of −60±1 mV and at resting conditions. The principal hippocampal neurons were classified as regular-spiking and bursting cells to account for electrophysiological differences in these populations using the criteria defined in our previous study ^27^ and others ^28^. Subsequent action potential frequencies (per pulse and per burst) and input resistances were determined. The membrane potential was measured at the beginning of each recording and was not corrected for the liquid junction potential, which was around 10 mV in our experiments. The membrane input resistance was calculated by dividing the end of steady-state hyperpolarizing voltage deflection by the injected current. Neuronal membrane responses were recorded using a Multiclamp 700B amplifier (Molecular Devices, CA, USA). Voltage current commands and digitization of the resulting voltages and currents were performed with Clampex 11.2 software (Molecular Devices), and voltage and current traces were analyzed using Clampfit 11.2 (Molecular Devices).

Whole-cell, stimulated current-clamp recordings were conducted in low Mg²⁺ aCSF containing (in mM): NaCl 130, KCl 3, MgCl₂ 0.5, CaCl₂ 2, NaH₂PO₄ 1.25, NaHCO₃ 26, and D-glucose 10, at 33 °C, using an internal K-gluconate solution. A concentric bipolar stimulating electrode (FHC) was placed in the stratum oriens near the alveus border (O/A).

Electrophysiological recordings were obtained randomly from mCherry-expressing principal hippocampal neurons within the pyramidal and polymorphic layers between the ventral CA1 and subiculum. Each recording consisted of 42 sweeps, each 500 ms in duration, with a stimulation pulse (0.1 ms) delivered at 58.5 ms. Stimulation intensity increased in 10 µA steps from 0– 200 µA, with each level repeated twice. As previously reported, the response to O/A stimulation is biphasic and stochastic ^37^; therefore, experiments were often repeated within the same cell, and responses at each stimulation intensity were averaged. Back-propagating action potentials were frequently observed and were followed by a biphasic potential that was determined to be non-synaptic; therefore, only non-AP-associated changes in membrane potential were used for analysis.

Voltage sweeps were analyzed using a custom Python pipeline. Action-potential-containing sweeps were excluded, and depolarizing and hyperpolarizing response amplitudes and voltage–time integrals were extracted from the remaining sweeps. Detailed signal-processing and event-detection parameters are provided in the Supplementary Methods.

### In vivo video-EEG

#### Video-EEG cohort and electrode implantation

A separate cohort of chronic pilocarpine-treated mice expressing AAV5-CaMKIIα-hM4D(Gi)-mCherry bilaterally in vCA1/Sub was used for continuous video-EEG experiments. Following establishment of chronic spontaneous seizures and sufficient time for viral expression, mice were anesthetized with isoflurane and implanted with two cortical screw electrodes positioned at AP +0.5 mm and ML ±1.7 mm, two cerebellar screw electrodes positioned at AP −5.5 mm and ML ±2.0 mm for ground and reference, and a twisted pair of hippocampal depth-wire electrodes lowered immediately dorsal to the vCA1/Sub viral-expression field. The electrode assembly was secured using dental cement (Lang Dental). Following recording, electrode placement and viral expression were verified histologically; animals with electrode placement outside the targeted hippocampal region or inadequate or off-target viral expression were excluded. Representative electrode placement and viral expression are shown in Fig. 5A.

Beginning one week after implantation, mice were acclimated to recording cages and connected to a tethered video-EEG system controlled through Stellate Harmonie software. Four-channel EEG, consisting of hippocampal depth-wire and cortical screw recordings, was acquired continuously at 2,000 Hz with synchronized video for approximately 2 months. Unless otherwise indicated, quantitative EEG analyses used the hippocampal depth-wire recording.

#### Video-EEG treatment and analysis design

Spontaneous seizures occurred in clusters consisting of several days of elevated seizure burden separated by periods of few or no seizures. Treatment series were therefore initiated after the onset of an active spontaneous-seizure cluster. Mice received CNO (1.5 mg/kg, i.p.) or saline vehicle every other day for three injections administered at approximately the same time during the light phase. Saline and CNO series were evaluated during separate seizure clusters. For each injection session, the 2-h period immediately preceding injection served as the session-specific baseline. Post-injection activity was evaluated in three consecutive 2-h windows beginning 45 min after administration: 45 min–2 h 45 min, 2 h 45 min–4 h 45 min, and 4 h 45 min–6 h 45 min. The first post-injection window encompassed the timing used for behavioral testing in the separate behavioral cohorts.

Seizures were identified electrographically using a custom detection pipeline and confirmed by manual review of synchronized EEG and video. Verified ictal intervals and the 2 min immediately preceding and following each seizure were excluded from analyses of non-ictal EEG activity. The prespecified target-engagement measures were hippocampal line length, raw delta power (1–4 Hz), and raw gamma power (30–70 Hz). Interictal spike frequency and amplitude were quantified separately. Post-injection EEG measures were expressed as fractional change from the corresponding session-specific baseline.

Because overt seizures were substantially less frequent than continuous EEG features, seizure burden was quantified over a single 4-h interval beginning 45 min after injection. Observed seizure counts were evaluated relative to mouse-specific expected counts derived from the corresponding circadian interval on non-injection seizure-cluster days. Detailed seizure-detection criteria, signal-processing parameters, interictal-spike detection, cluster-reference sensitivity analyses, and seizure-count quality-control procedures are provided in the Supplementary Methods.

### AI assistance disclosure

We used ChatGPT (OpenAI) to provide language suggestions (grammar, clarity, concision) for draft text and to assist in reorganizing and commenting Python code used for data analysis. Prompts generally asked for rephrasing of technical text for readability or for clarification of code comments without altering functionality. Representative prompts are included in **S. Table 9**; the full chat transcripts spanning tool development were not retained, as many iterations were discarded and not incorporated into the final work. All outputs were reviewed, edited, and verified by the authors, who accept full responsibility for the content. No figures, images, or other media in this paper were generated by AI.

### Statistics

Statistical outputs for primary comparisons are reported in the Results in abbreviated form, with complete model estimates, effect sizes, confidence intervals, sample sizes, and multiplicity corrections provided in the supplementary tables organized by figure. All tests were two-sided with α=0.05 unless otherwise specified.

#### Behavior

Behavioral analyses were conducted in GraphPad Prism and Python. Direct comparisons were performed using t tests or nonparametric alternatives where appropriate, and repeated-treatment outcomes were evaluated using repeated-measures, mixed-model, or regression approaches as specified for each analysis.

Nonparticipation and day-of-testing seizures resulted in approximately 18% missing social-discrimination data. Missing discrimination-index (DI) values were addressed using multiple imputation with bootstrap resampling to represent uncertainty in the imputed observations. Sixty imputed datasets were retained and pooled for the primary treatment-level analyses. Investigation-time outcomes were not imputed. Imputation diagnostics are shown in **Fig. S5** and **Table S1**, and behavioral participation and attrition are summarized in **Fig. S9**.

Treatment-level DI>0 outcomes were analyzed using logistic regression with treatment and treatment order/time as predictors and saline as the reference condition. For mouse-level tuning-state analyses, repeated sessions were collapsed using prespecified biological state definitions. In Control+DREADD mice, the no-activation/optimal state was the mean of saline and reversal sessions and the perturbed/off-optimal state was the mean of CNO and Sb sessions. In Pilo+DREADD mice, the retuned/optimal state was the mean of CNO and Sb sessions and the off-optimal state was the mean of saline and Sb3 sessions. Optimal/retuned and off-optimal states were compared within mouse using a paired t test with a bootstrap confidence interval for the paired mean difference. To test the predicted curvature, mouse-level observations were assigned to low/over-inhibited, intermediate optimal/retuned, and high/excited positions and analyzed using an ordinary least-squares quadratic model containing linear and squared position terms, with standard errors clustered by mouse to account for repeated observations from the same animal. The conceptual curve in **Fig. 7C** was not fit to the condition-level proportions.

Cross-assay relationships between open-field center preference and social-discrimination outcomes were evaluated using complete-case observations only; imputed DI values were excluded. Detailed regression and exploratory threshold-classification procedures are provided in the Supplementary Methods and **Table S8**.

#### Electrophysiology

Electrophysiological comparisons between Control and Pilo groups were performed separately for regular-spiking and bursting neurons for the intrinsic-physiology experiments in **Fig. 2**. Linear mixed-effects models incorporated experimental group and current injection or stimulation intensity as fixed effects where appropriate, with repeated observations accounted for at the cell and animal levels. Transformations were applied when required by outcome distributions. For alveus/oriens stimulation experiments in **Fig. 4**, repeated sweeps were averaged within cell and mixed-effects models accounted for cells nested within animals. Complete model specifications, transformations, sample sizes, and effect sizes are provided in Tables S3 and S5.

#### Video-EEG statistics

Continuous EEG outcomes were analyzed using linear mixed-effects models including treatment condition, post-injection epoch, treatment × epoch, and injection day, with repeated observations accounted for across mice, seizure clusters, and injection sessions. Treatment-by-epoch interactions were evaluated by likelihood-ratio comparison of interaction and additive models. Prespecified CNO–saline contrasts were Holm-adjusted within defined outcome families.

The primary target-engagement family comprised line length, delta power, and gamma power. Composite spike frequency and amplitude during the first, behavior-aligned post-injection epoch formed a separate multiplicity-corrected family, and planned temporal contrasts for spike frequency were corrected across the three post-injection epochs.

Observed seizure counts during the 4-h target window were analyzed using a Poisson generalized linear model with treatment, injection day, and mouse as predictors and the logarithm of mouse-specific expected seizure count as an offset. Robust standard errors were used. Cluster-reference-adjusted EEG analyses and seizure-count quality-control analyses were performed as sensitivity analyses. Full model specifications, multiplicity-correction families, sensitivity analyses, and session counts are provided in the Supplementary Methods and **Table S6**.

## Supporting information

Supplemental results and methods

## Declaration of generative AI and AI-assisted technologies in the manuscript preparation process

During the preparation of this work, the author(s) used [OpenAI, ChatGPT] for [grammatical edits, structural notes, and statistical organization]. The author(s) reviewed and edited the output as needed and take full responsibility for the content of the published article.

## Acknowledgments

We would like to acknowledge and thank Alicia White for her aid in colony management and generating chronic epileptic mice, without which this project would not be possible.

## Author contributions

Conceptualization: JBK, AMG, DAC

Data Curation & Software: AMG Formal Analysis: AMG

Funding Acquisition: DAC

Investigation: AMG, ES, SJ, JBK, HT

Methodology: AMG, JBK, SJ

Resources: DAC, HT

Visualization: AMG

Supervision: DAC, SJ

Writing—original draft: AMG

Writing—review & editing: AMG, SJ, JBK

## Data and materials availability

All data, code, and materials used in the analyses are available to any researcher for the purpose of reproducing or extending the analyses. Please contact the corresponding author for access to any resources outside of those which require a materials transfer agreement (MTA) which must be made separately to the appropriate institutions.

