## Supplemental results and methods for "Retuning ventral hippocampal output improves social memory and reduces anxiety-related behavior in temporal lobe epilepsy"

Anthoni Goodman *et al.*

**This PDF file includes:**

Expanded text including sections: Methods

Figs. S1 to S9

Tables S1 to S9

Supplementary Methods

**Imaging:**

Images for PV/SST labeled tissue were acquired using **Leica DMI8-CS** confocal microscope equipped with a HC PL APO CS2 10x/0.40 DRY objective. A Diode (405nm) and an Argon laser (458 nm) were used for excitation with an output power of 5% each. Photomultiplier tube (PMT) detectors with spectral ranges configured for 380-385, 495–550nm, and 558.1–571.35nm with a pinhole size of 5.30 µm (.998 Airy unit) were used to capture images. 8bit images were created at 1024x1024 with a resolution of 0.86 pixels/µm and 2.0µm depth.

**Analysis of alveus/oriens-evoked responses:**

Voltage sweeps were analyzed using a custom Python script. Traces were smoothed with a Gaussian filter (σ = 2). Baseline voltage was defined as the mean membrane potential from 1.5 to 57 ms. Antidromic action potentials were identified between 58.3 and 60 ms. In sweeps without action potentials, depolarization was defined as a rise of ≥1 mV above baseline, measured from threshold crossing to return to baseline or the end of the sweep. Hyperpolarization was defined as a ≥0.75mV decrease below baseline, whether occurring alone or following a depolarizing component, measured from initial threshold crossing to return above baseline. For each sweep, the script extracted the timing and amplitude of depolarization and hyperpolarization events for downstream analysis.

Extended video-EEG acquisition and analysis

Seizure identification and verification

Seizure detection was performed offline using custom MATLAB graphical user-interface modules. Candidate events were identified from all four EEG channels using three independently calculated features: median signal amplitude, line length, and a local minimum–maximum feature. Each feature was calculated in 20-s bins. Bins exceeding feature-specific candidate thresholds were assigned one point for each feature and channel, yielding a maximum candidate score of 12 points per bin across the three features and four channels. Higher scores therefore indicated stronger evidence of a sustained multichannel electrographic event.

Candidate events were reviewed manually using a MATLAB reviewer module displaying synchronized EEG channels. Video corresponding to each candidate event was independently reviewed in Stellate Harmonie to confirm behavioral seizure activity and establish seizure onset and offset. Only visually adjudicated events meeting the electrographic and video criteria were included as seizures in the seizure-count analyses.

**Exclusion of ictal and peri-ictal periods:**

Verified seizure intervals were removed from analyses intended to quantify non-ictal circuit and interictal spike activity. For each seizure, the complete ictal interval together with the 2 min preceding seizure onset and 2 min following seizure termination was excluded. Feature values within these intervals were replaced with NaN values and did not contribute to calculation of baseline or post-injection block means. This exclusion prevented seizure waveforms and immediate preictal or postictal activity from inflating line length, spectral power, or interictal spike measures.

**Continuous EEG feature extraction:**

EEG analysis and visualization were performed using custom MATLAB graphical user interfaces. For line-length analysis, hippocampal EEG was band-pass filtered from 1–70 Hz and notch filtered at 60 Hz. Line length, reflecting cumulative absolute sample-to-sample voltage change, was calculated in consecutive 1-s windows and smoothed using locally weighted scatterplot smoothing (LOWESS).

Spectral power was quantified in the delta (1–4 Hz), theta (4–8 Hz), alpha (8–13 Hz), beta (13–30 Hz), and gamma (30–70 Hz) frequency bands. For each 1-s window, raw band power, total power, and relative band power, defined as band power divided by total power, were calculated and smoothed using LOWESS. The prespecified paper-facing target-engagement outcomes were line length, raw delta power, and raw gamma power. Other spectral measures were retained in the source data as supportive or exploratory outcomes but were not included in the primary inferential comparison family.

**Interictal spike detection:**

For spike detection, hippocampal EEG was band-pass filtered from 1–200 Hz and notch filtered at 60 Hz. Positive-going spikes were detected using the MATLAB findpeaks function with a minimum peak height of five times the signal standard deviation, minimum peak prominence of two times the standard deviation, and minimum peak distance of 50 ms. Negative-going spikes were identified by applying the same procedure to the inverted EEG signal. Spike time, peak amplitude, and interspike interval were extracted separately for positive- and negative-going events.

Positive- and negative-going spike counts were initially retained separately for quality control. Because polarity primarily reflects local field orientation and source–sink geometry rather than distinct classes of epileptiform events for the present question, their normalized changes were averaged to generate a composite spike-frequency measure. Normalized positive- and negative-going spike amplitudes were similarly averaged to generate a composite spike-amplitude measure.

**Baseline normalization and post-injection analysis windows:**

For each injection session, the 2-h period immediately preceding injection served as the session-specific baseline. To permit CNO to enter the brain and activate hM4D(Gi), post-injection analysis began 45 min after administration. Three consecutive 2-h windows were defined:

1. 45 min to 2 h 45 min after injection;
2. 2 h 45 min to 4 h 45 min after injection; and
3. 4 h 45 min to 6 h 45 min after injection.

For each baseline and post-injection window, mean line length, spectral-power measures, spike count, spike amplitude, and interspike interval were calculated after exclusion of ictal and peri-ictal periods. Each post-injection value was expressed as fractional change from the corresponding pre-injection baseline:

$$\Delta X =\frac{X_{post}- X_{baseline}}{X_{baseline}}$$

where *X_post_* represents the mean value within a post-injection window and *X_baseline_* represents the mean value during the preceding 2-h baseline. Thus, values below zero indicate a reduction relative to the same session’s baseline.

**Matched non-injection EEG analysis:**
Identical baseline and post-baseline windows were applied to temporally matched non-injection days within each seizure cluster. Because injections occurred every other day, mice contributed up to three CNO sessions, three saline sessions, and corresponding non-injection days. Non-injection sessions were processed using the same filtering, ictal-exclusion, feature-extraction, and normalization procedures as active-treatment sessions.

The primary EEG analyses compared CNO directly with saline, because saline controlled for handling and injection effects. In a prespecified sensitivity analysis addressing natural within-cluster progression, the mean non-injection value from the same animal, seizure cluster, and time window was subtracted from each active-treatment observation:

$$X_{cluster-adjusted}= X_{active}- {x̄}_{non-injection, animal\times cluster\times time}$$

Cluster-adjusted CNO and saline values were then compared using the same model structure as the primary analysis. No missing non-injection observations were imputed.

**Post-injection seizure burden:**
Because overt seizures were substantially less frequent than continuous EEG features or interictal spikes, seizure counts were not subdivided into the three 2-h EEG windows. Instead, seizures occurring during a single 4-h target period beginning 45 min after each injection were counted. This window captured the period of expected hM4D(Gi) activation while providing sufficient duration to quantify relatively sparse ictal events.

Observed post-injection seizure counts were compared with mouse-specific expected counts estimated from non-injection seizure-cluster days. Expected counts were calculated from the corresponding 4-h circadian window on non-injection days from the same animal. To restrict estimation to periods of active seizure burden, only non-injection days with more than three seizures during the full 24-h recording period were included. The observed-to-expected ratio was calculated for descriptive visualization:

$$\frac{Observed}{Expected}ratio =\frac{Observed target-window seizures}{Expected target-window seizures}$$

A ratio of 1 indicates seizure burden equal to the mouse-specific untreated expectation, whereas values below 1 indicate fewer seizures than expected. Inferential analyses were performed on observed seizure counts with expected counts entered as a model offset rather than by directly analyzing these ratios.

**Seizure-count quality control:**
Two injection sessions were excluded from the primary seizure-count analysis based on dataset-quality annotations. One CNO session contained 16 seizures during the 4-h target window and was identified as an extreme observation during data review. One saline session occurred after the active seizure cluster had ended and contained no seizures over the entire 24-h recording day. The primary analysis therefore included 26 injection sessions from five mice.

Two additional saline sessions from one mouse were identified as borderline because treatment was initiated before the anticipated increase in cluster activity; each corresponding 24-h day contained only one seizure. These sessions were retained in the primary analysis because the three-injection treatment series had already been initiated, but were removed in a prespecified sensitivity analysis. This sensitivity dataset included 24 injection sessions from five mice.

**Video-EEG statistical analysis**

Continuous EEG outcomes were analyzed using linear mixed-effects models. Treatment condition (CNO versus saline), post-injection epoch, treatment × epoch, and injection day were included as fixed effects. Mouse was included as a random intercept, with additional variance components for seizure cluster and injection session:

$$Y \sim Treatment\times Epoch + InjectionDay + \left( 1 | Mouse \right)+ \left( 1 | Mouse:Cluster \right)+ \left( 1 | Mouse:Session \right)$$

Models used restricted maximum likelihood for estimation of fixed effects and variance components. Treatment-by-epoch interactions were evaluated by likelihood-ratio comparison of the full interaction model with a reduced additive model fit using maximum likelihood. Planned CNO–saline contrasts were calculated from fitted fixed effects and reported as model-estimated differences (β), standard errors, 95% confidence intervals, Wald z statistics, and two-sided p values.

The primary target-engagement outcomes were line length, delta power, and gamma power. Holm correction was applied across these three outcomes separately for the model-estimated treatment contrast averaged across post-injection epochs and for the first, behavior-aligned post-injection epoch. Composite spike frequency and composite spike amplitude during the first post-injection epoch formed a separate Holm-corrected family. Planned time-specific contrasts for composite spike frequency across the three post-injection epochs were corrected together using the Holm procedure. Cluster-reference-adjusted analyses were treated as sensitivity analyses and corrected within parallel prespecified outcome families.

Observed seizure counts during the 4-h target window were analyzed using a Poisson generalized linear model. Treatment, injection day, and mouse were included as predictors, and the logarithm of the mouse-specific expected seizure count was included as an offset:

$$\log\left( E\left[ Yi \right] \right)=\beta0+\beta1Treatment i+\beta2InjectionDay i+\beta3Mouse i+\log\left( ExpectedSeizures i \right)$$

$Yi$ is the observed seizure count for injection session $i$.

Heteroskedasticity-consistent robust standard errors were used. The exponentiated CNO coefficient is reported as the CNO-versus-saline seizure rate ratio, with values below 1 indicating reduced seizure burden after CNO after accounting for expected seizure activity. Because this was a single prespecified seizure-burden comparison, no multiplicity correction was applied. The model was repeated after removal of the two borderline cluster-timing saline sessions as a sensitivity analysis. Complete statistical output is provided in Table S6.

Cross-assay behavioral associations and exploratory classification

Relationships between open-field center preference (CP; percentage of test time spent in the center) and social-discrimination index (DI) were evaluated using observed complete-case mouse pairs only; imputed DI values were excluded. Separate simple linear regressions were performed within each experimental group for DI measured under saline, Sb, and CNO. Treatment-associated change scores were calculated as drug DI minus saline DI and were similarly evaluated against CP. Reported p values for these exploratory cross-assay regressions are unadjusted and two-sided.

Because **Fig. S8** also asked whether CP could separate animals according to the corresponding social-discrimination response, exploratory within-sample thresholds were estimated using receiver-operating-characteristic analysis and Youden's index. For treatment-level analyses, the behavioral outcome was DI>0. For treatment-change analyses, the outcome reflected a positive drug-minus-saline DI change. The CP value maximizing Youden's (J) within each displayed dataset was used as the threshold.

Threshold performance was summarized descriptively using the estimated CP threshold, number and percentage of mice correctly classified, sensitivity, specificity, and confusion-matrix counts. These analyses use the same observations as the corresponding regressions in Fig. S8A and therefore represent within-sample characterization rather than independent prediction or external classifier validation. Full regression and classification results are provided in Table S8.

Supplementary Results

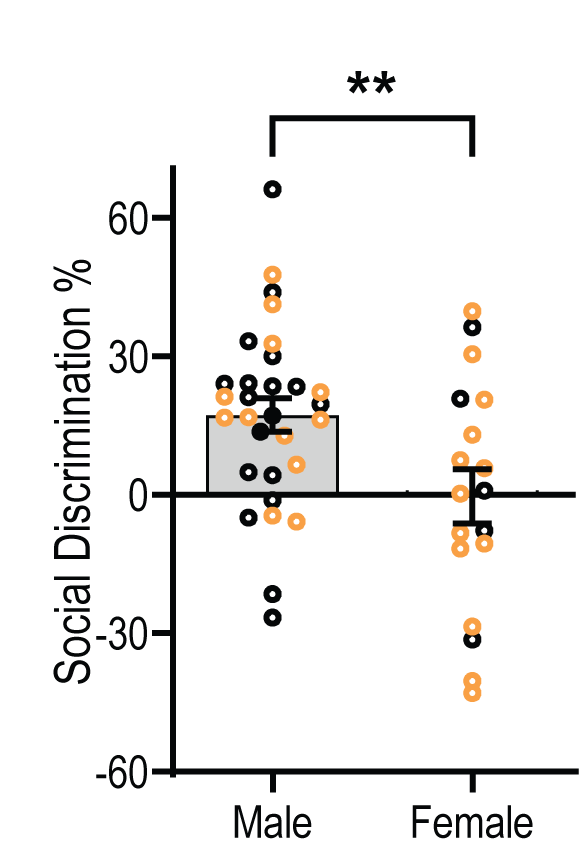

Fig. S1. Female control mice do not show reliable novel-directed social discrimination.
Social-discrimination index (DI) was compared between male and female non-Pilo mice to determine whether this social-memory task produced comparable baseline discrimination across sexes. Male and female mice differed significantly in DI (independent-samples t test, p<0.01). When each sex was tested separately against the chance value of DI=0 using one-sample t tests, male mice showed significant novel-directed discrimination (p<0.01), whereas female mice did not (*t*_(28)_=1.182, p=0.23). Thus, under the conditions used in this study, stable baseline social discrimination could not be established in female controls. Black circles represent control mice and orange circles represent control mice following stereotaxic surgery and viral transfection. Only saline-condition data are shown.

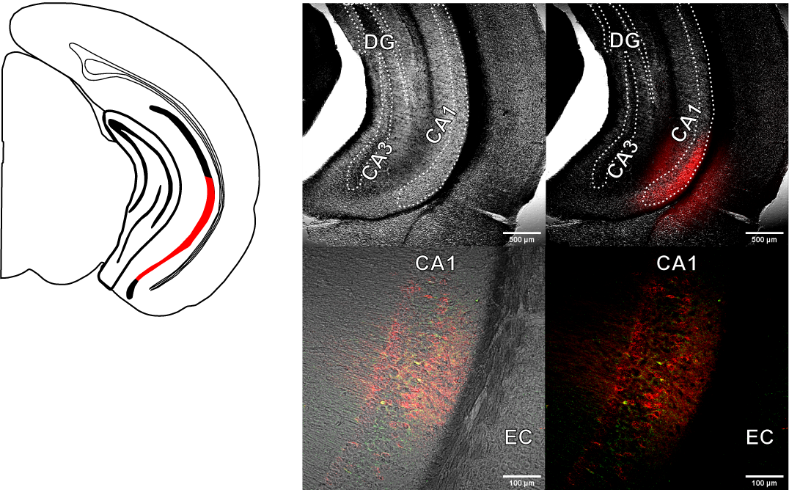

**Fig. S2.** **Viral infusion of DREADDs and KORD in vCA1/Sub.**
Proposed location of infusion (left) and appropriately targeted region in hippocampus of hM3Dq or hM4Di + mCherry and KORD+mCitrine (right).

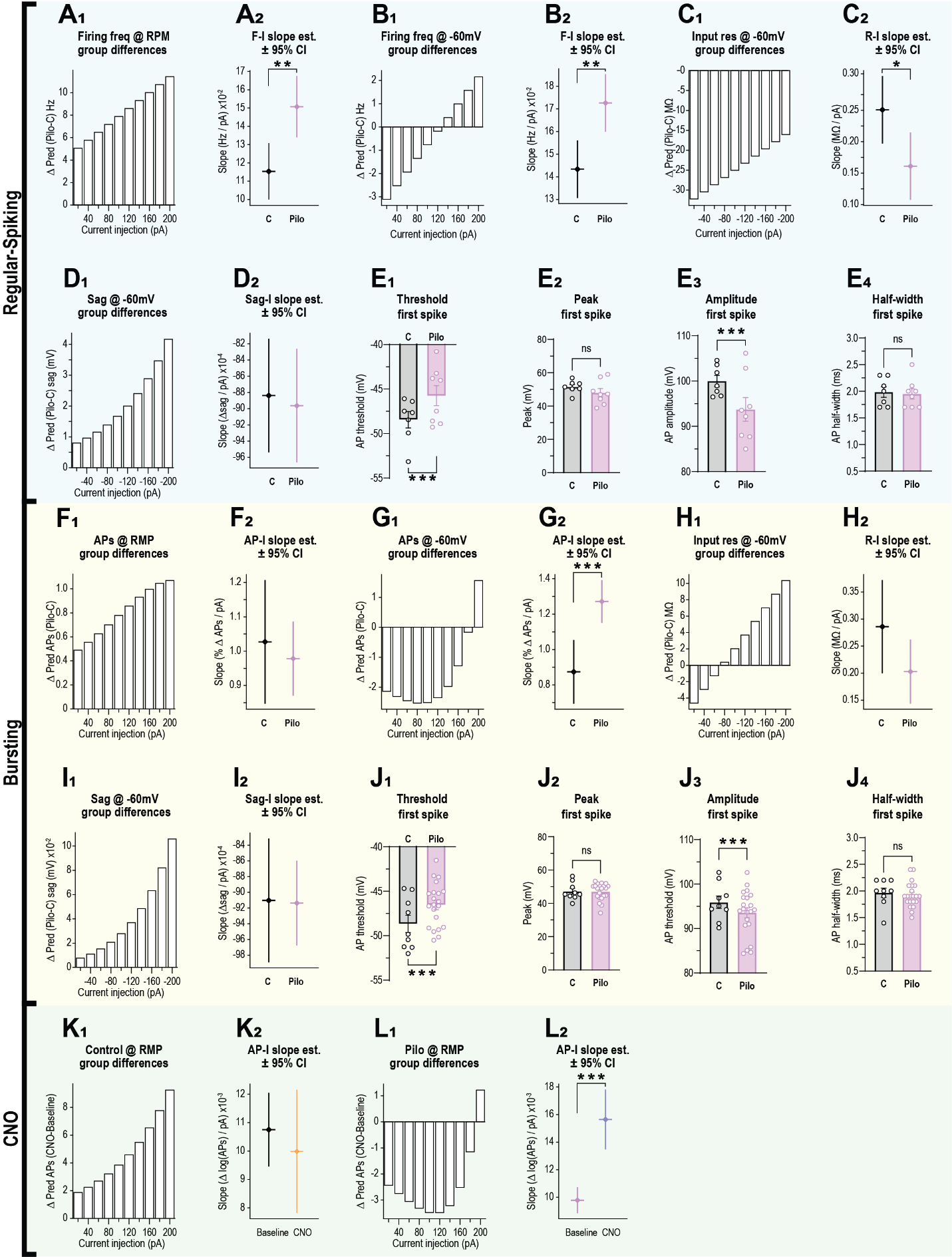

**Fig. S3.** **Changes in vCA1/Sub principal-neuron physiology in Pilo mice are often current dependent and accompany a more depolarized resting membrane potential.**
Most analyses were mixed linear models, and this figure includes depictions of interaction effects in both Δ(Pilo – Control) bar charts [_1_] and a slope estimate plot [_2_]. Main effects of group can be found in **Fig. 2**.

**Regular-spiking cells**. **A**. Firing frequency (F) was enhanced in Pilo mice compared to control when using RMP as the baseline (group x current). **B**. The effect was reduced but maintained when a constant membrane potential baseline was used (group x current) **C**. Input resistance (R) when held at -60mV baseline created an interaction (group x current) but was not different between groups. **D**. Pilo mice showed a trend toward larger sag deflection (mV) at hyperpolarizing current injection than control (group) but no significant interaction with current. **E**. Spike analysis of the first AP. The threshold to first spike was more depolarized in Pilo cells (group) (**E_1_**), the peak mV was not different (**E_2_**), yet the total amplitude was smaller in Pilo cells (group) (**E_3_**), the half-width was not different (**E_4_**). **Bursting cells**. **F**. There was no interaction effect for the number of action potentials between control and Pilo groups at RMP, (**G**) but a significant effect showed reduced number of APs in Pilo at lower current injection steps when the membrane was held at -60mV (group x current). **H**. Input resistance was not different. **I**. Sag was not different. **J**. First spike analysis revealed a lower threshold to AP in control mice compared to Pilo (group) (**J_1_**), but no differences in the peak (**J_2_**), or half-width (**J_4_**) between groups, yet amplitude was significantly but modestly reduced in Pilo (group) (**J_3_**). **Application of CNO**. **K**. Increased number of APs after CNO application to Control+DREADD mice did not create a significant interaction. **L**. Reduced number of APs during current injection steps in Pilo+DREADD mice following CNO application (group x current). Error bars indicate ±SEM. Group and interaction differences depicted with: *p*<0.05 *, *p*<0.01 **, *p*<0.001 ***

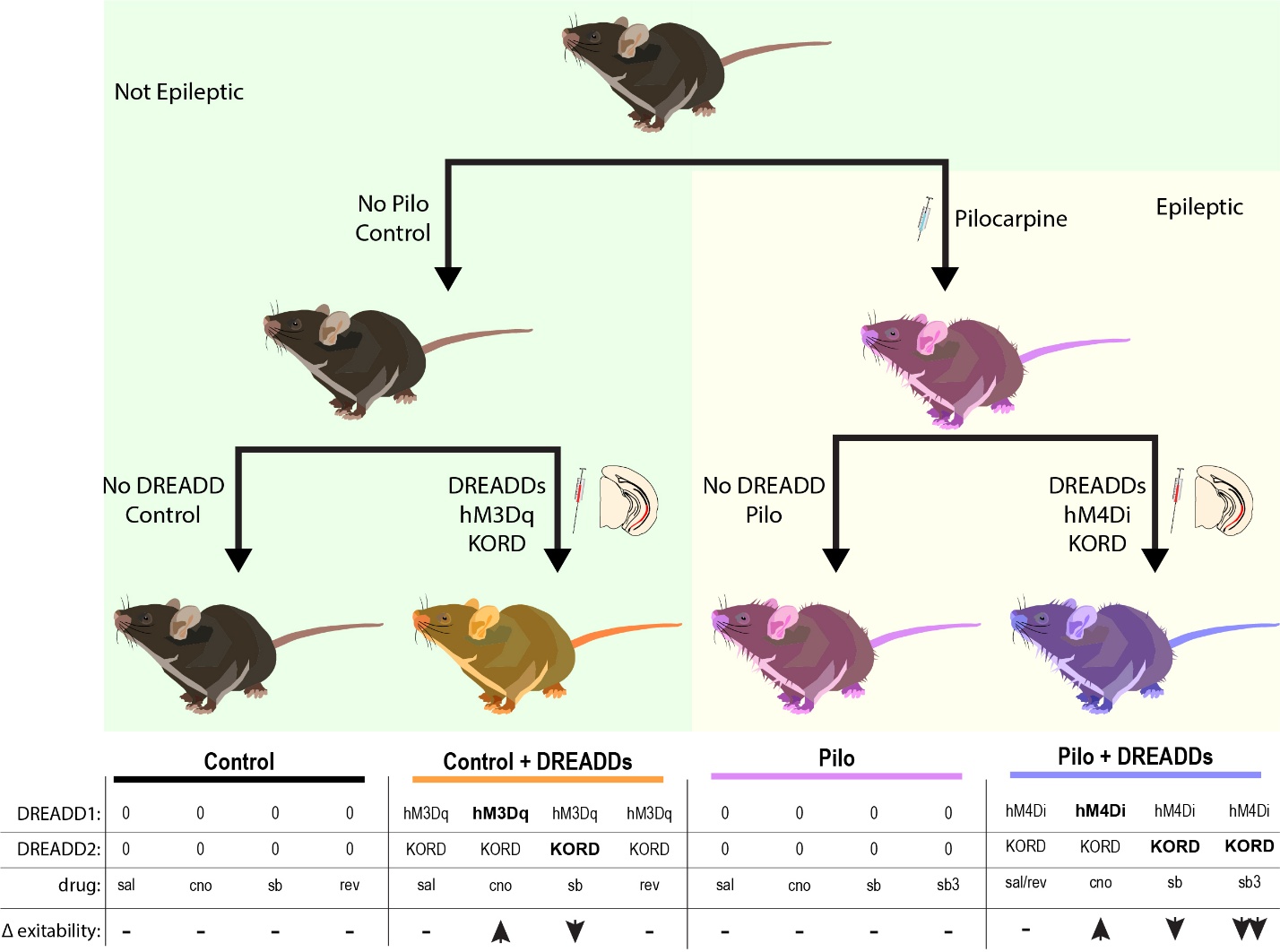

**Fig. S4.** **Group Diagram.**

Wild-type C57BL/6(E) mice were pseudorandomly assigned into one of four groups to create balanced cohorts for behavior. Mice assigned to Pilo or Pilo + DREADDs underwent pilocarpine injection at approximately 7-weeks of age per the listed protocol. Mice assigned to No DREADD control or Control + DREADD did not receive pilocarpine treatment. At approximately 8-9 weeks of age, mice assigned to the DREADD + groups underwent stereotaxic surgery. Specifically, non-pilo, control mice that received DREADDs received a mix of excitatory AAV5-CaMKIIα-hm3D(Gq)-mCherry (abbreviated “hM3Dq”) and inhibitory AAV5-CaMKIIα-HA-KORD-IRES-mCitrine (abbreviated “KORD”). Control group mice did not receive infusions. Pilo mice that received DREADDs received a mix of inhibitory AAV5-CaMKIIα-hM4D(Gi)-mCherry (abbreviated “hM4Di”) and inhibitory AAV5-CaMKIIα-HA-KORD-IRES-mCitrine. Pilo group mice did not receive infusions. The diagram illustrates the path mice are assigned to and the order of the operations to get them to the final groups. The bottom table lists the DREADD constructs that are present in each group divided into columns for the treatments they will receive in certain behavioral assays. Bolded DREADDs are those which should be activated by that given treatment/drug. Δ excitability is the proposed change to the excitability of the transfected cells for that treatment/drug (i.e. when a Control + DREADD mouse receives CNO, the excitability of the transfected vCA1/Sub pyramidal cells should increase from their baseline because only the excitatory hM3Dq and not the KORD is being activated).

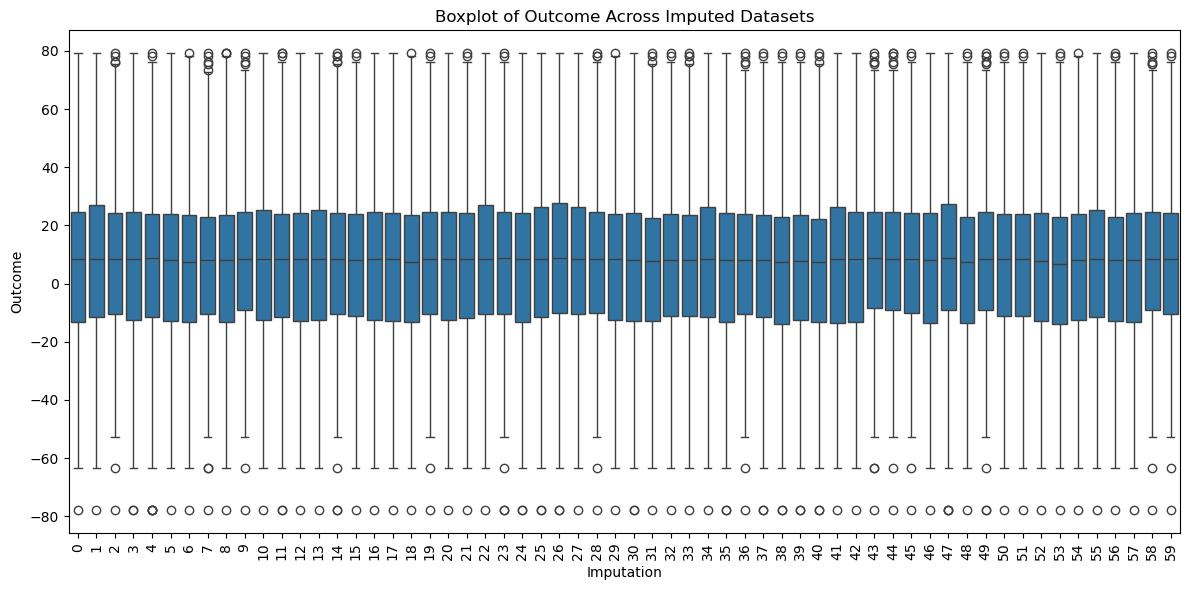

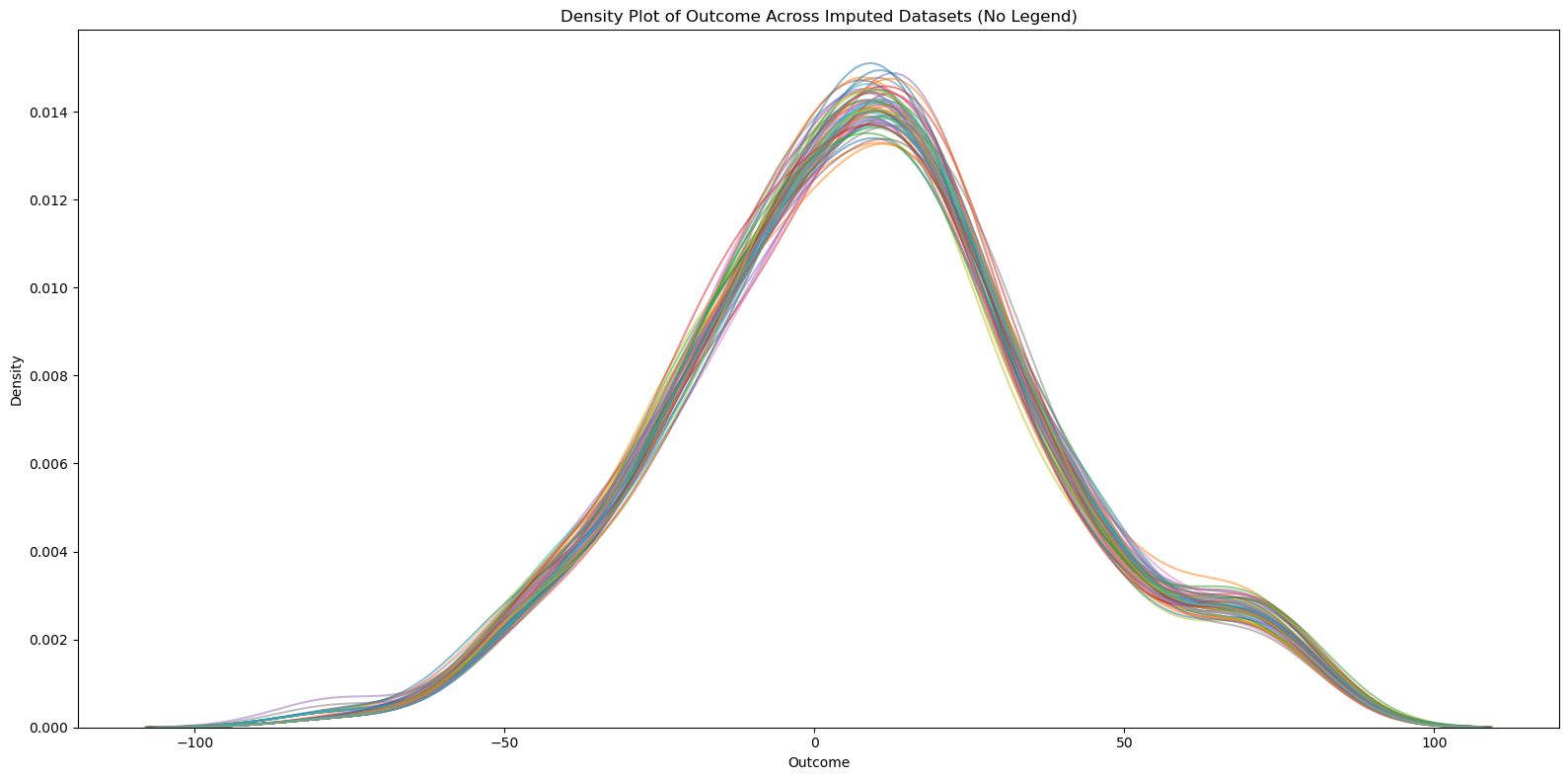

**Fig. S5.** **Imputation output boxplots and density plots.**

Boxplot of outcome (social discrimination score) distribution (quartiles + outliers) for all 60 imputations (top). Density plot of the outcome (social discrimination score) distribution, each line is a different imputation (bottom).

| Group | Treatment | Mean_Before | Variance_Before | Non-missing | Imputed | Ratio_Imputed | Mean_After | Variance_After |
| --- | --- | --- | --- | --- | --- | --- | --- | --- |
| Control+DREADD | CNO | 3.270481 | 399.4231 | 9 | 0 | 0 | 3.270481 | 355.7015 |
| Control+DREADD | Rev | 20.8137 | 275.7864 | 8 | 1 | 0.111111 | 19.51949 | 350.9097 |
| Control+DREADD | Saline | 16.06786 | 470.3821 | 9 | 0 | 0 | 16.06786 | 418.8931 |
| Control+DREADD | Sb | -1.08513 | 273.7978 | 8 | 1 | 0.111111 | -0.26816 | 301.1356 |
| Control/no-DREADD | CNO | 21.41725 | 88.12255 | 8 | 5 | 0.384615 | 16.39864 | 455.4885 |
| Control/no-DREADD | Rev | 6.93281 | 509.1545 | 8 | 5 | 0.384615 | 7.992773 | 628.9951 |
| Control/no-DREADD | Saline | 21.46361 | 502.8069 | 13 | 0 | 0 | 21.46361 | 464.7252 |
| Control/no-DREADD | Sb | 19.09388 | 315.9642 | 8 | 5 | 0.384615 | 13.79295 | 574.3839 |
| Pilo+DREADD | CNO | 9.297513 | 622.7662 | 15 | 1 | 0.0625 | 9.376214 | 593.6921 |
| Pilo+DREADD | Rev | 6.363111 | 812.5598 | 14 | 2 | 0.125 | 6.31622 | 768.3225 |
| Pilo+DREADD | Saline | -6.90933 | 568.7686 | 15 | 1 | 0.0625 | -6.25009 | 533.5504 |
| Pilo+DREADD | Sb | 11.03221 | 1324.674 | 14 | 2 | 0.125 | 11.12236 | 1164.768 |
| Pilo+DREADD | Sb3 | -10.6308 | 966.9488 | 15 | 1 | 0.0625 | -9.39583 | 928.2112 |
| Pilo/no-DREADD | CNO | -2.43448 | 1317.03 | 14 | 2 | 0.125 | -0.57612 | 1196.66 |
| Pilo/no-DREADD | Rev | 14.65098 | 1191.176 | 7 | 9 | 0.5625 | 10.23442 | 932.1901 |
| Pilo/no-DREADD | Saline | 13.81703 | 1491.44 | 16 | 0 | 0 | 13.81703 | 1399.683 |
| Pilo/no-DREADD | Sb | 9.941758 | 1381.221 | 15 | 1 | 0.0625 | 10.20485 | 1277.024 |

**Table S1. Imputation metrics by group and treatment**
Combined Statistics (Mean and Variance before and after Imputation). Total Original Values: 270, Ratio of Missing to Total: 0.1778, Percentage of Imputed Data: 17.78%.

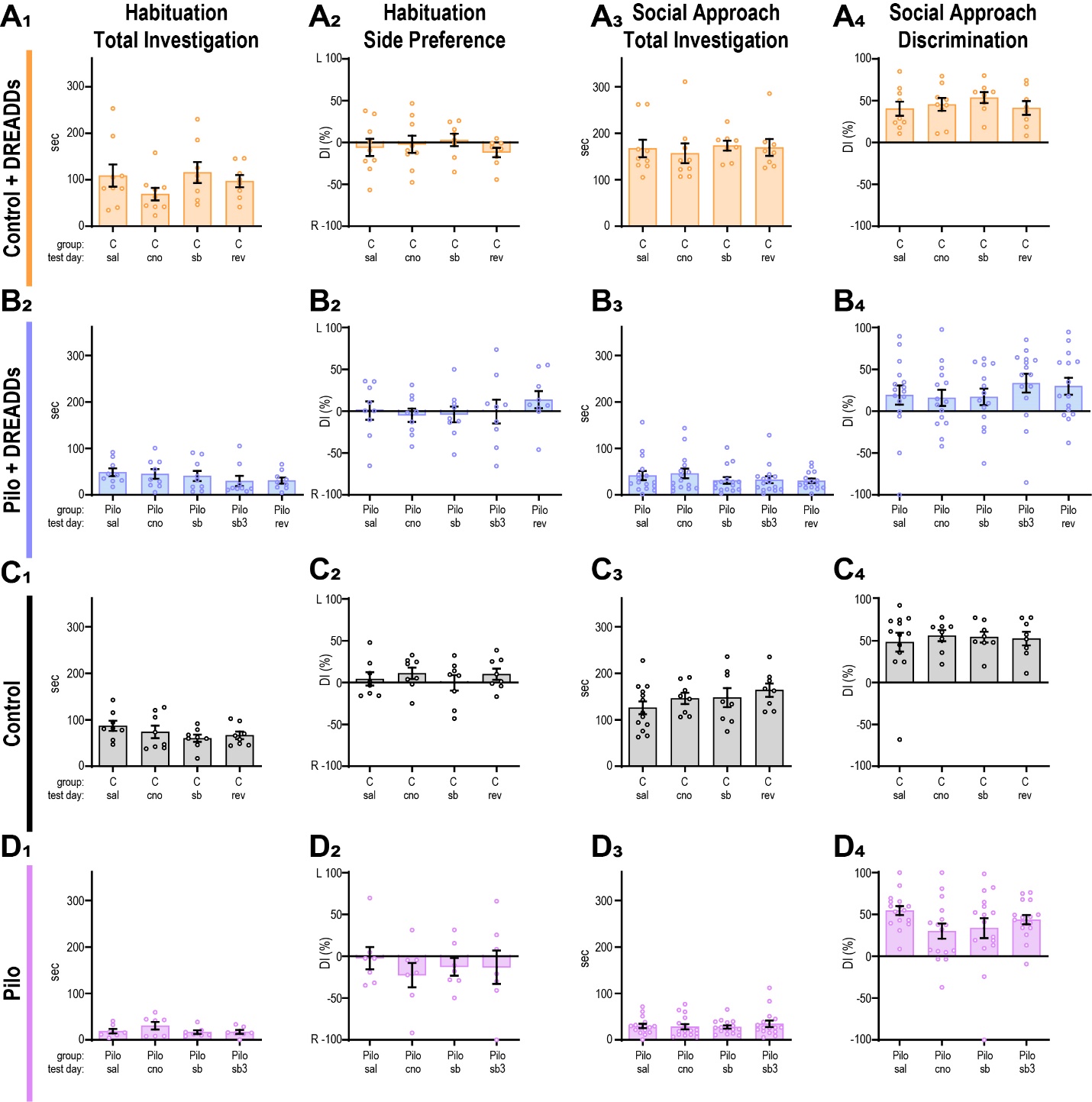

**Fig. S6.** **Mouse total investigation during habituation and social approach.**

Total time spent investigating Left or Right empty pencil cup in habituation and preference as a Discrimination index score (left two columns) and total investigation time during social approach (column 3) and social approach discrimination (column 4) for each group. These measures were prior to any treatment injection but sorted by day via the treatment type in independent columns within each graph: (C/Pilo), and test day (saline, CNO [1.5 mg/kg], Sb [1.5 mg/kg], and reversal saline; Sb3 [3.0 mg/kg] additionally for Pilo+DREADD mice).

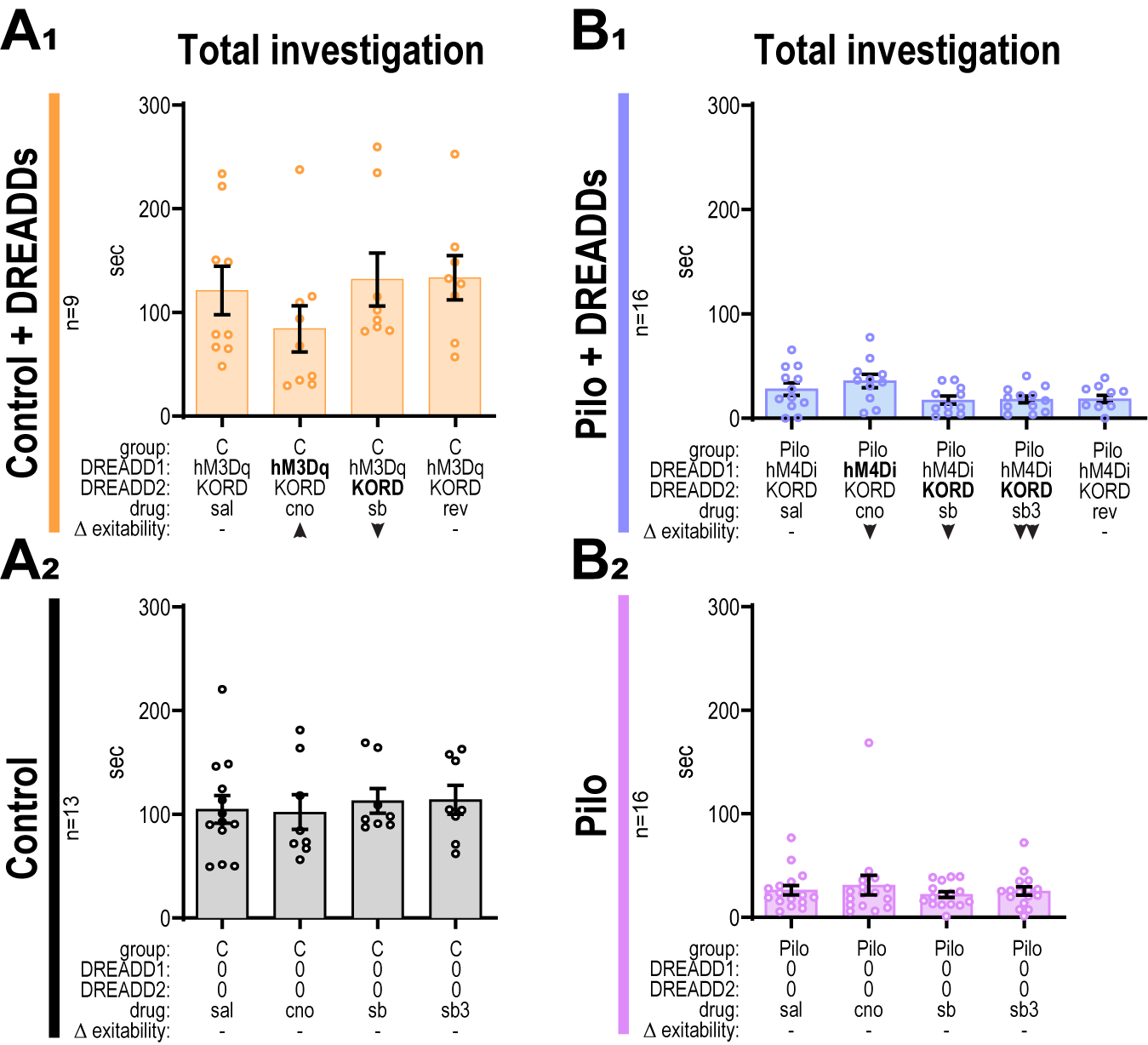

**Fig. S7.** **Mouse total investigation during social discrimination.**

Total time spent investigating both novel and familiar conspecific between groups for each treatment. Columns within each graph are specified by group (C/Pilo), DREADD type 1 (hM3Dq / hM4Di / none (0)), DREADD type 2 (KORD / 0), treatment/drug (saline, CNO [1.5 mg/kg], Sb [1.5 mg/kg], Sb3 [3.0 mg/kg] (for Pilo+DREADD mice), reversal [saline], and expected change in excitability ( - none, ▲ increased, ▼ decreased, ▼▼ substantially decreased). Bolded DREADD is the activated construct in a given treatment x group column. **A_1_**. Total investigation time is not different for C + DREADDs following treatments of saline, cno, salvinorin-b, and a reversal saline. **A_2_**. Total investigation time is not different for C following treatments of saline, cno, salvinorin-b, and a reversal saline. **B_1_**. Total investigation time is not different for Pilo + DREADDs following treatments of saline, cno, salvinorin-b, double salvinorin-b, and a reversal saline. **B_2_**. Total investigation time is not different for Pilo following treatments of saline, cno, salvinorin-b, and a reversal saline.

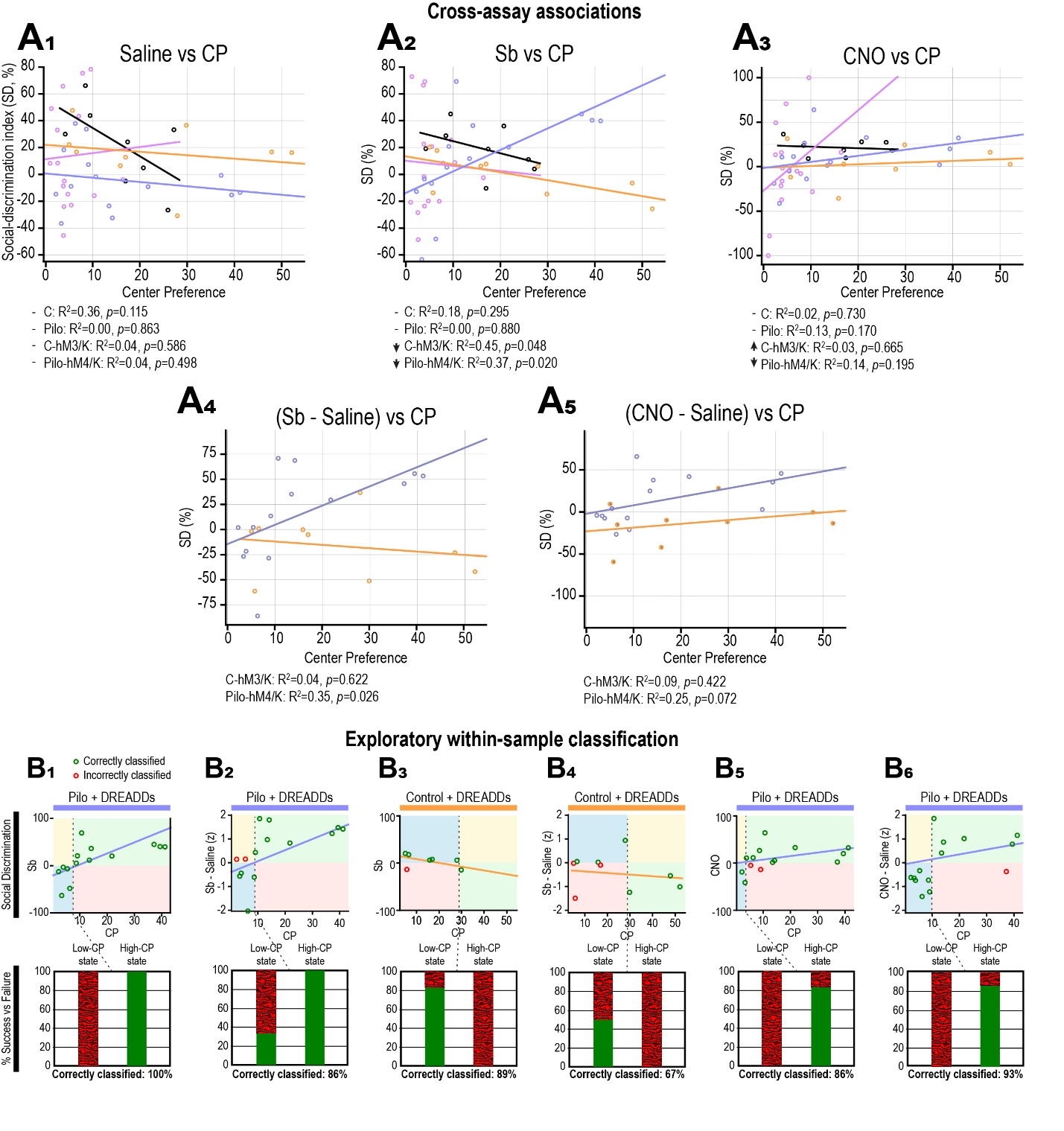
 **Fig. S8.** **Open-field center preference covaries with social-discrimination responses to vCA1/Sub modulation and supports exploratory within-sample classification.**

Center preference (CP; percentage of time spent in the center of the open field) was measured 1 h after salvinorin B (Sb) administration. Social-discrimination index (DI) was measured during separate social-memory sessions under saline, Sb, or CNO. All analyses used complete-case mouse pairs; imputed DI values were excluded. **A. Cross-assay associations.** Points represent individual mice, and lines show group-specific linear regressions. Black denotes Control/no-DREADD mice, magenta denotes Pilo/no-DREADD mice, orange denotes Control+DREADD mice, and blue denotes Pilo+DREADD mice. Model $R^{2}$and unadjusted two-sided *p* values are shown beneath each panel. **A1.** CP was not significantly associated with saline DI in any group, indicating that center preference did not generally predict baseline social-discrimination performance. **A2.** During Sb treatment, CP was not associated with DI in either no-DREADD group. In Control+DREADD mice, greater CP was associated with lower DI ($R^{2}=0.45$, *p*=0.048), whereas in Pilo+DREADD mice, greater CP was associated with higher DI ($R^{2}=0.37$, *p*=0.020). The opposing relationships are consistent with different behavioral consequences of KORD-mediated inhibition in healthy and epileptic circuits. **A3.** CP was not significantly associated with CNO DI in any group. **A4.** Association between CP and the Sb-related change in DI relative to saline. The relationship was not significant in Control+DREADD mice ($R^{2}=0.04$, *p*=0.622) but remained significant in Pilo+DREADD mice ($R^{2}=0.35$, *p*=0.026), indicating that greater center exploration covaried with a larger social-discrimination response to KORD-mediated inhibition. **A5.** Association between CP and the CNO-related change in DI relative to saline. No significant relationship was detected in Control+DREADD mice ($R^{2}=0.09$, *p*=0.422). Pilo+DREADD mice showed a similar positive trend that did not reach significance ($R^{2}=0.25$, *p*=0.072). **B. Exploratory within-sample classification.** The same complete-case observations were re-expressed using CP thresholds selected within each displayed dataset by Youden’s index. These analyses are exploratory and do not constitute independent validation of the associations in panel A. In the upper plots, the vertical dotted line indicates the estimated CP threshold and the horizontal line indicates the relevant DI or change-score boundary. Green-outlined points were correctly classified and red-outlined points were incorrectly classified. The lower plots show the observed proportions meeting or not meeting the behavioral criterion below and above the CP threshold. **B1.** In Pilo+DREADD mice tested with Sb, a CP threshold of 7.50 classified all 14 mice correctly according to whether DI was greater than 0 (within-sample accuracy, 100%).

**B2.** For the Sb-related change in DI in Pilo+DREADD mice, a CP threshold of 8.88 correctly classified 12 of 14 mice (85.7%). **B3.** In Control+DREADD mice tested with Sb, a CP threshold of 28.88 correctly classified 8 of 9 mice (88.9%). The direction of classification was opposite that observed in Pilo+DREADD mice, consistent with the negative association in A2.

**B4.** For the Sb-related change in DI in Control+DREADD mice, the same CP threshold correctly classified 6 of 9 mice (66.7%). **B5.** In Pilo+DREADD mice tested with CNO, a CP threshold of 3.63 correctly classified 12 of 14 mice according to whether DI was greater than 0 (85.7%).

**B6.** For the CNO-related change in DI in Pilo+DREADD mice, a CP threshold of 9.88 correctly classified 13 of 14 mice (92.9%).

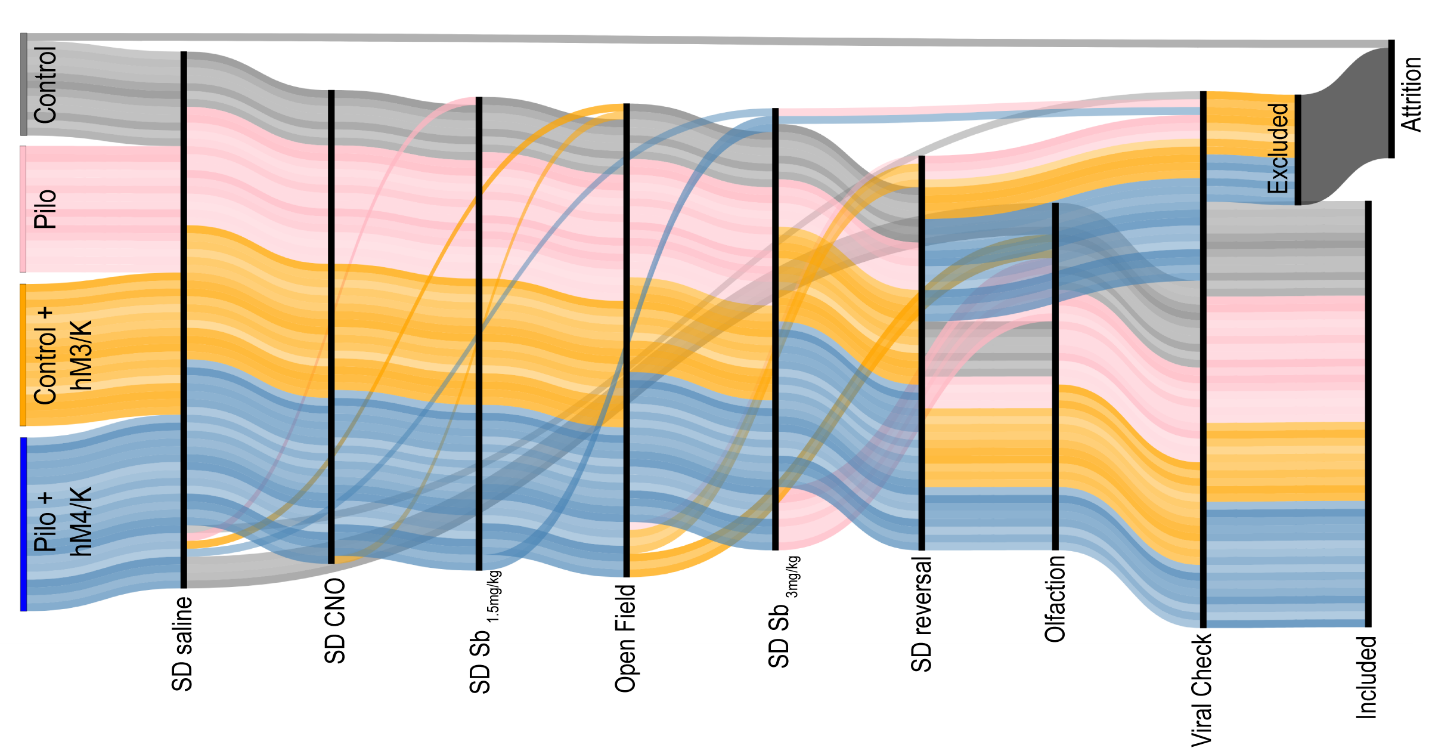

**Fig. S9.** **Sankey plot of mouse behavior.**

Mice are represented as individual lines colored within the four experimental groups. Though mice were pseudorandomly assigned an order for behaviors, some behaviors had higher attrition or had to occur later in the behavioral battery (social discrimination reversal has to follow other SD behaviors). Individual mice which did not participate in a given behavior are represented as curved lines emanating from the base of the behavioral column that they did participate in and skip the column(s) of those behaviors they did not participate in. Following all behavior a viral check was conducted to verify both the presence of the expressed fluorophores, and that the virus was not expressed in adjacent regions. Mice with appropriate viral expression were included in further analyses, while those without were excluded and counted amongst attrition.

| **Figure** | **Test Description** | **Statistical Test** | **Statistic** | **p-value** | **Effect Size** | **Directional Summary** |
| --- | --- | --- | --- | --- | --- | --- |
| 1B1 | Social Recognition: Preference for novel (Control) | one-sample t test vs DI=0 | t(12)=3.45 | < 0.01 | — | Control mice preferred novel conspecific over chance level |
| 1B1 | Social Recognition: Preference for novel (Pilo) | one-sample t test vs DI=0 | t(15)=1.43 | 0.17 | — | Pilo mice showed no preference above chance |
| 1B1 | Control vs Pilo DI | Independent t-test | t(27)=0.6314, | p=0.5331 | 95% CI [−32.50, 17.20] | No significant difference in continuous DI between groups. |
| 1C | Social Approach: Control (Stimulus vs Object) | one-sample t test vs DI=0 | t(12)=4.296 | 0.001 | — | Control mice preferred the stimulus mouse over the object |
| 1C | Social Approach: Pilo (Stimulus vs Object) | one-sample t test vs DI=0 | t(15)=10.23 | < 0.0001 | — | Pilo mice preferred the stimulus mouse over the object |
| 1D1 | Total investigation time (SD): Control vs Pilo | Independent t-test | t(26.51)=6.36 | < 0.0001 | η² = 0.60 | Pilo mice investigated less than controls in discrimination phase |
| 1D2 | Total interaction time (SA): Control vs Pilo | Independent t-test | t(14.62)=6.58 | < 0.0001 | η² = 0.75 | Pilo mice spent significantly less time investigating overall |
| 1E | Group × Stimulus interaction (LMM omnibus) | LMM (F-test from model) | F(2,84) = 9.34 | < 0.001 | — | Interaction significant: stimulus effect differs by group |
| 1E | Control: Opposite sex vs Vanilla | LMM (interaction term) | β = 19.86 | < 0.001 | — | Opposite sex scent investigated 4.96× longer than vanilla |
| 1E | Control: Opposite sex vs Water | LMM (interaction term) | β = 20.23 | < 0.001 | — | Opposite sex scent investigated 5.35× longer than water |
| 1E | Pilo: Opposite sex vs Vanilla | LMM (interaction term) | β = 6.44 | < 0.001 | — | Opposite sex scent investigated 3.2× longer than vanilla |
| 1E | Pilo: Opposite sex vs Water | LMM (interaction term) | β = 8.29 | < 0.001 | — | Opposite sex scent investigated 8.74× longer than water |

**Table S2.** **Group effect: Pilo mice had significantly reduced overall investigation times compared to controls (β = −15.52, p < 0.001).**

**Stimulus effect: Both vanilla and water scents were significantly less investigated than the opposite sex scent (vanilla: β = −19.86, p < 0.001; water: β = −20.23, p < 0.001).**
**Interaction effects: Pilo mice showed smaller differences between opposite sex and other stimuli than controls (Pilo × vanilla: β = 13.42, p < 0.001; Pilo × water: β = 11.94, p < 0.001).**

| **Figure** | **Test Description** | **Statistical Test** | **Statistic** | **p-value** | **Transformation** | **Directional Summary** |
| --- | --- | --- | --- | --- | --- | --- |
| 2B | RMP: Control vs Pilo (RS) | LMM | β = 4.71, z = 2.14 | 0.033 | None | Pilo cells more depolarized (~4.7 mV) |
| S3A | AP Frequency at RMP: Group × Current (RS) | LMM | β = 0.035, z = 3.10 | 0.002 | None | Firing rate increases more steeply in Pilo |
| 2D1 | AP Frequency at RMP: Group | LMM | β = 4.35, z = 0.84 | 0.402 | None | No baseline difference between groups |
| 2D1 | AP Frequency at RMP: Current | LMM | β = 0.115, z = 14.87 | <0.001 | None | Firing rate increases with current injection |
| S3B | AP Frequency at −60 mV: Group × Current (RS) | LMM | β = 0.029, z = 3.06 | 0.002 | None | Firing rate increases more steeply in Pilo at −60 mV |
| 2D2 | AP Frequency at −60 mV: Group (RS) | LMM | β = −3.97, z = −1.14 | 0.255 | None | No baseline group difference |
| 2D2 | AP Frequency at −60 mV: Current (RS) | LMM | β = 0.143, z = 21.20 | <0.001 | None | Firing rate strongly increases with current |
| S3C | Input Resistance: Group × Current (RS) | LMM | β = −0.090, z = −2.37 | 0.018 | None | Pilo cells show a shallower R–I slope (reduced sensitivity) |
| 2E | Input Resistance: Group (RS) | LMM | β = −33.89, z = −1.36 | 0.174 | None | No baseline group difference |
| 2E | Input Resistance: Current (RS) | LMM | β = 0.251, z = 9.35 | <0.001 | None | Input resistance was lower at stronger hyperpolarizing current steps |
| S3D | Sag Voltage (RS): Group × Current | LMM | β = −0.000, z = −0.15 | 0.880 | Log-transformed | No interaction, slopes parallel |
| 2F | Sag Voltage (RS): Group | LMM | β = 0.522, z = 1.95 | 0.051 | Log-transformed | Trend toward greater sag in Pilo (~1–5 mV higher) |
| 2F | Sag Voltage (RS): Current | LMM | β = −0.009, z = −15.11 | <0.001 | Log-transformed | Sag decreases with less hyperpolarization |
| S3E1 | Threshold to Spike (RS) | LMM | β = −0.057, z = −3.32 | 0.001 | Log-transformed | More depolarized spike threshold in Pilo. |
| S3E2 | Peak Voltage (RS) | LMM | β = −3.56, z = −1.21 | 0.227 | None | No difference |
| S3E3 | Spike Amplitude (RS) | LMM | β = −6.26, z = −3.24 | 0.001 | None | Reduced amplitude in Pilo |
| S3E4 | Half-width (RS) | LMM | β = −0.012, z = −0.28 | 0.783 | Log-transformed | No difference |
| 2G | RMP: Control vs Pilo (Bursting) | LMM | β = 3.33, z = 2.37 | 0.018 | None | Pilo cells more depolarized (~3.3 mV) |
| S3F | APs at RMP: Group × Current (Bursting) | LMM | β = −0.000, z = −0.46 | 0.647 | Log-transformed | No interaction, groups have parallel slopes |
| 2I1 | APs at RMP: Group (Bursting) | LMM | β = 0.146, z = 0.74 | 0.460 | Log-transformed | No difference |
| 2I1 | APs at RMP: Current (Bursting) | LMM | β = 0.010, z = 11.34 | <0.001 | Log-transformed | Strong current–firing relationship (+~10% APs per +10 pA) |
| S3G | APs at −60 mV: Group × Current (Bursting) | LMM | β = 0.004, z = 3.67 | <0.001 | Log-transformed | Pilo cells show steeper gain with current |
| 2I2 | APs at −60 mV: Group (Bursting) | LMM | β = −0.718, z = −3.39 | 0.001 | Log-transformed | Pilo cells fire fewer APs overall |
| 2I2 | APs at −60 mV: Current (Bursting) | LMM | β = 0.009, z = 9.74 | <0.001 | Log-transformed | Firing strongly increases with current |
| S3H | Input Resistance: Group × Current (Bursting) | LMM | β = −0.083, z = −1.58 | 0.114 | None | No significant interaction |
| 2J | Input Resistance: Group (Bursting) | LMM | β = −6.28, z = −0.26 | 0.794 | None | No overall group difference |
| 2J | Input Resistance: Current (Bursting) | LMM | β = 0.286, z = 6.58 | <0.001 | None | Input resistance increases with current step |
| S3I | Sag Amplitude: Group × Current (Bursting) | LMM | β = −0.000, z = −0.07 | 0.943 | None | No interaction; slopes identical |
| 2K | Sag Amplitude: Group (Bursting) | LMM | β = 0.003, z = 0.02 | 0.987 | None | No overall group difference |
| 2K | Sag Amplitude: Current (Bursting) | LMM | β = −0.009, z = −22.87 | <0.001 | None | Sag increases with stronger hyperpolarization |
| S3J1 | Threshold to Spike (Bursting) | LMM | β = 2.14, z = 41.58 | <0.001 | None | Higher threshold in Pilo |
| S3J2 | Peak Voltage (Bursting) | LMM | β = −0.005, z = −0.28 | 0.784 | Log-transformed | No difference |
| S3J3 | Spike Amplitude (Bursting) | LMM | β = −2.35, z = −9.31 | <0.001 | None | Reduced amplitude in Pilo |
| S3J4 | Half-width (Bursting) | LMM | β = −0.014, z = −0.58 | 0.561 | Log-transformed | No difference |
| S3K | AP Count: CNO × Current (Control hM3Dq) | LMM | β = −0.001, z = −0.60 | 0.550 | Log-transformed | No interaction; slope unaffected by CNO |
| 2N | AP Count: CNO main effect (Control hM3Dq) | LMM | β = +0.608, z = 3.83 | <0.001 | Log-transformed | CNO depolarizes control neurons (~+2 to +9 APs across current steps) |
| 2N | AP Count: Current effect (Control hM3Dq) | LMM | β = +0.011, z = 16.51 | <0.001 | Log-transformed | APs increase strongly with depolarizing current |
| 2M | DREADD RMP change (Control hM3Dq) | Paired t-test | t(5)=4.988 | 0.0041 | None | More depolarized RMP after CNO in control |
| S3L | AP Count: CNO × Current (Pilo hM4Di) | LMM | β = +0.006, z = 4.97 | <0.001 | Log-transformed | CNO altered the current–firing relationship; suppression was greatest at lower current steps and diminished as current increased. |
| 2P | AP Count: CNO main effect (Pilo hM4Di) | LMM | β = −1.121, z = −6.16 | <0.001 | Log-transformed | CNO reduced firing, with the clearest suppression at lower current steps |
| 2P | AP Count: Current effect (Pilo hM4Di) | LMM | β = +0.010, z = 21.22 | <0.001 | Log-transformed | Firing increased strongly with depolarizing current under both conditions. |
| 2O | DREADD RMP change (Pilo hM4Di) | Paired t-test | t(3)=11.09 | 0.0016 | None | More hyperpolarized RMP after CNO in Pilo |

**Table S3.** Statistical details associated with figure 2 and supplemental figure 3

| **Figure** | **Test Description** | **Statistical Test** | **Statistic** | **p-value** | **Effect Size** | **Directional Summary** |
| --- | --- | --- | --- | --- | --- | --- |
| 3C | PV Density: Full ROI (Control vs Pilo) | Independent t-test | t(19) = 2.685 | 0.0146 | η² = 0.275 | 28% reduction in Pilo mice |
| 3C | PV Density: Stratum oriens (Control vs Pilo) | Independent t-test | t(19) = 3.170 | 0.005 | η² = 0.346 | 45% reduction in Pilo mice |
| 3C | PV Density: Stratum pyramidale (Control vs Pilo) | Independent t-test | t(19) = 2.004 | 0.0595 | η² = 0.175 | Trend toward reduction |
| 3C | PV Density: Stratum radiatum (Control vs Pilo) | Welch-corrected t-test | t(13.39) = 0.888 | 0.39 | η² = 0.056 | No difference |
| 3D | SST Density: Full ROI (Control vs Pilo) | Independent t-test | t(12) = 0.972 | 0.3502 | η² = 0.073 | No difference |
| 3D | SST Density: Stratum oriens (Control vs Pilo) | Independent t-test | t(12) = 2.304 | 0.0399 | η² = 0.307 | 32% reduction in Pilo mice |
| 3D | SST Density: Stratum pyramidale (Control vs Pilo) | Independent t-test | t(12) = 0.151 | 0.8824 | η² = 0.002 | No difference |
| 3D | SST Density: Stratum radiatum (Control vs Pilo) | Welch-corrected t-test | t(7.57) = 0.775 | 0.4617 | η² = 0.074 | No difference |

**Table S4.** Statistical details associated with figure 3.

| **Figure** | **Test Description** | **Statistical Test** | **Statistic** | **p-value** | **Effect Size** | **Directional Summary** |
| --- | --- | --- | --- | --- | --- | --- |
| 4G | AP Probability: Control vs Pilo | Marginal Cox model | HR = 0.642 (95% CI: 0.224–1.837) | 0.353 | — | No difference in AP generation probability across groups |
| 4F | Hyperpolarization Amplitude: Stim × Group | LMM (Box-Cox) | β = 0.997 (Pilo = 0.256), z = 3.89 | < 0.001 | pseudo-R² = 0.726, f² = 2.65 | Control cells show stronger hyperpolarization, especially at higher stimulus |
| 4F | Hyperpolarization voltage-time integral: Stim × Group | LMM (Yeo-Johnson) | β = 0.060 (Pilo = 0.020), z = 2.98 | 0.003 | pseudo-R² = 0.2281, f² = 0.296 | Hyperpolarization voltage-time integral larger in control; moderate effect |
| — | Amplitude vs voltage-time integral: Hyperpolarizing | Pearson correlation | r = 0.500 | — | — | Amplitude measures moderately correlated, no singularity |
| 4F | Depolarization Amplitude: Stim × Group | LMM (Yeo-Johnson) | β = −0.027 (Pilo = 0.009), z = −3.00 | 0.004 | pseudo-R² = 0.2299, f² = 0.299 | Pilo cells show slightly reduced depolarization amplitude |
| 4F | Depolarization voltage-time integral: Stim main effect | LMM (Yeo-Johnson) | β = 0.110 (Pilo = 0.008), z = 13.75 | < 0.001 | pseudo-R² = 0.2061, f² = 0.260 | Depolarization voltage-time integral increases with stimulus, no group difference |
| — | Amplitude vs voltage-time integral: Depolarizing | Pearson correlation | r = 0.612 | — | — | voltage-time integral measures moderately correlated, no singularity |

**Table S5.** Statistical details associated with figure 4.

| **Figure** | **Test Description** | **Statistical Test** | **Statistic** | **p-value** | **Effect Size / Notes** | **Directional Summary** |
| --- | --- | --- | --- | --- | --- | --- |
| 5F1 | Line length: Treatment × epoch | Likelihood-ratio test, full vs additive LMM | χ²(2) = 0.57 | 0.751 | Planned omnibus temporal test; unadjusted | No evidence that the CNO–saline difference varied across epochs. |
| 5F1 | Line length: CNO vs saline, averaged across all post-injection epochs | Planned contrast from LMM | β = -0.159; SE = 0.031; z = -5.14; 95% CI [-0.220, -0.099] | 2.70e-07 | Holm p = 8.10e-07; Holm family: target engagement, time-averaged | Line length was lower after CNO across the post-injection period. |
| 5F1 | Line length: CNO vs saline, epoch 1 (behavior-aligned) | Planned contrast from LMM | β = -0.146; SE = 0.037; z = -3.99; 95% CI [-0.218, -0.074] | 6.64e-05 | Holm p < 0.001; Holm family: target engagement, epoch 1 | Line length was already lower during the behavior-aligned epoch. |
| 5F2 | Delta power: Treatment × epoch | Likelihood-ratio test, full vs additive LMM | χ²(2) = 5.76 | 0.056 | Planned omnibus temporal test; unadjusted | The temporal interaction approached but did not reach significance. |
| 5F2 | Delta power: CNO vs saline, averaged across all post-injection epochs | Planned contrast from LMM | β = -0.290; SE = 0.066; z = -4.40; 95% CI [-0.419, -0.161] | 1.07e-05 | Holm p = 1.19e-05; Holm family: target engagement, time-averaged | Delta power was lower after CNO across the post-injection period. |
| 5F2 | Delta power: CNO vs saline, epoch 1 (behavior-aligned) | Planned contrast from LMM | β = -0.195; SE = 0.083; z = -2.35; 95% CI [-0.358, -0.032] | 0.019 | Holm p = 0.019; Holm family: target engagement, epoch 1 | Delta power was lower during the behavior-aligned epoch. |
| 5F3 | Gamma power: Treatment × epoch | Likelihood-ratio test, full vs additive LMM | χ²(2) = 1.64 | 0.441 | Planned omnibus temporal test; unadjusted | No evidence that the CNO–saline difference varied across epochs. |
| 5F3 | Gamma power: CNO vs saline, averaged across all post-injection epochs | Planned contrast from LMM | β = -0.340; SE = 0.075; z = -4.53; 95% CI [-0.488, -0.193] | 5.97e-06 | Holm p = 1.19e-05; Holm family: target engagement, time-averaged | Gamma power was lower after CNO across the post-injection period. |
| 5F3 | Gamma power: CNO vs saline, epoch 1 (behavior-aligned) | Planned contrast from LMM | β = -0.286; SE = 0.087; z = -3.28; 95% CI [-0.457, -0.115] | 0.001 | Holm p = 0.002; Holm family: target engagement, epoch 1 | Gamma power was lower during the behavior-aligned epoch. |
| 5H1 | Composite spike frequency: Treatment × epoch | Likelihood-ratio test, full vs additive LMM | χ²(2) = 4.38 | 0.112 | Planned omnibus temporal test; unadjusted | The overall treatment effect did not significantly vary by epoch. |
| 5H1 | Composite spike frequency: CNO vs saline, epoch 1 | Planned contrast from LMM | β = -0.193; SE = 0.179; z = -1.08; 95% CI [-0.544, 0.158] | 0.281 | Holm p = 0.282 (temporal family); Holm p = 0.562 (epoch-1 spike family); Epoch 1 is the behavior-aligned interval | No detectable early reduction in interictal spike frequency. |
| 5H1 | Composite spike frequency: CNO vs saline, epoch 2 | Planned contrast from LMM | β = -0.411; SE = 0.179; z = -2.30; 95% CI [-0.762, -0.060] | 0.022 | Holm p = 0.043; Holm family: spike-frequency temporal follow-up | Spike frequency was lower after CNO in epoch 2. |
| 5H1 | Composite spike frequency: CNO vs saline, epoch 3 | Planned contrast from LMM | β = -0.512; SE = 0.179; z = -2.86; 95% CI [-0.863, -0.161] | 0.004 | Holm p = 0.013; Holm family: spike-frequency temporal follow-up | Spike frequency was lower after CNO in epoch 3. |
| 5H2 | Composite spike amplitude: Treatment × epoch | Likelihood-ratio test, full vs additive LMM | χ²(2) = 0.92 | 0.631 | Planned omnibus temporal test; unadjusted | No evidence that the CNO–saline difference varied across epochs. |
| 5H2 | Composite spike amplitude: CNO vs saline, epoch 1 | Planned contrast from LMM | β = -0.033; SE = 0.035; z = -0.96; 95% CI [-0.101, 0.035] | 0.337 | Holm p = 0.562; Holm family: epoch-1 spike outcomes | No detectable change in spike amplitude during the behavior-aligned epoch. |
| 5G | Overt seizure count: CNO vs saline during the 4-h target window | Poisson GLM with log(expected count) offset and HC0 robust SE | log β = 0.305; SE = 0.304; z = 1.00; RR = 1.36; 95% CI [0.75, 2.46] | 0.316 | Single prespecified test; n = 26 sessions; Pearson χ²/df = 1.14 | CNO did not detectably reduce overt seizure counts. |
| S. analysis | Cluster-adjusted line length: Treatment × epoch | Likelihood-ratio test, full vs additive LMM | χ²(2) = 1.96 | 0.376 | Matched non-injection reference; unadjusted omnibus test | No evidence that the adjusted treatment difference varied across epochs. |
| S. analysis | Cluster-adjusted line length: CNO vs saline, averaged across epochs | Planned contrast from LMM | β = -0.147; SE = 0.057; z = -2.59; 95% CI [-0.258, -0.036] | 0.010 | Holm p = 0.019; Holm family: adjusted target engagement, time-averaged | The time-averaged CNO reduction persisted after cluster adjustment. |
| S. analysis | Cluster-adjusted line length: CNO vs saline, epoch 1 | Planned contrast from LMM | β = -0.120; SE = 0.060; z = -1.99; 95% CI [-0.238, -0.002] | 0.046 | Holm p = 0.139; Holm family: adjusted target engagement, epoch 1 | Direction was consistent, but the adjusted epoch-1 contrast was not significant after correction. |
| S. analysis | Cluster-adjusted delta power: Treatment × epoch | Likelihood-ratio test, full vs additive LMM | χ²(2) = 3.20 | 0.202 | Matched non-injection reference; unadjusted omnibus test | No significant adjusted treatment-by-epoch interaction. |
| S. analysis | Cluster-adjusted delta power: CNO vs saline, averaged across epochs | Planned contrast from LMM | β = -0.284; SE = 0.101; z = -2.81; 95% CI [-0.481, -0.086] | 0.005 | Holm p = 0.015; Holm family: adjusted target engagement, time-averaged | The time-averaged CNO reduction persisted after cluster adjustment. |
| S. analysis | Cluster-adjusted delta power: CNO vs saline, epoch 1 | Planned contrast from LMM | β = -0.198; SE = 0.117; z = -1.70; 95% CI [-0.427, 0.030] | 0.089 | Holm p = 0.179; Holm family: adjusted target engagement, epoch 1 | Direction was consistent, but the adjusted epoch-1 contrast was not significant. |
| S. analysis | Cluster-adjusted gamma power: Treatment × epoch | Likelihood-ratio test, full vs additive LMM | χ²(2) = 3.05 | 0.218 | Matched non-injection reference; unadjusted omnibus test | No significant adjusted treatment-by-epoch interaction. |
| S. analysis | Cluster-adjusted gamma power: CNO vs saline, averaged across epochs | Planned contrast from LMM | β = -0.298; SE = 0.129; z = -2.31; 95% CI [-0.551, -0.045] | 0.021 | Holm p = 0.021; Holm family: adjusted target engagement, time-averaged | The time-averaged CNO reduction persisted after cluster adjustment. |
| S. analysis | Cluster-adjusted gamma power: CNO vs saline, epoch 1 | Planned contrast from LMM | β = -0.231; SE = 0.137; z = -1.69; 95% CI [-0.499, 0.037] | 0.091 | Holm p = 0.179; Holm family: adjusted target engagement, epoch 1 | Direction was consistent, but the adjusted epoch-1 contrast was not significant. |
| S. analysis | Cluster-adjusted composite spike frequency: Treatment × epoch | Likelihood-ratio test, full vs additive LMM | χ²(2) = 1.73 | 0.421 | Matched non-injection reference; unadjusted omnibus test | No significant adjusted treatment-by-epoch interaction. |
| S. analysis | Cluster-adjusted composite spike frequency: CNO vs saline, epoch 1 | Planned contrast from LMM | β = -0.349; SE = 0.348; z = -1.00; 95% CI [-1.030, 0.333] | 0.316 | Holm p = 0.633 (temporal family); Holm p = 0.572 (epoch-1 spike family); Matched non-injection reference | No detectable adjusted spike-frequency reduction in epoch 1. |
| S. analysis | Cluster-adjusted composite spike frequency: CNO vs saline, epoch 2 | Planned contrast from LMM | β = -0.505; SE = 0.348; z = -1.45; 95% CI [-1.187, 0.177] | 0.147 | Holm p = 0.440; Holm family: adjusted spike-frequency temporal follow-up | The later active-treatment contrast was not retained after cluster adjustment. |
| S. analysis | Cluster-adjusted composite spike frequency: CNO vs saline, epoch 3 | Planned contrast from LMM | β = -0.305; SE = 0.348; z = -0.88; 95% CI [-0.987, 0.377] | 0.380 | Holm p = 0.633; Holm family: adjusted spike-frequency temporal follow-up | No detectable adjusted spike-frequency difference in epoch 3. |
| S. analysis | Cluster-adjusted composite spike amplitude: Treatment × epoch | Likelihood-ratio test, full vs additive LMM | χ²(2) = 4.07 | 0.130 | Matched non-injection reference; unadjusted omnibus test | No significant adjusted treatment-by-epoch interaction. |
| S. analysis | Cluster-adjusted composite spike amplitude: CNO vs saline, epoch 1 | Planned contrast from LMM | β = -0.042; SE = 0.039; z = -1.07; 95% CI [-0.119, 0.035] | 0.286 | Holm p = 0.572; Holm family: adjusted epoch-1 spike outcomes | No detectable adjusted change in spike amplitude. |
| S. analysis | Overt seizure count sensitivity: CNO vs saline after excluding borderline cluster-timing sessions | Poisson GLM with log(expected count) offset and HC0 robust SE | log β = -0.028; SE = 0.270; z = -0.10; RR = 0.97; 95% CI [0.57, 1.65] | 0.916 | Single sensitivity test; n = 24 sessions; Pearson χ²/df = 1.02 | The conclusion of no detectable seizure reduction was unchanged. |

**Table S6.** Statistical details associated with Figure 5 and cluster-reference sensitivity analyses.
Notes: Active EEG analyses included 84 observations from five mice (45 CNO, 39 saline). Cluster-adjusted sensitivity analyses used the same active observations after subtraction of the matched non-injection mean from the same mouse, seizure cluster, and epoch. LMMs included treatment, epoch, treatment-by-epoch interaction, and injection day as fixed effects, with a random mouse intercept and variance components for cluster and session. β is the model-estimated CNO-minus-saline difference. Planned contrasts were Holm-adjusted within the listed families; likelihood-ratio interaction tests were unadjusted. Epochs were 45 min–2 h 45 min, 2 h 45 min–4 h 45 min, and 4 h 45 min–6 h 45 min. Seizure models used n=26 primary and n=24 sensitivity sessions and included log(expected count) as an offset.

| **Figure** | **Test Description** | **Statistical Test** | **Statistic** | **p-value** | **Effect Size / Notes** | **Directional Summary** |
| --- | --- | --- | --- | --- | --- | --- |
| 6B1 | Average Speed: Across 4 groups | Welch’s ANOVA | F(3, 23.16) = 3.437 | 0.0335 | Bartlett’s χ²(3) = 9.25, p = 0.0261 | Significant group differences |
| 6B1 | Avg Speed: Control/no-DREADD vs Control+DREADD | Dunnett’s T3 | Δ = +0.02283 m/s, t(13.26) = 2.845 | 0.0394 | 95% CI: [0.0010, 0.0446] | Speed increased in Control+DREADD mice |
| 6B1 | Avg Speed: Control/no-DREADD vs Pilo/no-DREADD | Dunnett’s T3 | Δ = −0.01589 m/s, t(19.84) = 1.762 | 0.2475 | 95% CI: [−0.0393, 0.0075] | No significant difference |
| 6B1 | Avg Speed: Pilo/no-DREADD vs Pilo+DREADD | Dunnett’s T3 | Δ = −0.00171 m/s, t(27.20) = 0.141 | 0.9986 | 95% CI: [−0.0327, 0.0293] | No effect of KORD in Pilo |
| Not shown | Distance traveled | Welch’s ANOVA | F(3,21.37) = 2.908 | 0.0581 | Bartlett’s χ²(3) = 9.16, p = 0.0272 | Near significant group differences; Not Shown |
| 6B2 | Max Speed: Across 4 groups | Welch’s ANOVA | F(3, 23.73) = 4.498 | 0.0123 | Brown-Forsythe F(3, 21.88) = 8.019, p = 0.0009 | Significant group differences |
| 6B2 | Max Speed: Control/no-DREADD vs Pilo/no-DREADD | Dunnett’s T3 | Δ = −0.2593 m/s, t(16.13) = 3.395 | 0.0108 | 95% CI: [−0.4617, −0.0569] | Pilo mice had faster max speeds |
| 6B2 | Max Speed: Pilo/no-DREADD vs Pilo+DREADD | Dunnett’s T3 | Δ = +0.2371 m/s, t(19.59) = 2.939 | 0.0237 | 95% CI: [0.0278, 0.4464] | KORD reduced max speed in Pilo mice |
| 6B2 | Max Speed: Control/no-DREADD vs Control+DREADD | Dunnett’s T3 | Δ = −0.05129 m/s, t(14.78) = 2.023 | 0.1662 | 95% CI: [−0.1190, 0.0164] | No difference |
| 6C1 | Center Preference (% time): Across 4 groups | Kruskal–Wallis | H = 14.05, df = 3 | 0.0028 | Bartlett’s and BF tests p < 0.05 | Significant group differences |
| 6C1 | CP: Control/no-DREADD vs Pilo/no-DREADD | Dunn’s test | Mean rank diff = 16.86, Z = 2.713 | 0.02 | — | Pilo spent less time in center |
| 6C1 | CP: Pilo/no-DREADD vs Pilo+DREADD | Dunn’s test | Mean rank diff = −12.29, Z = 2.448 | 0.043 | — | KORD increased center preference in Pilo mice |
| 6C1 | CP: Control/no-DREADD vs Control+DREADD | Dunn’s test | Mean rank diff = −1.14, Z = 0.169 | > 0.9999 | — | No effect of KORD in controls |
| 6C2 | Center Speed: Across 4 groups | Kruskal–Wallis | H = 4.614, df = 3 | 0.2024 | Bartlett’s and BF tests p < 0.05 | No group difference in center speed |

**Table S7.** Statistical details associated with figure 6.

| **Figure** | **Test Description** | **Statistical Test** | **Statistic** | **p-value** | **Effect Size / Notes** | **Directional Summary** |
| --- | --- | --- | --- | --- | --- | --- |
| 7A1 | DI>0 across treatment, Control+DREADD | Logistic regression: Time + treatment; saline reference | n=36; pseudo-R2=0.1058; LL=-18.102; LLR p=0.3690 | 0.369 | 9 mice × 4 modeled sessions | Overall model not significant |
| 7A1 | Treatment order/time, Control+DREADD | Logistic regression | beta=0.1981; SE=0.4741; z=0.418; 95% CI [-0.731, 1.127] | 0.676 | OR=1.22 (95% CI 0.48-3.09) | No detectable order/time effect |
| 7A1 | CNO vs saline, Control+DREADD | Logistic regression; Wald contrast | beta=-1.5866; SE=1.3645; z=-1.163; 95% CI [-4.261, 1.088] | 0.2449 | OR=0.205 (95% CI 0.014-2.97) | No significant CNO effect |
| 7A1 | Sb vs saline, Control+DREADD | Logistic regression; Wald contrast | beta=-2.3400; SE=1.7058; z=-1.372; 95% CI [-5.683, 1.003] | 0.1701 | OR=0.096 (95% CI 0.003-2.73) | No significant Sb effect |
| 7A1 | Reversal vs saline, Control+DREADD | Logistic regression; Wald contrast | beta=-1.0019; SE=2.8055; z=-0.357; 95% CI [-6.500, 4.497] | 0.721 | OR=0.367 (95% CI 0.002-89.72) | No significant reversal difference |
| 7A1 | Total investigation: CNO vs saline, Control+DREADD | Paired t test | t(8)=1.22 | 0.257 | n=9 paired mice | No significant CNO-saline difference |
| 7A2 | Total investigation time, Control/no-DREADD | Repeated-measures ANOVA | F(2.29, 16.07)=0.719 | 0.52 | Greenhouse-Geisser-adjusted df | No detectable treatment effect |
| 7A2 | Continuous DI across treatments, Control/no-DREADD | One-way ANOVA | F(4,60)=1.0498 | 0.3893 |  | No detectable treatment effect |
| 7B1 | Total investigation, Pilo+DREADD | Mixed-model / repeated-measures ANOVA | F(2.829,29)=5.072 | 0.0068 | Omnibus effect; no corrected post hoc treatment-vs-saline difference | Total investigation varied by session |
| 7B1 | DI>0 across treatment, Pilo+DREADD | Logistic regression: Time + treatment; saline reference | n=80; pseudo-R2=0.08090; LL=-50.759; LLR p=0.1116 | 0.1116 | 16 mice x 5 sessions; CNO, Sb, Sb3, reversal distinct contrasts | Overall model not significant; prespecified CNO and Sb contrasts significant |
| 7B1 | Treatment order/time, Pilo+DREADD | Logistic regression | beta=-0.1169; SE=0.2053; z=-0.570; 95% CI [-0.519, 0.285] | 0.569 | OR=0.890 (95% CI 0.595-1.33) | No detectable order/time effect |
| 7B1 | CNO vs saline, Pilo+DREADD | Logistic regression; prespecified Wald contrast | beta=1.9745; SE=0.8085; z=2.442; 95% CI [0.390, 3.559] | 0.0146 | OR=7.20 (95% CI 1.48-35.13) | CNO associated with higher odds of DI>0 |
| 7B1 | Sb vs saline, Pilo+DREADD | Logistic regression; prespecified Wald contrast | beta=1.7802; SE=0.8478; z=2.100; 95% CI [0.119, 3.442] | 0.0357 | OR=5.93 (95% CI 1.13-31.25) | Sb associated with higher odds of DI>0 |
| 7B1 | Sb3 vs saline, Pilo+DREADD | Logistic regression; Wald contrast | beta=0.8560; SE=0.9329; z=0.918; 95% CI [-0.972, 2.684] | 0.3588 | OR=2.35 (95% CI 0.38-14.65) | No significant high-dose Sb3 effect |
| 7B1 | Reversal vs saline, Pilo+DREADD | Logistic regression; Wald contrast | beta=1.3073; SE=1.1852; z=1.103; 95% CI [-1.016, 3.630] | 0.27 | OR=3.70 (95% CI 0.36-37.72) | No significant reversal difference |
| 7B2 | Total investigation time, Pilo/no-DREADD | Repeated-measures ANOVA | F(1.59,23.94)=0.426 | 0.61 | Greenhouse-Geisser-adjusted df | No detectable treatment effect |
| 7B2 | Continuous DI across treatments, Pilo/no-DREADD | One-way ANOVA | F(4,75)=0.4633 | 0.7624 |  | No detectable treatment effect |
| 7B1 sens. | CNO vs saline, Pilo+DREADD: mouse-clustered SE sensitivity | Same Logit coefficients; SE clustered by MouseID | beta=1.9745; SE=0.8476; z=2.330; 95% CI [0.313, 3.636] | 0.0198 | OR=7.20 (95% CI 1.37-37.93) | Primary CNO inference retained |
| 7B1 sens. | Sb vs saline, Pilo+DREADD: mouse-clustered SE sensitivity | Same Logit coefficients; SE clustered by MouseID | beta=1.7802; SE=0.9185; z=1.938; 95% CI [-0.020, 3.580] | 0.0526 | OR=5.93 (95% CI 0.98-35.89) | Estimate unchanged; interval widens modestly |
| 7B1 sens. | Sb3 vs saline, Pilo+DREADD: mouse-clustered SE sensitivity | Same Logit coefficients; SE clustered by MouseID | beta=0.8560; SE=1.0448; z=0.819; 95% CI [-1.192, 2.904] | 0.4126 | OR=2.35 (95% CI 0.30-18.24) | No significant high-dose Sb3 effect |
| 7B1 sens. | Reversal vs saline, Pilo+DREADD: mouse-clustered SE sensitivity | Same Logit coefficients; SE clustered by MouseID | beta=1.3073; SE=1.0576; z=1.236; 95% CI [-0.766, 3.380] | 0.2164 | OR=3.70 (95% CI 0.47-29.37) | No significant reversal difference |
| 7D2 | Predicted optimal/retuned vs off-optimal DI, within mouse | Paired t test + bootstrap CI | mean delta=+17.43 points; t(24)=3.200; bootstrap 95% CI [6.80,27.55] | 0.00384 | n=25; Cohen dz=0.64; 21/25 mice higher in optimal/retuned state | DI higher in predicted optimal/retuned state |
| 7D3 | Linear predicted-position term | Quadratic OLS with SEs clustered by MouseID | beta=1.6436; SE=3.9760; z=0.413; 95% CI [-6.149,9.436] | 0.6793 | 75 state observations from 25 mice | No detectable linear position term |
| 7D3 | Quadratic predicted-position term | Quadratic OLS with SEs clustered by MouseID | beta2=-17.4315; SE=5.5223; z=-3.157; 95% CI [-28.255,-6.608] | 0.0016 | 75 state observations from 25 mice | Significant inverted-U-like curvature |
| S8A1 | Saline DI vs CP, Control/no-DREADD | Simple linear regression; observed complete cases | R2=.36 | 0.115 | Imputed DI excluded; unadjusted two-sided p value | No association |
| S8A1 | Saline DI vs CP, Pilo/no-DREADD | Simple linear regression; observed complete cases | R2=.00 | 0.863 | Imputed DI excluded; unadjusted two-sided p value | No association |
| S8A1 | Saline DI vs CP, Control+DREADD | Simple linear regression; observed complete cases | R2=.04 | 0.586 | Imputed DI excluded; unadjusted two-sided p value | No association |
| S8A1 | Saline DI vs CP, Pilo+DREADD | Simple linear regression; observed complete cases | R2=.04 | 0.498 | Imputed DI excluded; unadjusted two-sided p value | No association |
| S8A2 | Sb DI vs CP, Control/no-DREADD | Simple linear regression; observed complete cases | R2=.18 | 0.295 | Imputed DI excluded; unadjusted two-sided p value | No association |
| S8A2 | Sb DI vs CP, Pilo/no-DREADD | Simple linear regression; observed complete cases | R2=.00 | 0.88 | Imputed DI excluded; unadjusted two-sided p value | No association |
| S8A2 | Sb DI vs CP, Control+DREADD | Simple linear regression; observed complete cases | R2=.45 | 0.048 | Imputed DI excluded; unadjusted two-sided p value | Higher CP associated with lower DI |
| S8A2 | Sb DI vs CP, Pilo+DREADD | Simple linear regression; observed complete cases | R2=.37 | 0.02 | Imputed DI excluded; unadjusted two-sided p value | Higher CP associated with higher DI |
| S8A3 | CNO DI vs CP, Control/no-DREADD | Simple linear regression; observed complete cases | R2=.02 | 0.73 | Imputed DI excluded; unadjusted two-sided p value | No association |
| S8A3 | CNO DI vs CP, Pilo/no-DREADD | Simple linear regression; observed complete cases | R2=.13 | 0.17 | Imputed DI excluded; unadjusted two-sided p value | No association |
| S8A3 | CNO DI vs CP, Control+DREADD | Simple linear regression; observed complete cases | R2=.03 | 0.665 | Imputed DI excluded; unadjusted two-sided p value | No association |
| S8A3 | CNO DI vs CP, Pilo+DREADD | Simple linear regression; observed complete cases | R2=.14 | 0.195 | Imputed DI excluded; unadjusted two-sided p value | No association |
| S8A4 | Sb - saline DI change vs CP, Control+DREADD | Simple linear regression; observed complete cases | R2=.04 | 0.622 | Imputed DI excluded; unadjusted two-sided p value | No association |
| S8A4 | Sb - saline DI change vs CP, Pilo+DREADD | Simple linear regression; observed complete cases | R2=.35 | 0.026 | Imputed DI excluded; unadjusted two-sided p value | Higher CP associated with larger positive Sb-related DI change |
| S8A5 | CNO - saline DI change vs CP, Control+DREADD | Simple linear regression; observed complete cases | R2=.09 | 0.422 | Imputed DI excluded; unadjusted two-sided p value | No association |
| S8A5 | CNO - saline DI change vs CP, Pilo+DREADD | Simple linear regression; observed complete cases | R2=.25 | 0.072 | Imputed DI excluded; unadjusted two-sided p value | Positive association did not reach significance |
| S8B1 | Pilo+DREADD: CP threshold classifies Sb DI>0 | Exploratory within-sample ROC/Youden threshold | CP threshold=7.50; 14/14 correct | -- | n=14; accuracy 100%; sensitivity 100%; specificity 100%; TP=9 FP=0 TN=5 FN=0 | High-CP profile associated with DI>0 |
| S8B2 | Pilo+DREADD: CP threshold classifies positive Sb - saline DI change | Exploratory within-sample ROC/Youden threshold | CP threshold=8.88; 12/14 correct | -- | n=14; accuracy 85.7%; sensitivity 80%; specificity 100%; TP=8 FP=0 TN=4 FN=2 | High-CP profile associated with positive Sb-related change |
| S8B3 | Control+DREADD: CP threshold classifies Sb DI>0 | Exploratory within-sample ROC/Youden threshold | CP threshold=28.88; 8/9 correct | -- | n=9; accuracy 88.9%; sensitivity 100%; specificity 75%; TP=5 FP=1 TN=3 FN=0 | Low-CP profile associated with DI>0 |
| S8B4 | Control+DREADD: CP threshold classifies positive Sb - saline DI change | Exploratory within-sample ROC/Youden threshold | CP threshold=28.88; 6/9 correct | -- | n=9; accuracy 66.7%; sensitivity 100%; specificity 50%; TP=3 FP=3 TN=3 FN=0 | Low-CP profile associated with positive Sb-related change |
| S8B5 | Pilo+DREADD: CP threshold classifies CNO DI>0 | Exploratory within-sample ROC/Youden threshold | CP threshold=3.63; 12/14 correct | -- | n=14; accuracy 85.7%; sensitivity 100%; specificity 50%; TP=10 FP=2 TN=2 FN=0 | High-CP profile associated with DI>0 |
| S8B6 | Pilo+DREADD: CP threshold classifies positive CNO - saline DI change | Exploratory within-sample ROC/Youden threshold | CP threshold=9.88; 13/14 correct | -- | n=14; accuracy 92.9%; sensitivity 100%; specificity 87.5%; TP=6 FP=1 TN=7 FN=0 | High-CP profile associated with positive CNO-related change |

**Table S8.** Statistical details associated with figure 7 and supplementary figure 8.

| **Prompt (verbatim or paraphrased)** | **Purpose** | **Representative AI Response (summary)** | **How it was used** |
| --- | --- | --- | --- |
| “This sentence feels clunky—can you suggest a smoother way to phrase it while keeping the meaning the same?” | Text clarity | Suggested shorter, active-voice phrasing without changing meaning. | Used in revision of Introduction and figure legends. |
| “Can you check this paragraph for readability and reduce redundancy?” | Grammar and flow | Returned a streamlined version, eliminating repetition and simplifying sentence structure. | Incorporated edits into Results section. |
| “Can you help me add a short code block that saves collapsed results to a new .csv file?” | Coding clarification | Provided a small Python snippet showing how to export results with pandas. | Adapted and integrated into analysis pipeline. |
| “I’d like to add a section to my script that calculates effect sizes after running ANOVA. Can you show me how?” | Code addition | Supplied code to calculate Cohen’s d and partial eta squared. | Modified and used in final statistical scripts. |
| “Please reword this statistical reporting sentence so that it’s more concise but still clear to a general reader.” | Reporting clarity | Suggested language: “We fit a linear mixed-effects model (LMM) with treatment as a fixed effect and mouse as a random intercept.” | Adopted for consistency in Methods and figure legends. |

**Table S9.** Representative AI-assisted prompts and responses
